# Steric-Free Bioorthogonal Labeling of Acetylation Substrates Based on a Fluorine-Thiol Displacement Reaction (FTDR)

**DOI:** 10.1101/2020.09.09.290221

**Authors:** Zhigang Lyu, Yue Zhao, Zakey Yusuf Buuh, Nicole Gorman, Aaron R. Goldman, Md Shafiqul Islam, Hsin-Yao Tang, Rongsheng E. Wang

## Abstract

We have developed a novel bioorthogonal reaction that can selectively displace fluorine substitutions alpha to amide bonds. This fluorine-thiol displacement reaction (FTDR) allows for fluorinated cofactors or precursors to be utilized as chemical reporters; hijacking acetyltransferase mediated acetylation both in vitro and in live cells, which cannot be achieved with azide- or al- kyne- based chemical reporters. The fluoroacetamide labels can be further converted to biotin or fluorophore tags using FTDR, enabling the general detection and imaging of acetyl substrates. This strategy may lead to a steric-free labeling platform for substrate proteins, expanding our chemical toolbox for functional annotation of post-translational modifications (PTMs) in a systematic manner.

## INTRODUCTION

Bioorthogonal reactions have greatly facilitated protein labeling in complex biological systems and led to widespread applications including imaging, enrichment, and identification, etc.^1–2^ Representative bioorthogonal reactions such as azide-alkyne cycloaddition,^3^ Staudinger ligation,^4^ and tetrazine cycloaddition^2^ have resulted in the successful development of chemical reporters, which in conjunction with detection/affinity tags, have advanced our understanding of important biological pathways including PTMs.^5^ For instance, chemical reporters on acetylation have revealed many new protein substrates of p300,^6–7^ providing a more robust substrate recovery from proteome in comparison to anti-acetyl lysine antibodies.^8–9^ Yet, current chemical reporters are bulky in length and size, thereby impeding their general applications. Even alkyne or azide-based minimalist reporters for copper-catalyzed azide-alkyne cycloaddition (CuAAC) are still significantly larger than the inherent carbon-hydrogen bond (Figure 1A). This intrinsic steric hindrance has largely limited the application of chemical reporters to metabolic incorporation by enzymes possessing spacious active site pockets. For example, alkyne/azide-labelled PTM precursors or cofactors for acetylation^10–11^ and methylation^12–13^ were too bulky to be incorporated by many cognate transferases other than p300. While the “bump-hole” protein engineering strategy could work for a given enzyme, careful balances between mutations, structure folding, and function (potency, selectivity, etc) are required.^14^ For many acetyltransferases that function as subunits in protein complexes, mutations in the “hole” may alter their substrate specificities, leading to results different from in vivo sub-acylome,^6^ which undermines this approach’s broad applications.

**Figure 1.**
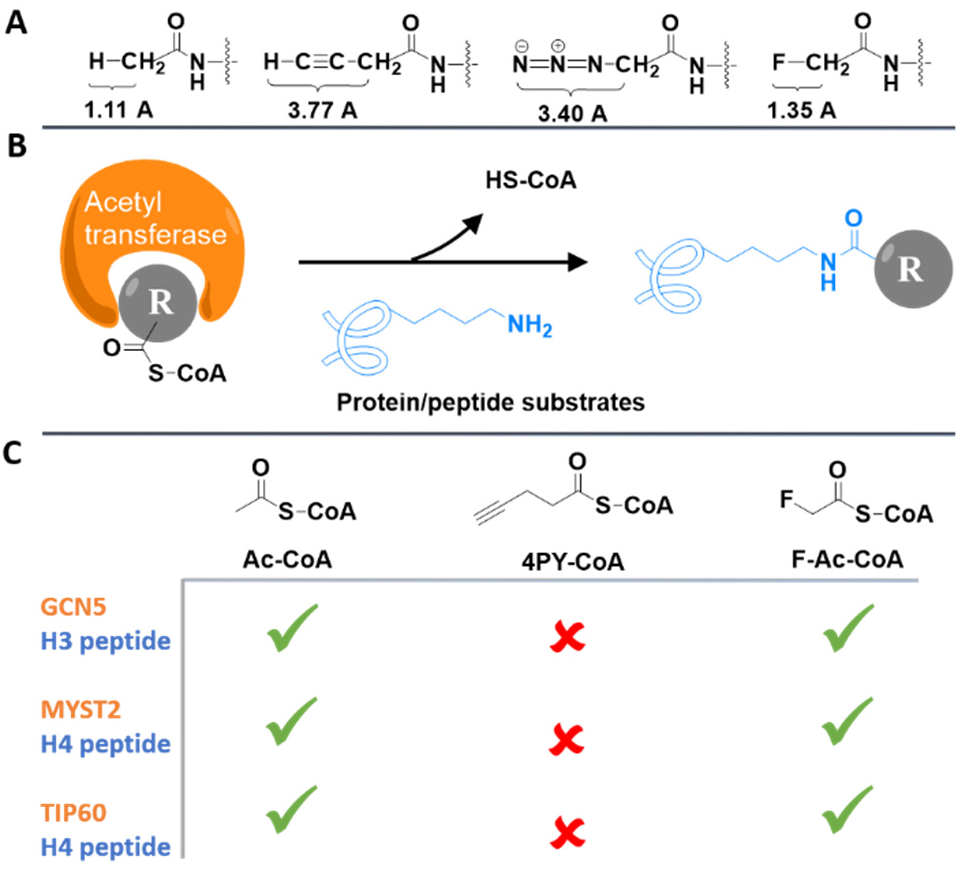
Acetyl-CoA analogs to label acetylation substrates through acetyltransferase assay. (A) Bond lengths of C-H, C-C≡CH, C-N_3_, and C-F on acetamide. (B) Illustration of the acetyltransferase assay. (C) Summary of the acetyltransferase assay results with Ac-CoA, 4-pentynoyl (4PY)-CoA, and F-Ac-CoA analogs, respectively.

Thus, global profiling of PTM substrates such as those of acetylation was still achieved by antibody-based detections.^15^ Yet, the enriched substrates and sites varied significantly per antibody.^15^ In our hands, a list of commercially available anti-lysine acetylation antibodies also revealed different sensitivity and substrate specificity (Figure S1). Taken together, elucidating the molecular targets of PTMs such as acetylation has been thereby compromised, despite being a key step towards the systematic dissection of PTMs and their roles in biological and pathology-related cellular signaling regulation.^11, 13, 16^ To this end, we asked whether a bioorthogonal reaction can be developed to generate reporters for the steric-free labeling of protein substrates and thereby allow for the global profiling of molecular targets.

## RESULTS AND DISCUSSION

The fluorine atom attracted our interest due to its orthogonality from biological molecules,^17^ similarity in size to hydrogen,^17–19^ and the close length of the carbon-fluorine bond to the carbon-hydrogen bond in acetamide, a representative acetylation functional group (Figure 1A).^19^ Fluorinated amino acids have been exploited to label proteins, bringing in minimal perturbations to protein structure and function.^17–18, 20^ Using acetylation for a proof of concept, we first wanted to investigate whether fluorinated acetyl-CoA can hijack the CoA metabolism, and be used by acetyltransferases to label their protein substrates (Figure 1B). Despite the in vivo toxicity of its pro-metabolite fluoroacetate,^21^ only the late-stage metabolite fluorocitrate was found accountable, mostly damaging organ tissues such as the kidney.^21^ Fluorinated acetyl-CoA and fluoroacetate haven’t yet been found to inhibit any enzyme, and have been successfully used to study metabolisms.^22–23^ Ac-CoA analogs with fluorine (F-Ac-CoA)^24^ or alkyne (4-pentynoyl (4PY)-CoA)^11^ functional groups were thereby synthesized, and each mixed with key acetyltransferase GCN5, MYST2, or TIP60 and their corresponding histone peptide substrates.^11, 25–26^ Mass spectrometric analysis (Figure S2) indicated successful acetyl or acetyl analog labelling after either acetyltransferase was incubated with wild type Ac-CoA or F-Ac-CoA. On the contrary, all acetyltransferases failed to incorporate the alkyne-modified 4-PY-CoA, which is consistent with previous reports.^6, 11^ Substitution with the electron-withdrawing fluorine also endowed a much higher hydrolysis rate to F-Ac-CoA, which somehow did not display significantly greater non-enzymatic reactivity (Figure S3).^27^ Taken together (Figure 1C), fluorine modification on the acetyl group afforded a relatively steric-free chemical reporter that can satisfactorily label substrates of acetyltransferases.

Next, we asked whether we can further modify the resulting fluoroacetamides with other tags such as a fluorescent dye “TAMRA”^28^ or a biotin affinity probe^29^ for detection, imaging, and future identification of protein substrates. Despite substantial progress in the development of fluorination methodology, there are few efforts on the replacement of fluorine. One relevant study^30^ has incorporated an alpha-fluorinated acetophenone moiety (**1**) into proteins and observed its reaction with a proximal cysteine. This makes us hypothesize that fluorinated acetyl groups may react with thiol derivatives, even at the small molecule level. To test this, we reacted a representative thiol compound, benzenethiol, with a few fluorinated acetyl substrates in water (Scheme 1). In the presence of a strong base, the primary fluorine alpha to acetophenone (**1**), ketone (**2**), and amide (**3**) were all efficiently displaced (>90% yields). We also observed a similar conversion with the fluoroacetate derivative (**4**) despite a lower yield, which is likely caused by hydrolysis of the ester, given the isolated benzeneethanol side product. The result with **3** particularly encouraged us as it is a model fluoroacetamide substrate for fluorinated peptides/proteins.

**Scheme 1.**
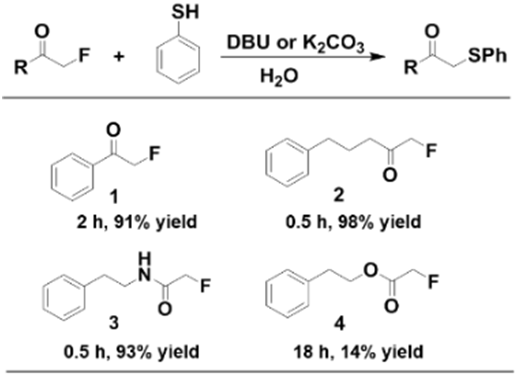
Exploration of Fluorinated Substrate

To identify the optimal pH range for this reaction, we titrated the pH (Figure S4) and found that the reaction rate significantly increased with pH, likely due to the increased deprotonation of benzenethiol. Using mildly basic condition, pH 8.5, in which the reaction has the best reaction rate, we also examined substrate **3**’s reactivity with glutathione and cysteine, both as strong intrinsic nucleophiles that exist in cellular environments (Figures S5-S6). Surprisingly, no reaction was observed for both cases upon incubation for 24h, suggesting that substrate **3** could be bioorthogonal to other nucleophilic species. We then investigated substitution effects on benzenethiol in order to efficiently convert fluoroacetamide under the mild physiological reaction condition.^31–32^ Exploration of a series of benzenethiol derivatives (**5**-**13**) found that electron donating groups at the ortho- and para-positions facilitate the reaction by increasing the nucleophilicity (Figure 2, S7). Given their theoretical pKa’s (< 7.0) (Figure S8), almost all of these derivatives should be fully deprotonated. The most reactive derivative, **13**, is a trimethoxy analog that can reach ~ 81% reaction conversion within 13 hrs. Despite potential issues regarding sterics for the ortho-substitutions, the superior reactivity of **13** in comparison to the meta-substituted derivative **12** indicated that the methoxy groups are not bulky enough to perturb reactivity. Likewise, derivative **9** that only bears two substitutions at the electron donating sites ended up possessing the same reactivity as **12**. Follow-up kinetics studies based on the reported procedures^33^ suggested that the bimolecular reaction between **3** and **13** followed second order kinetics, and had an observed rate constant of (1.03 ± 0.06) × 10^−3^ M^−1^ S^−1^ (Figure S9), similar in rates to the classic Staudinger ligation reaction.^33–34^

**Figure 2.**
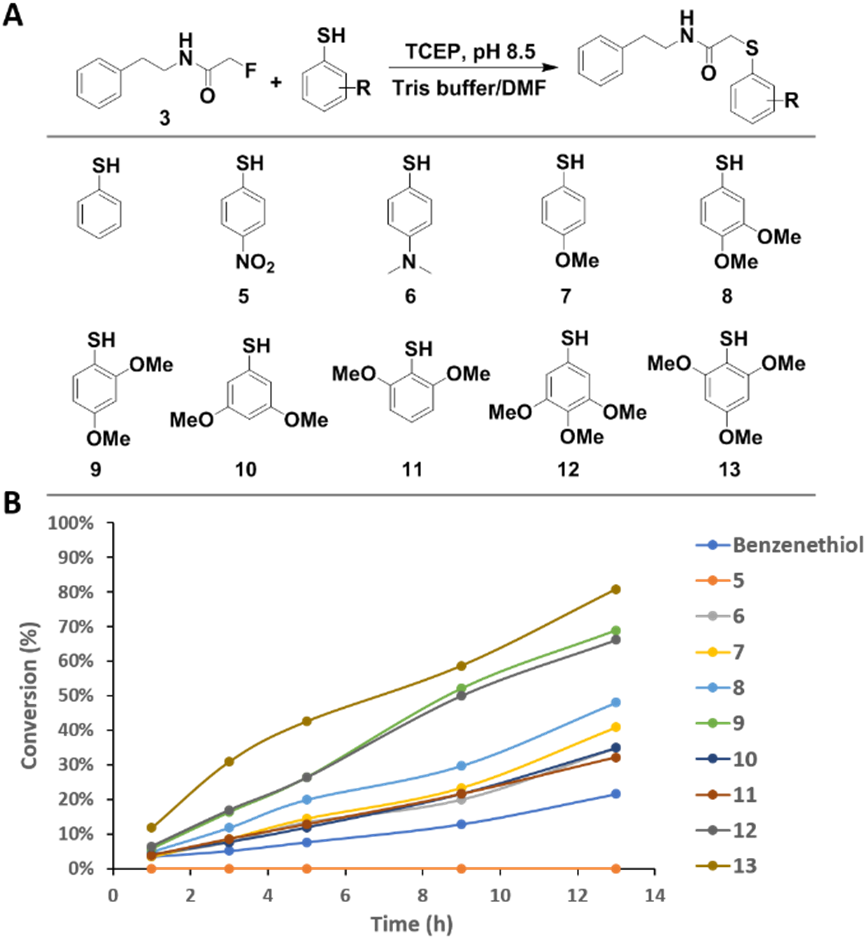
Structure-activity relationship study of benzenethiol derivatives. (A) Scheme of the benzenethiol derivatives. (B) Plot of the conversion against reaction time.

Moreover, the stability of **13** along with substrate **3** were further evaluated in cell lysates (Figure S10), and most of them remained intact (61.8% and 100%, respectively). On the contrary, control **3-Cl** possessing the reported chloroacetamide^35^ had only 0.15% left, consistent with previous observations that chloroacetamide readily reacted with cellular proteins.^36^ With the aid of LC-MS/MS, the FTDR product of **3** and **13** in cell lysates was also confirmed (Figure S11), further corroborating the bioorthogonality of FTDR. Notably, wild-type Ac-CoA has been regarded as a central metabolite that can be also used for ketogenesis, mevalonate pathway, and fatty acid synthesis.^37^ F-Ac-CoA as a mimic of Ac-CoA was demonstrated to be taken by fatty acid synthetases to make fluorofatty acids,^38^ most of which are also classic PTMs. We thereby tested the FTDR reaction between probe **13** and the alpha-fluorinated model substrates of fatty acids such as butyrate, myristic acid, palmitic acid, malonic acid, and succinic acid, etc. Surprisingly, no FTDR reaction was observed with these substrates, suggesting that due to the increased steric hindrance the secondary fluorides in these fatty acids may not be readily displaceable as the primary fluoride in fluoroacetamide. On the other hand, these observations also indicated the potential uniqueness of fluoroacetamide and its orthogonality to other fluorine-substituted natural molecules for FTDR-based detection, imaging, and target identification.

Hence, with the most active benzenethiol derivative in hand, we returned to the intriguing question of converting fluoroacetamide to functional detection tags such as fluorescent dyes and biotin probes,^29, 39^ Thus, we started by constructing a biotin probe (**14**, Biotin-SH) that contains the benzenethiol structure of **13** as the warhead, and the glutamic acid building block (**29**) as the connecting unit to improve overall solubility (supplementary schemes 7-9). Due to the convenience of ESI-MS characterization on peptides, we first demonstrated the successful conversion of the fluorine label to a biotin tag on the aforementioned histone H3-20 peptide substrate (Figure S12). Next, we examined the whole process of labelling and tagging (Figure 3A) on histone protein substrates. Histone H3.1 (Figure 3B) and H4 subunit (Figure S13) were each incubated with F-Ac-CoA, and the corresponding KAT under standard in vitro enzymatic reaction conditions.^11^ The mixtures were then incubated with the Biotin-SH probe for varied time periods, with subsequent in-gel fluorescent imaging to detect the biotin tagging. For both histone proteins, efficient biotinylation occurred within 1h of incubation, and gradually reached saturation after 3h. Staining with coomassie brilliant blue (CBB) also revealed a complete change of molecular weight, presumably due to the two-step modification. The little signal observed from the control group without prior fluorination also indicated the relative specificity of Biotin-SH. Concurrently, failure of biotinylation was observed for the group using CuAAC labeling, which was consistent with literature reports, and confirmed that many KATs are incapable of up-taking sterically hindered substrates.^6, 11^ To check if the fluoroacetyl label could be recognized by histone deacetylases, we also incubated these fluoroacetylated histone proteins with a few reported deacetylases^40^ before Biotin-SH tagging. Interestingly, most fluoroacetylation can be efficiently removed (Figure S14), suggesting that the label could be substrates for deacetylases as well.

**Figure 3.**
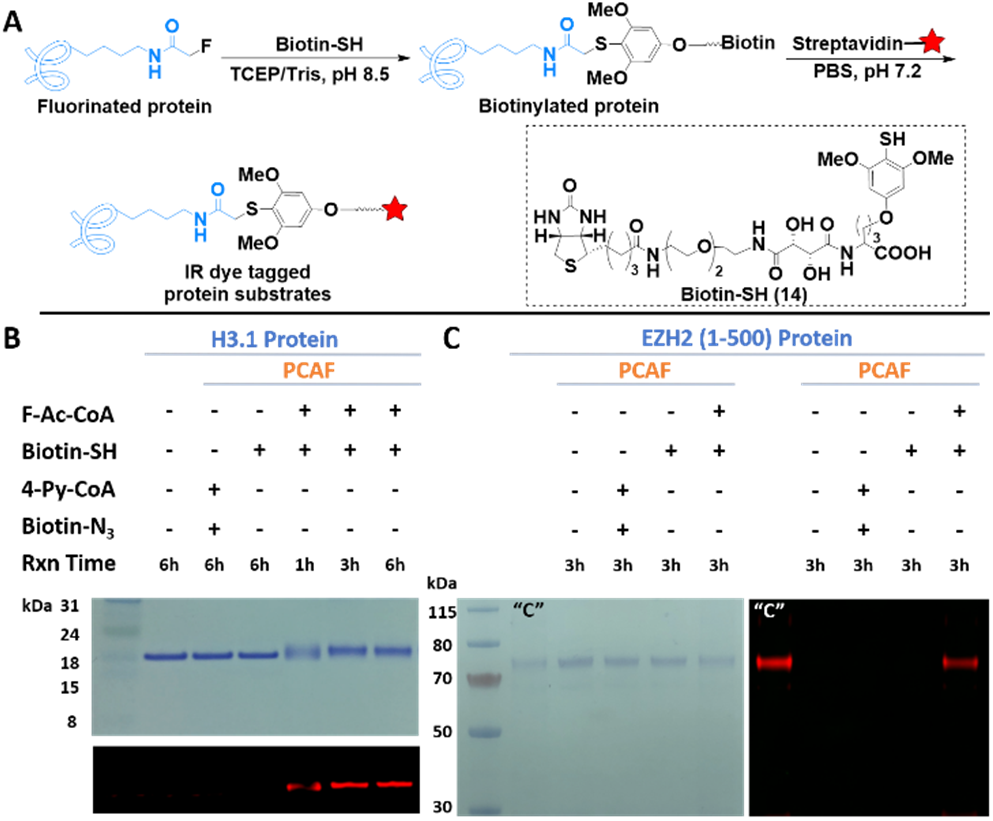
FTDR-based tagging of protein substrates with Biotin-SH probe. (A) Reaction scheme; The red star indicates IR dye. (B-C) Labeling of histone protein H3.1, and nonhistone substrate EZH2 (1-500), respectively; The top panel for (B) and the left panel for (C) are gel images after CBB staining. The other images are for in-gel fluorescent detection of IR dye. “C” is the positive control for CuAAC, which has been prepared by NHS ester labeling of the lysines on EZH2 with the ‘click chemistry’ tag, followed by CuAAC reaction with the corresponding biotin linker.

In addition to histones, a growing number of non-histone substrates were recently uncovered, with their roles in protein function regulation being increasingly appreciated.^16^ For instance, enhancer of zeste homolog 2 (EZH2) was revealed to be acetylated by PCAF mainly at lysine 348, which improved EZH2’s stability and promoted the migration of lung cancer cells.^41^ To evaluate whether our labeling strategy is applicable to non-histone substrates, we specifically constructed and expressed the truncated form of EZH2 (1-500) in E. coli, with a shake-flask yield of approximately 2 mg/L. The protein’s purity and size were confirmed by SDS-PAGE (Figure S15). Reaction of the EZH2 fragment under the same conditions as those for histone proteins resulted in a similar type of tagging as evident by Figure 3C. Taken together, these observations displayed the steric-free in vitro labeling and tagging of a range of known protein substrates using the FTDR reaction, which could not be achieved by the known CuAAC method.

For labeling proteins in living cells, azide or alkyne analogs of fatty acids have been exploited, which were metabolized intracellularly into CoA derivatives.^3–5, 10^ To increase their cellular delivery, pro-metabolites with esters masking the polar carboxylate group constituted an effective strategy in recent years.^42^ Nevertheless, the intrinsic sterics of these pro-metabolites resulted in varied and suboptimal labeling results,^10, 42^ sometimes requiring extensive structural optimization.^42^ To explore the utility of our probing system for studying acetylation in the cellular level, we designed the fluorinated version of pro-metabolite, ethyl fluoroacetate (Figure 4A). Given the in vivo toxicity of fluoroacetate,^21^ we first evaluated the cell cytotoxicity, and found that this pro-metabolite exhibited minimal toxicity with doses up to 2 mM after 12h of incubation (Figure S16). Additional LC-MS/MS studies indeed confirmed its conversion by enzymes to fluoroacetyl-CoA in live cells (Figure S17), which, taken together with the observed minimal toxicity in cell lines, could support the applicability of ethyl fluoroacetate and its CoA metabolite to studies in the cell level.

**Figure 4.**
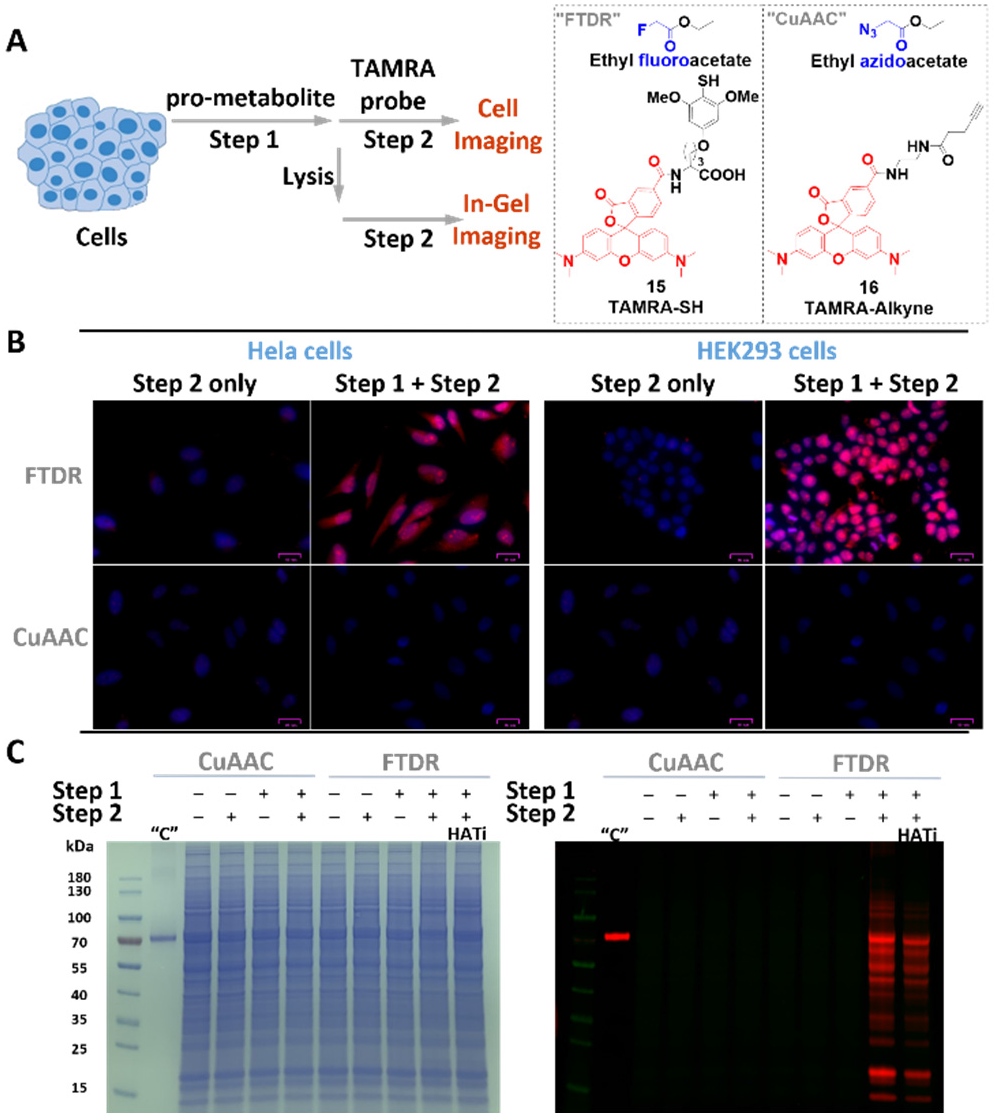
Cellular evaluation of FTDR-based tagging with TAMRA-SH probe. (A) Scheme for cellular pro-metabolite incorporation (1 mM, 6h, at 37 °C, step 1) and protein substrate detection (step 2). (B) Fluorescent microscopy of fixed and permeabilized cells that were stained by Hoechst 33342 (blue) and TAMRA probes (red); Scale bars: 25 μm. (C) Cell lysate protein labeling by pro-metabolites and detection by TAMRA probes (red); Left panel: PAGE gel stained by CBB; Right panel: In-gel fluorescent detection. “C” is the positive control for CuAAC, which randomly labelled lysines on BSA with azide-NHS ester, followed by CuAAC mediated conjugation with TAMRA-Alkyne. “HATi” indicates the addition of HAT inhibitors (anacardic acid/MG149) prior to step 1.

With confidence in the safety profile of our pro-metabolite, we started by treating it with two representative cell lines, HeLa, and HEK293, followed by subsequent FTDR with the TAMRA-SH probe (**15**) as the second step for fluorescent detection (Figure 4A). Treatment with the azido modified pro-metabolite^42^ and the TAMRA-alkyne (**16**) for CuAAC chemistry was performed in parallel as a control, wherein weak signals were previously reported.^42^ Direct microscope imaging studies (Figure 4B) revealed much stronger intracellular labeling and tagging with TAMRA following the FTDR based approach, suggesting a drastically more complete profiling of acetylation substrates. We also observed significant fluorescence not only in the nucleus but also cytoplasm, which may indicate the successful labeling of both histones and non-histone proteins. The little background signals emitted from the cells treated with only **15** (step 2) further demonstrated the specificity of the developed-SH probes. Following similar procedures, we also tested the labeling of the cell lysates after metabolic incorporation of fluorine reporters. Lysates following FTDR or CuAAC mediated ligation with TAMRA were separated on SDS-PAGE, and visualized by CBB to confirm the equal amount of protein loading (Figure 4C). Yet, multiple labelled protein bands spanning a wide range of molecular weights were observed only for the lysates of cells that underwent a complete two-step process of FTDR (Figure 4C). This discovery was consistent with the microscope imaging results. Further, pretreatment of the cells with anacardic acid and MG149 (p300, pCAF, Tip60 and MOF inhibitor)^43–45^ to inhibit certain histone acetyltransferases before the two-step process of FTDR also resulted in the general weakening of the labeling intensity. In addition, we exploited the concurrent incubation with HDAC inhibitor cocktails and observed slightly decreased labeling (Figure S18), which was consistent to literature reports,^9, 42, 46^ suggesting that incorporation of F-acetylation needs prior deacetylation of intrinsically acetylated lysine residues in order to make the lysine available, and blocking the removal of wild-type acetylation prevents metabolic incorporation of F-acetylation.^9, 42, 46^ Taken together, these observations fully supported that FTDR allowed for profiling the proteome-wide substrates of acetyltransferases from the cellular contexts.

To further validate that the FTDR-based two-step metabolic labeling occurs on acetylation protein substrates, we then performed histone extraction (Figure 5A) and confirmed the existence of TAMRA labeling on the primary acetylation substrates, histones (Figure 5B). As controls, both the treatment with HAT inhibitors and the competition with acetate have resulted in decreased TAMRA-SH probe labeling, suggesting that the FTDR-based two-step labeling is acetyltransferase-dependent, and relies on F-Ac-CoA metabolites. To test if the FTDR-based labeling can be used to enrich these protein substrates, we treated cell lysates with the Biotin-SH probe in step 2, and pulled down the labelled proteins (Figure 5A). Western blot analysis confirmed the presence of known acetylation substrates including histones, and alpha-tubulin^47–48^ only in the protein pool enriched from the cell lysates that have been subjected to the two-step process (Figure 5C). Accordingly, pretreatment with HAT inhibitors decreased the amount of proteins enriched, in particular for H3 and H4. The level of alpha-tubulin was not weakened as much, presumably due to the fact that certain acetylation sites such as lysine 40 on alpha-tubulin is mediated by other acetyltransferases (e.g. αTAT1)^47^ that were not targeted by the administered HAT inhibitors.

**Figure 5.**
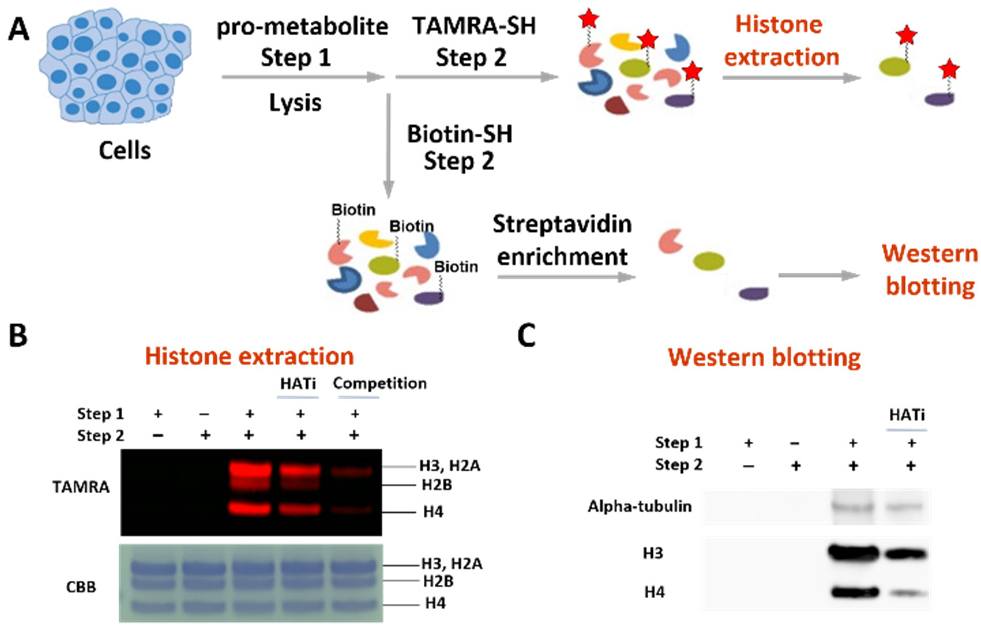
Validation of the FTDR-based labeling of acetylation substrates. (A) Scheme for cellular pro-metabolite incorporation (step 1), protein substrate labeling by TAMRA-SH (step 2), and extraction of the known acetylation substrates histones; or protein substrate labeling by Biotin-SH probe (step 2), enrichment with streptavidin beads, followed by western blot analysis of the proteins pulled down to examine the existence of alpha-tubulin, histone H3 and H4. (B) The histone extraction results. Top panel: In-gel fluorescent detection; Bottom panel: CBB staining. (C) The western blotting results.

Lastly, to gain insight of the specific labeling sites by our tagging strategy, we extracted out histones and pulled down alpha-tubulin from the cells having incorporated the pro-metabolite ethyl fluoroacetate, and carried out proteomics studies (Figure 6A). As shown in Figure 6B and Figures S19-S21, fluoroacetylation has been observed on lysine 30, lysine 34, lysine 43, and lysine 57 of histone H2B; lysine 5, lysine 8, lysine 12, lysine 16, lysine 31, lysine 44, lysine 59, lysine 77, lysine 79, and lysine 91 of histone H4; lysine 40, lysine 60, lysine 163, lysine 164, lysine 166, and lysine 280 of alpha-tubulin. Almost all of these sites have been consistent with the previous literature reports on acetylation sites,^48–51^ further demonstrating that this labeling strategy can probe lysine acetylation in a general manner. Notably, N-terminal acetylation recently draws significant research interest, which is catalyzed by a specific class of N-terminal acetyltransferases using Ac-CoA, and has been revealed to play important roles in protein functions and cellular locations, etc.^52–53^ During the preliminary proteomics studies of histone H2B and H4 (Figure S19-S20), we haven’t been able to observe any F-acetylation at the N-terminus. Although N-terminal acetylation is widespread and common in human proteins, it may only exist partially within the pool of molecules of a given protein substrate,^54^ and there are many internal lysine acetylation sites for each N-terminal protein.^52^ More importantly, N-terminal acetylation is considered irreversible,^52^ which could block the incorporation of F-acetylation to the intrinsic N-terminus. Thus, our FTDR-based labeling and imaging could be actually specific to internal lysine sites.

**Figure 6.**
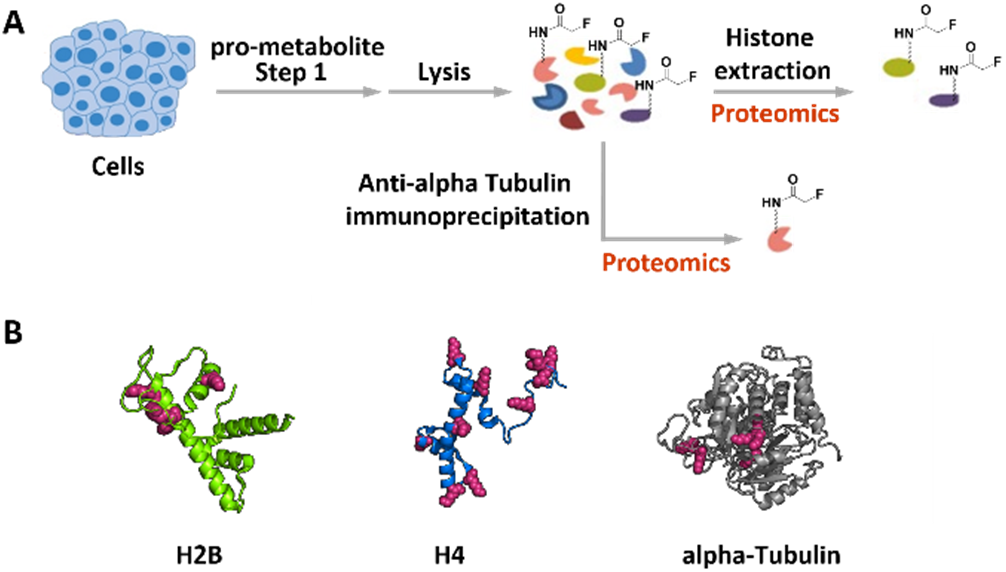
Validation of the fluoroacetyl labeling sites on known protein substrates by proteomics analysis. (A) Scheme for cellular pro-metabolite incorporation, lysis, and the histone proteins extraction; or immunoprecipitation of alpha-tubulin after cellular pro-metabolite incorporation. (B) Summary of the F-Ac labeling sites (red) on representative histone proteins H2B, H4 (PDB: 1kx5), and alpha-tubulin (PDB: 1tub). Details of the proteomics results are available in the supporting information (Figures S19-S21)

## CONCLUSION

In summary, we have developed a fluorine-thiol displacement reaction (FTDR), and used it for steric-free labeling of protein substrates of a representative PTM, acetylation. Along with the benzenethiol derived functional tags, the FTDR-based imaging and detection of substrates demonstrated great potential for globally profiling acetyl transferases’ substrates, which are of vital significance for understanding the roles of acetylation in physiology and disease. Although the fluorine tag possesses less steric than the alkyne/azide tags commonly used for CuAAC chemistry, our FTDR rate appeared to be slower and requires mildly basic pH, which could limit certain biological applications such as real-time imaging. Thus, future work would focus on improving and optimizing the FTDR reaction conditions. Nevertheless, this tool kit, together with future applications to quantitative proteomics studies, is expected to offer versatile probes for identifying targets of acetylation, and possibly many other PTMs that are mediated by transferases with restricted active sites.

## Supporting Information

Materials and methods, supplementary figures, and NMR spectra are included in the supporting information, and is available free of charge via the internet at http://pubs.acs.org.

## Notes

The authors declare no competing financial interests.

## ACKNOWLEDGMENT

This work was supported by funding from the National Cancer Institute of the National Institute of Health (NIH) under Award Number P30 CA006927, R50 CA221838, and National Institute of General Medical Sciences of the NIH under Award 1R35GM133468-01. Support for the Wistar Proteomics and Metabolomics Facility was provided by P30 CA010815 and S10 OD023586. Support for the NMR facility at Temple University by a CURE grant from the Pennsylvania Department of Health is gratefully acknowledged.

---

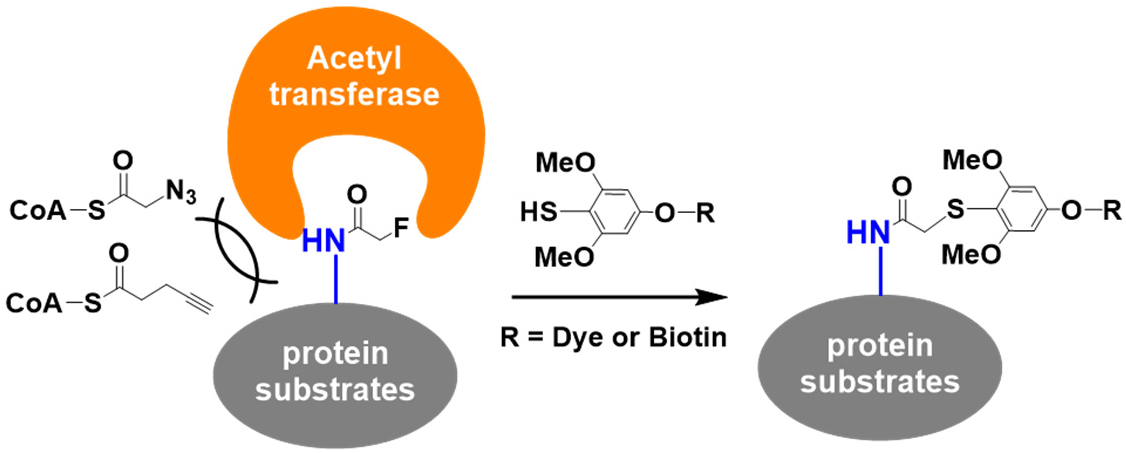

## Notes

### Competing Interest Statement

The authors have declared no competing interest.

