## Supporting for "Steric-Free Bioorthogonal Labeling of Acetylation Substrates Based on a Fluorine-Thiol Displacement Reaction (FTDR)"

#### Table of Contents

Materials and Methods

Chemical Synthesis…………………………………………………… S2

Biological Experiments……………………………………………… S22

Supplemental Figures………………………………………………….. S31

$^{1}$H-, $^{13}$C-, and $^{19}$F NMR Spectra for New Compounds…………………………. S52

References……………………………………………………………… S69

### Materials and Methods:

#### *Chemical Synthesis*

**General Information:**

Chemical reagents and solvents were purchased from commercial resources such as VWR, Thermo Fisher, and Sigma Aldrich, and were used directly without further purification. Analytical TLC was carried out with Silica Gel 60 $F_{254}$ plates (EMD Chemicals). The chemicals on TLC were either visualized by UV 254 nm (UV lamp, Chemglass Life Sciences) or stained by phosphomolybdic acid or $KMnO_4$ oxidation. Compound purification was performed by normal-phase flash column chromatography on columns manually loaded with silica gel grade 60 (230-400 mesh, Fisher Scientific) or by reverse-phase Combi-Flash on prepacked C18 columns (Teledyne ISCO). Further purification by preparative high-performance liquid chromatography (HPLC) was implemented on Waters 1525 series that consist of a 2489 UV/vis detector, 1525 binary pump, and an XBridge Prep C18 column. Routine mass spectrometry analysis was done using liquid chromatography-mass spectrometry (LC-MS) Agilent 1100 series. High resolution LC-MS analysis was performed on an Agilent 6520 Accurate-Mass Quadrupole-Time-of-Flight (Q-TOF) coupled with an electrospray ionization source. For NMR analysis, $^{1}H$ NMR and $^{13}C$ NMR spectra were recorded on 400 MHz or 500 MHz Bruker Advance. The raw data were processed with MestReNova, and the chemical shifts were reported in parts per million (ppm) downfield from the internal standard tetramethylsilane (TMS).

**Supplementary scheme 1** – synthetic scheme of 2-fluoro-1-phenylethan-1-one.

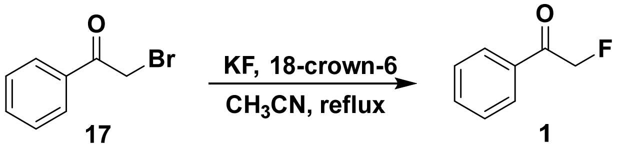


**Supplementary scheme 2** – synthetic scheme of 1-fluoro-5-phenylpentan-2-one.

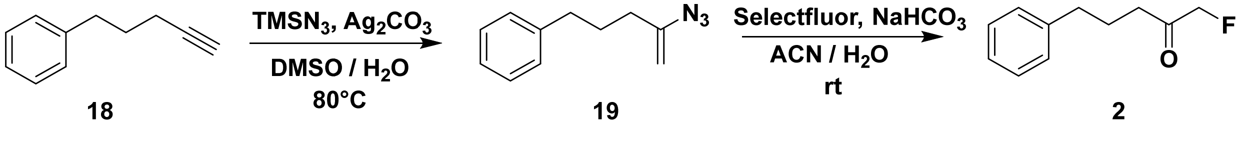


**Supplementary scheme 3** – synthetic scheme of 2-fluoro-N-phenethylacetamide.

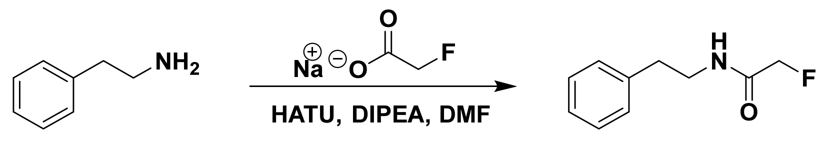


20 3

**Supplementary scheme 4** – synthetic scheme of 2-fluoro-phenethylacetate.

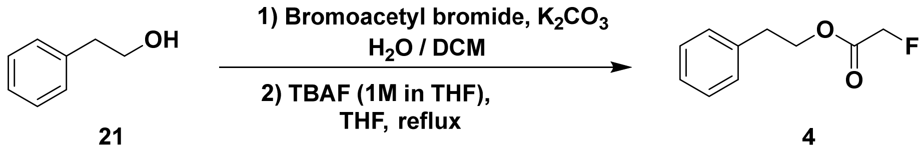


21 4

**Supplementary scheme 5** – synthetic scheme of 3,4,5-trimethoxybenzenethiol.

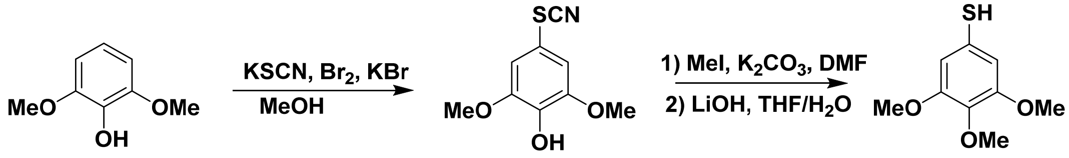


22 23 12

**Supplementary scheme 6** – synthetic scheme of 2,4,6-trimethoxybenzenethiol.

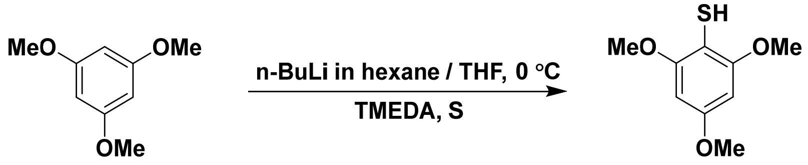


24 13

**Supplementary scheme 7** – synthetic scheme of N,N-bis[(1,1-dimethylethoxy)carbonyl]-5-iodo-phenylmethyl ester.

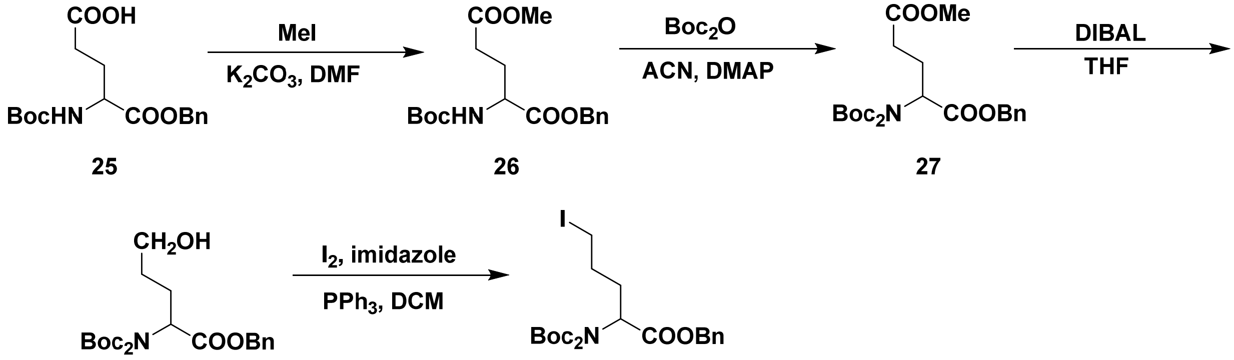


25 26 27

28 29

**Supplementary scheme 8** – synthetic scheme of Benzyl 2-amino-5-(3,5-dimethoxy-4-thiocyanatophenoxy) pentanoate.

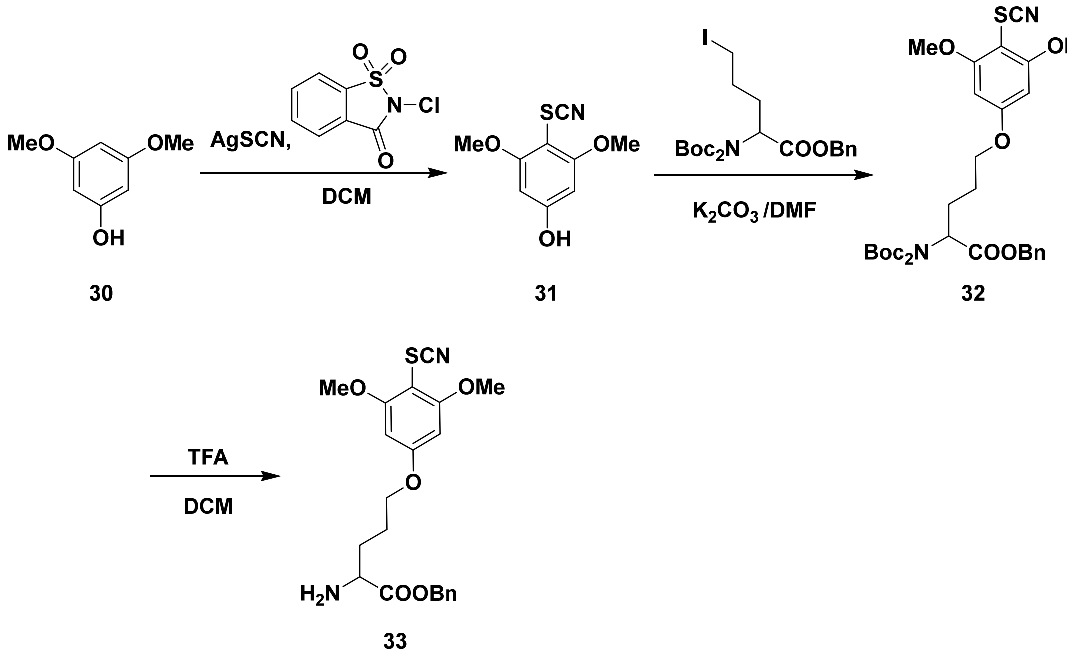


**Supplementary scheme 9** – synthetic scheme of the diol cleavable biotin linker conjugated with the 4-mercapto-3,5-dimethoxyphenoxy probe (Biotin-SH).

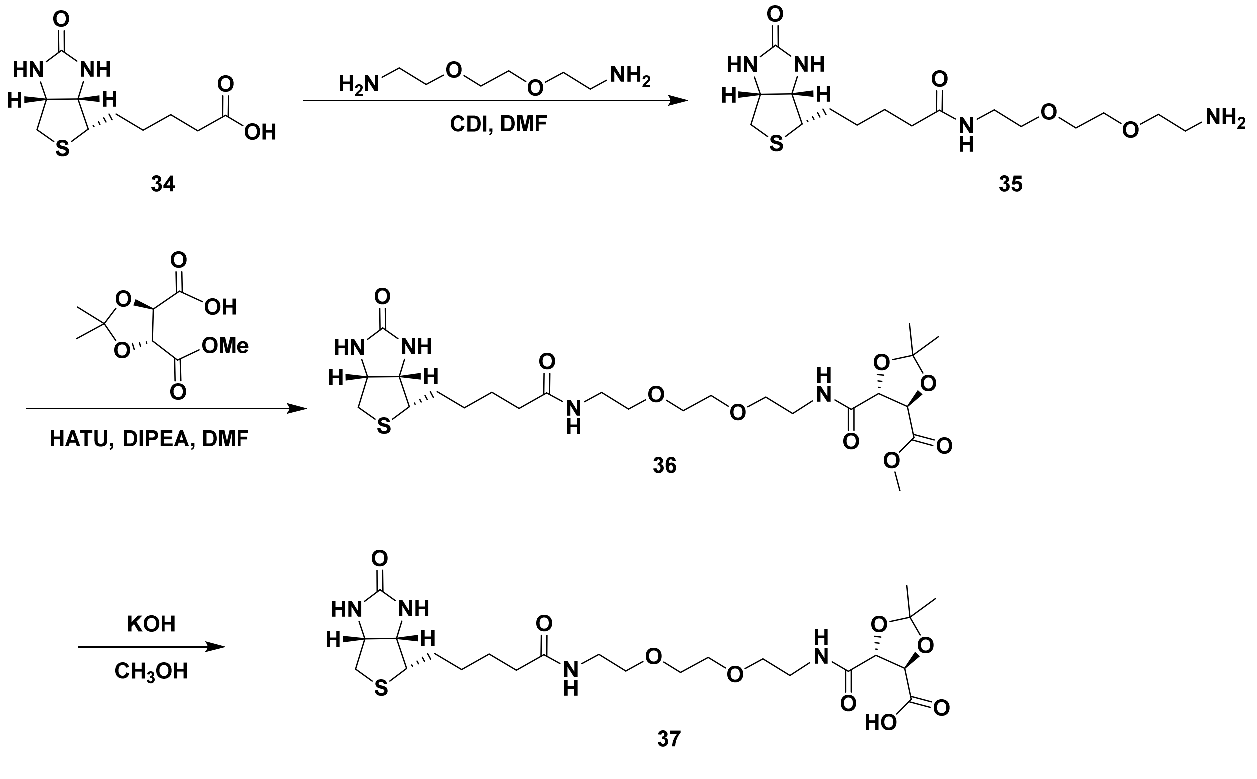

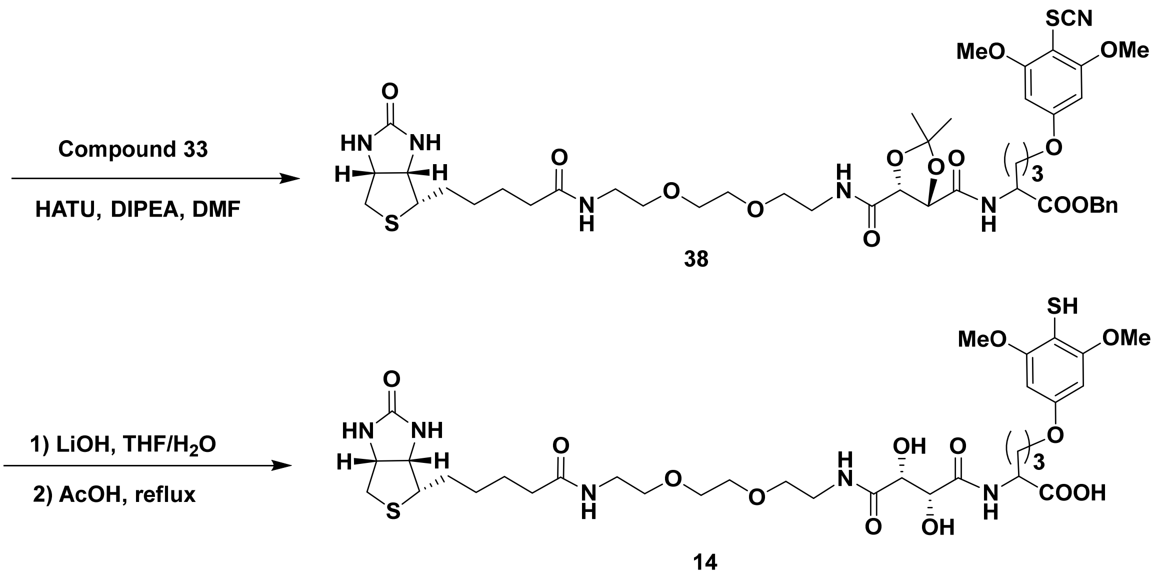


**Supplementary scheme 10** – synthetic scheme of the TAMRA dye conjugated 4-mercapto-3,5-dimethoxyphenoxy probe (TAMRA-SH).

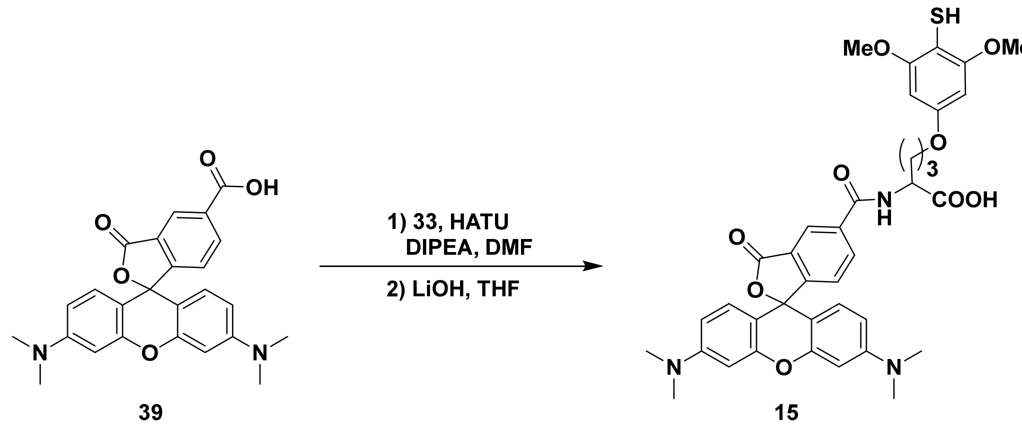


**Supplementary scheme 11** – synthetic scheme of the TAMRA dye conjugated alkyne probe (TAMRA-Alkyne).

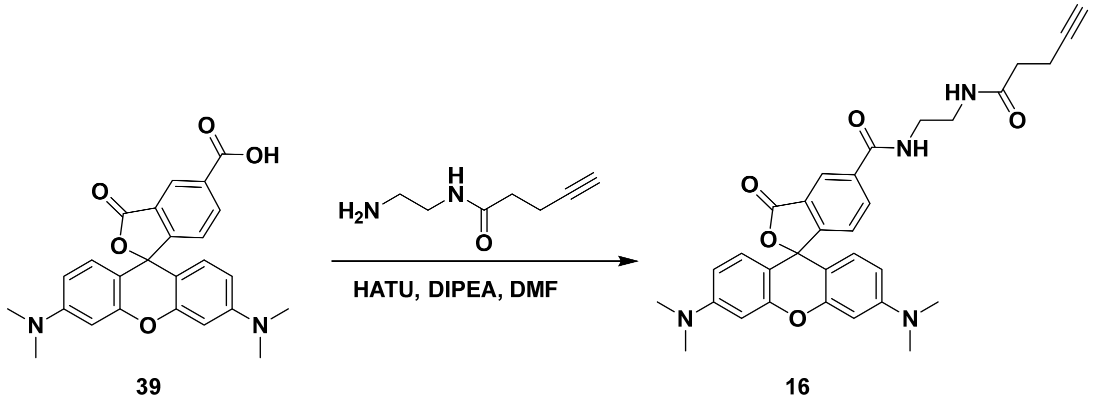

**General Procedure for Exploration of Fluorinated Substrate:**

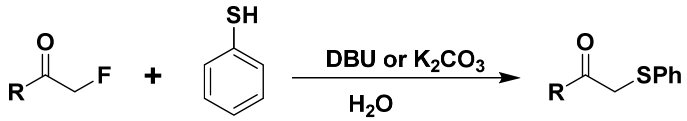


**General Procedure A:** The nucleophile thiophenol (0.4 mmol, 44 mg) was mixed with 0.2 mmol α-fluorocarbonyl derivative (compounds **1** - **4**) in 1 mL water. Then 0.6 mmol DBU (91 mg) or 0.4 mmol potassium carbonate (55 mg) was added to the reaction mixture. After stirring at room temperature with indicated time, 5 mL ethyl acetate was added to the flask to quench the reaction. The organic layer was separated, dried with anhydrous sodium sulfate, vacuum concentrated, and subsequently purified via flash column chromatography to provide the desired thiophenol adduct (compounds **42** - **45**).

**Optimizing Reaction Conditions - pH Titration:**

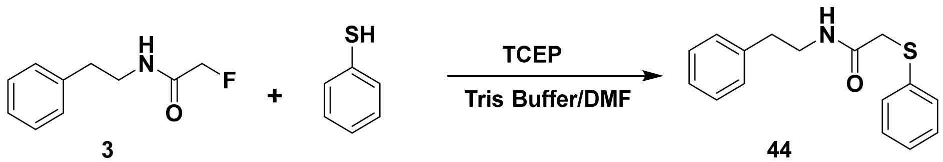


Substrate **3**, 2-Fluoro-N-phenethylacetamide, was dissolved in DMF to make a 1 M stock solution. About 0.75 µL of it was mixed with 5 µL of thiophenol stock (300 mM in DMF), and 2 µL of TCEP solution (1.5 M stock in water, pH 7.4). DMF/Tris buffer were added to make a total volume of 30 µL (50% of Tris buffer), during which 2M HCl or NaOH was slightly added to adjust the final pH value to 6.5, 7.5, or 8.5. Thus, the final concentration of substrate **3**, thiophenol, and TCEP was 25 mM, 50 mM, and 100 mM, respectively. The mixture was reacted at 37 ºC. Approximately 3 µL of the reaction mixture was taken out at indicated time points (12 h) and was mixed with 30 µL 0.5% TFA/ACN to quench the reaction. The samples were analyzed by LC-MS. To reduce the inter-assay variations and errors, the yield of product **44** was determined by comparing the UV peak area ratio of the product / (the product + unreacted substrate) in each LC-MS assay with the standard curve. The standard curve was plotted by the known concentration ratios of "[the product]/([the product] + [substrate])" against the corresponding UV peak area ratios. The denominator of the equation equals the very initial concentration of the substrate (25 mM for the current reaction).

#### Bioorthogonality of the Fluorine-Thiol Displacement Reaction

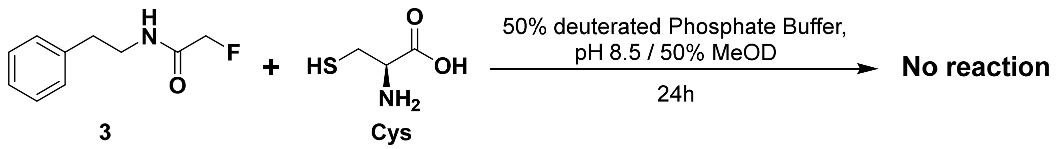


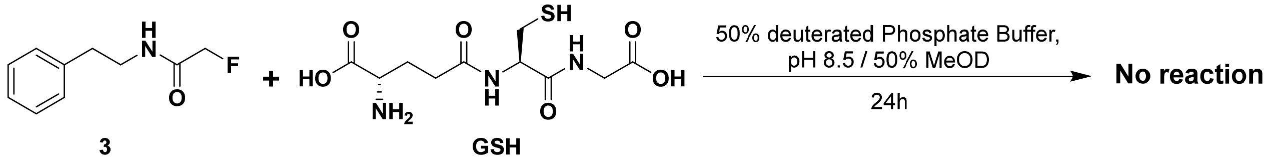


The common substrate 2-fluoro-N-phenethylacetamide (compound **3**) was dissolved in MeOD as a 10x stock solution (250 mM). Sixty microliter of the stock was mixed with another 60 µL of either the reduced glutathione or cysteine 10x stock solution (250 mM in $D_2O$). The mixture was added with additional deuterated solvents (1:1 mix of deuterated sodium phosphate buffer and MeOD) to make a final volume of 600 µL (pH adjusted to 8.5). The resulting solution was incubated at 37 °C water bath for 24 h, and then analyzed by $^1$H NMR spectroscopy.

#### General Procedure for Structure-Activity Relationship Studies of Nucleophiles:

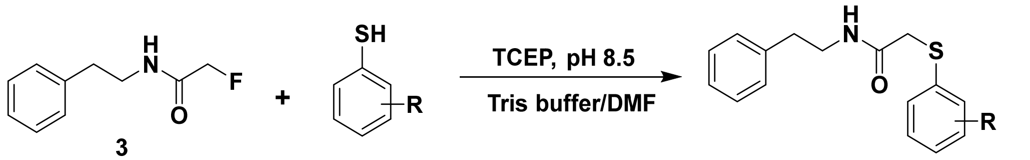


**General Procedure B:** 2-Fluoro-N-phenethylacetamide (**3**) (25 mM, 5 µL 0.5 M stock in DMF) and substituted benzenethiol (50 mM, 5 µL 1 M stock in DMF) were dissolved in 40 µL DMF and 43 µL Tris buffer (50 mM, pH 8.5). Reducing reagent TCEP (100 mM, 5 µL 2 M stock in water) was added to the mixture and the final pH value was adjusted to 8.5 by adding 2 µL 6M NaOH solution. The reaction mixture was incubated at 37°C. At indicated time points, 5 µL of the reaction mixture was taken out and mixed with 30 µL 0.5% TFA/$CH_3CN$ that was expected to quench the reaction. The sample was analyzed by LC/MS, and the relative product yield was determined the same as mentioned in the section of pH titration.

#### Measurement of Reaction Kinetics

Reaction kinetics were evaluated similarly to reported procedures.[1-2] Stock solutions of substrate **3** and nucleophiles were prepared in DMF, while TCEP stock (pH 7.4) was dissolved in $H_2O$. Equal concentrations (40 mM, 80 mM, or 160 mM) of the substrate and the nucleophile (3,4,5-trimethoxybenzenethiol or 2,4,6-trimethoxybenzenethiol) were mixed in 40 µL of DMF/Tris buffer (70/30). Over excess amount of TCEP (100 mM, 200 mM, or 400 mM) was added, and the final pH of the mixture was adjusted to 8.5 to initiate the reaction at 37 ºC. Time dependent measurements were carried out by taking 2 μL of the reaction mixture at indicated time points (30 min, 60 min, 90 min, 120 min, 150 min, 180 min), and mixing it with 18 μL 0.5% TFA/ACN to quench the reaction. The samples were eventually analyzed by LC/MS and the concentrations of product and reactant were determined by comparing peak area ratios with those of the standard curves. Plotting $1/[X]_t$ against time yielded the desired rate constant (k), based on the second order rate equation "$1/[X]_t = 1/[X]_0 + kt$" ($[X]_0$ : initial concentration of either reactant; t: reaction time; $[X]_t$ : concentration of either reactant at time t).

**Compound Characterization:**

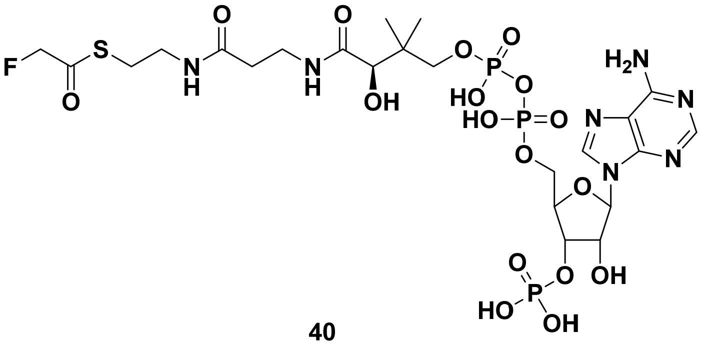


**40**

The fluoroacetyl-CoA (compound **40**) was synthesized and purified following the reported procedures.[3] $^1$H NMR (500 MHz, $D_2O$): $\delta$ 8.66 (s, 1H), 8.42 (s, 1H), 6.21 (d, $J$ = 5.5 Hz, 1H), 5.01 (d, $J$ = 46.5 Hz, 2H), 4.87-4.82 (m, 2H), 4.58 (s, 1H), 4.24 (s, 2H), 4.00 (s, 1H), 3.85-3.82 (m, 1H), 3.59-3.56 (m, 1H), 3.44 (t, $J$ = 6.5 Hz, 2H), 3.35 (t, $J$ = 6.0 Hz, 2H), 3.08 (t, $J$ = 6.5 Hz, 2H), 2.42 (t, $J$= 6.5 Hz, 2H), 0.92 (s, 3H), 0.79 (s, 3H). HRMS (ESI) m/z calculated for $C_{23}H_{38}FN_7O_{17}P_3S$ $[M + H]^+$: 828.1236, found 828.1229.

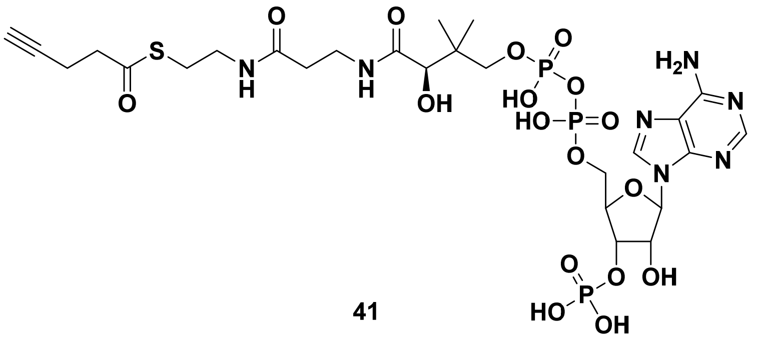


**41**

The 4-pentynoyl CoA (compound **41**) was prepared according to the published synthetic and purification procedures.[4] $^1$H NMR (500 MHz, $D_2O$): $\delta$ 8.66 (s, 1H), 8.45 (s, 1H), 6.22 (d, $J$ = 6.0 Hz, 1H), 4.61 (s, 1H), 4.28 (s, 2H), 4.02 (s, 1H), 3.90-3.86 (m, 1H), 3.65-3.61 (m, 1H), 3.46 (t, $J$

= 6.5 Hz, 2H), 3.35 (t, *J* = 6.0 Hz, 2H), 3.03 (t, *J* = 6.5 Hz, 2H), 2.84 (t, *J* = 7.0 Hz, 2H), 2.52-2.48 (m, 2H), 2.44 (t, *J*= 6.5 Hz, 2H), 2.34 (t, *J*= 2.5 Hz, 2H), 0.94 (s, 3H), 0.82 (s, 3H). HRMS (ESI) m/z calculated for $C_{26}H_{41}N_7O_{17}P_3S$ [M + H]$^+$: 848.1487, found 848.1477.

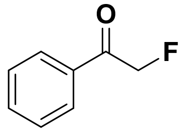


**1**

Dried potassium fluoride (580 mg, 10 mmol) was added to a solution of 18-crown-6 (264 mg, 1 mmol) in anhydrous acetonitrile (6 mL). After stirred at room temperature for 20 min, 2-bromoacetophenone (**17**) (398 mg, 2 mmol) in anhydrous acetonitrile (2 mL) was added and then heated to reflux and stirred overnight. After being concentrated under reduced pressure, the mixture was purified via flash column chromatography (hexane/ethyl acetate: 6/1) to afford compound **1** as a light yellow oil (210 mg, 1.52 mmol, 76% yield). $^1$H NMR (500 MHz, $CDCl_3$): $\delta$ 7.90 (d, *J* = 8.0 Hz, 2H), 7.64-7.61 (m, 1H), 7.52-7.48 (m, 2H), 5.54 (d, *J* = 47.0 Hz, 2H); $^{13}$C NMR (126 MHz, $CDCl_3$): $\delta$ 193.4 (d, *J* = 15.5 Hz), 134.2, 133.7, 129.0, 127.8 (d, *J* = 2.5 Hz), 83.6 (d, *J* = 182.8 Hz); $^{19}$F NMR (471 MHz, $CDCl_3$): $\delta$ -230.75; GC-MS m/z calculated for $C_8H_7FO$ [M]$^+$: 138.0 found 138.0.

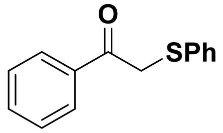


**42**

General procedure A was followed from **1** (28 mg, 0.2 mmol) in the presence of potassium carbonate to give **42** (42 mg, 91% yield) as a colorless oil after flash column chromatography (hexane/ethyl acetate: 10/1). $^1$H NMR (500 MHz, $CDCl_3$): $\delta$ 7.96-7.93 (m, 2H), 7.60-7.56 (m, 1H), 7.48-7.44 (m, 2H), 7.40-7.38 (m, 2H), 7.30-7.26 (m, 2H), 7.24-7.20 (m, 1H), 4.28 (s, 2H); $^{13}$C NMR (126 MHz, $CDCl_3$): $\delta$ 194.1, 135.4, 134.8, 133.5, 130.6, 129.1, 128.7, 127.2, 41.3; MS (ESI) m/z calculated for $C_{14}H_{13}OS$ [M + H]$^+$: 229.1, found 229.1.

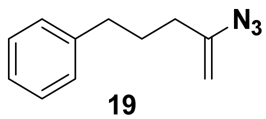


**19**

To a solution of pent-4-yn-1-ylbenzene (**18**) (455 μL, 3 mmol), trimethylsilyl azide (788 μL, 6 mmol), water (108 μL, 6 mmol), DMSO (10 mL), and silver carbonate (83 mg, 0.3 mmol) were added. The mixture was then stirred at 80 ºC for 1h. After cooled down to room temperature, water was added. The aqueous phase was extracted with ethyl acetate. The combined organic phase was washed with brine, then water, dried over anhydrous sodium sulfate and concentrated under reduced pressure. The residue was purified by flash column chromatography (100% hexane) to provide the desired compound **19** as a colorless oil (338 mg, 1.8 mmol, 60% yield). $^1$H NMR (500 MHz, $CDCl_3$): $\delta$ 7.31-7.28 (m, 2H), 7.22-7.18 (m, 3H), 4.67-4.66 (m, 2H), 2.64 (t, $J$ = 7.5 Hz, 2H), 2.11 (t, $J$ = 7.5 Hz, 2H), 1.86-1.79 (m, 2H); $^{13}$C NMR (126 MHz, $CDCl_3$): $\delta$ 146.5, 141.8, 128.5, 128.4, 125.9, 98.4, 35.0, 33.2, 28.9; HRMS (ESI) m/z calculated for $C_{11}H_{14}N_3$ [M + H]$^+$: 188.1182, found 188.1180.

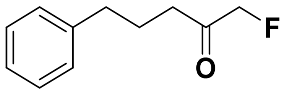


**2**

Vinyl azide (**19**) (187 mg, 1 mmol) was added to a suspension of Selectfluor (480 mg, 1.5 mmol), sodium bicarbonate (168 mg, 2 mmol), and water (36 µL, 2 mmol) in acetonitrile (10 mL). The resulting mixture was stirred at room temperature overnight. After concentrated under reduced pressure, the residue was purified by flash column chromatography (hexane/ethyl acetate: 20/1) to give compound **2** as a colorless oil (86 mg, 0.48 mmol, 48% yield). $^1$H NMR (500 MHz, CDCl3): $\delta$ 7.31-7.27 (m, 2H), 7.22-7.17 (m, 3H), 4.75 (d, $J$ = 48 Hz, 2H), 2.66 (t, $J$ = 7.5 Hz, 2H), 2.55 (dt, $J$ = 7.5, 2.5 Hz, 2H), 1.97 (quint, $J$ = 7.5 Hz, 2H); $^{13}$C NMR (126 MHz, $CDCl_3$): $\delta$ 206.8 (d, $J$ = 20.2 Hz), 141.2, 128.48, 128.47, 126.1, 85.0 (d, $J$ = 185.2 Hz), 37.4, 35.0, 24.1 (d, $J$ = 1.8 Hz); $^{19}$F NMR (471 MHz, $CDCl_3$): $\delta$ -227.52; MS (ESI) m/z calculated for $C_{11}H_{14}FO$ [M + Na]$^+$: 203.0, found 203.1.

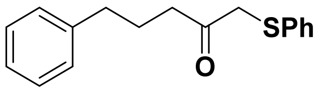


**43**

General procedure A was followed from **2** (36 mg, 0.2 mmol) in the presence of DBU to give **43** (53 mg, 98% yield) as a colorless oil after flash column chromatography (hexane/ethyl acetate: 10/1). $^1$H NMR (500 MHz, $CDCl_3$): $\delta$ 7.37-7.16 (m, 10H), 3.68 (s, 2H), 2.63 (t, $J$ = 7.5 Hz, 2H), 2.62 (t, $J$ = 7.5 Hz, 2H), 1.94 (quint, $J$ = 7.5 Hz, 2H); $^{13}$C NMR (126 MHz, $CDCl_3$): $\delta$ 205.4, 141.5, 134.9, 129.6, 129.2, 128.5, 128.4, 126.9, 126.0, 44.0, 39.8, 35.0, 25.2; MS (ESI) m/z calculated for $C_{17}H_{19}OS$ [M + H]$^+$: 271.1, found 271.1.

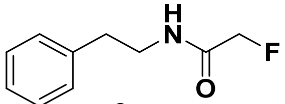


**3**

Sodium fluoroacetate (100 mg, 1 mmol) was mixed with 1-[Bis(dimethylamino)methylene]-1H-1,2,3-triazolo[4,5-b]pyridinium 3-oxid hexafluorophosphate (HATU) (456 mg, 1.2 mmol) and DIPEA (209 µL, 1.2 mmol) in DMF (5 mL). After the mixture was stirred at room temperature for 20 min, phenethylamine (**20**) (251 µL, 2 mmol) was added dropwisely. The resulting mixture was continuously stirred at room temperature overnight, and then quenched by water. Ethyl acetate was added to extract the product from aqueous layer. The organic layer was dried with anhydrous sodium sulfate and concentrated under vacuum. The crude mixture was then purified via flash column chromatography (hexane/ethyl acetate: 3/1) to afford compound **3** as a white solid (147 mg, 0.81 mmol, 81% yield). $^{1}$H NMR (500 MHz, $CDCl_3$): $\delta$ 7.36-7.33 (m, 2H), 7.28-7.22 (m, 3H), 6.37 (br, 1H), 4.79 (d, $J$ = 47.5 Hz, 2H), 3.62 (q, $J$ = 6.5 Hz, 2H), 2.89 (t, $J$ = 7.5 Hz, 2H); $^{13}$C NMR (126 MHz, $CDCl_3$): $\delta$ 167.5 (d, $J$ = 17.1 Hz), 138.4, 128.8, 128.7, 126.7, 80.3 (d, $J$ = 186.1 Hz), 40.0, 35.6; $^{19}$F NMR (471 MHz, $CDCl_3$): $\delta$ -227.23; HRMS (ESI) m/z calculated for $C_{10}H_{13}FNO$ $[M + H]^+$: 182.0976, found 182.0980.

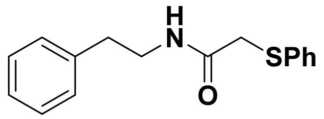

**44**

General procedure A was followed from **3** (37 mg, 0.2 mmol) in the presence of DBU to generate **44** (50 mg, 93% yield) as a white solid after flash column chromatography (hexane/ethyl acetate: 2/1). $^{1}$H NMR (500 MHz, $CDCl_3$): $\delta$ 7.30-7.17 (m, 8H), 7.06-7.04 (m, 2H), 3.61 (s, 2H), 3.51 (q, $J$ = 7.0 Hz, 2H), 2.73 (t, $J$ = 6.5 Hz, 2H); $^{13}$C NMR (126 MHz, $CDCl_3$): $\delta$ 167.7, 138.5, 134.7, 129.3, 128.70, 128.66, 127.8, 126.58, 126.55, 40.9, 37.2, 35.5; MS (ESI) m/z calculated for $C_{16}H_{18}NOS$ $[M + H]^+$: 272.1, found 272.1.

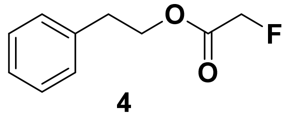

**4**

2-Phenylethanol (**21**) (359 µL, 3 mmol) was dissolved in DCM (10 mL), and mixed with potassium carbonate (828 mg, 6 mmol) in 2 mL water. The mixture was cooled in an ice bath, and a solution of bromoacetyl bromide (392 µL, 4.5 mmol) in DCM (3 mL) was dropwisely added. After 30 min of stirring, the reaction solution was warmed to room temperature and continuously stirred for another 2 h. The aqueous layer was then separated and extracted with DCM. The combined organic phase was washed with brine and water. After drying with anhydrous sodium sulfate, the organic layer was vacuum concentrated to yield an oily intermediate. After resuspension in THF, the oily intermediate was mixed with TBAF (6 mL, 6 mmol, 1 M in THF), refluxed for 1h, and concentrated under reduced pressure. The crude mixture was purified via flash column chromatography (hexane/ethyl acetate: 15/1) to afford colorless oil-like compound **4** (337 mg, 1.85 mmol, 62% yield over two steps). $^{1}$H NMR (500 MHz, $CDCl_3$): $\delta$ 7.33-7.29 (m, 2H), 7.26-7.20 (m, 3H), 4.81 (d, $J$ = 47.0 Hz, 2H), 4.43 (t, $J$ = 7.5 Hz, 2H), 2.98 (t, $J$ = 7.0 Hz, 2H); $^{13}$C NMR (126 MHz, $CDCl_3$): $\delta$ 167.8 (d, $J$ = 21.8 Hz), 137.2, 128.9, 128.6, 126.8, 78.3 (d, one peak overlap with $CDCl_3$), 65.8, 35.0; $^{19}$F NMR (471 MHz, $CDCl_3$): $\delta$ -229.98. HRMS (ESI) m/z calculated for $C_{10}H_{12}FO_2$ $[M + H]^+$: 183.0816, found 183.0862.

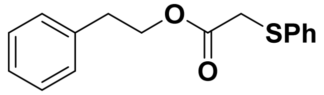

**45**

General procedure A was followed from **4** (37 mg, 0.2 mmol) in the presence of potassium carbonate to afford **45** (8 mg, 14% yield) as a colorless oil after flash column chromatography (hexane/ethyl acetate: 20/1). $^{1}$H NMR (500 MHz, $CDCl_3$): $\delta$ 7.38-7.35 (m, 2H), 7.30-7.26 (m, 4H), 7.24-7.17 (m, 4H), 4.32 (t, $J$ = 7.5 Hz, 2H), 3.63 (s, 2H), 2.90 (t, $J$ = 7.0 Hz, 2H); $^{13}$C NMR (126 MHz, $CDCl_3$): $\delta$ 169.7, 137.5, 135.0, 129.9, 129.1, 128.9, 128.6, 127.0, 126.6, 66.0, 36.7, 34.9; MS (ESI) m/z calculated for $C_{16}H_{17}O_2S$ $[M + H]^+$: 273.1, found 273.1.

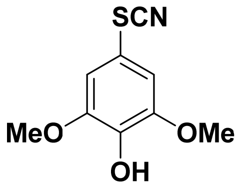

**23**

Bromine (774 µL, 15 mmol) was slowly added to the saturated solution of sodium bromide in methanol (10 mL) at 0 ºC. After stirring for 15 min, the solution was added dropwisely to a mixture of 2,6-dimethoxypheno1(**22**) (1.54 g, 10 mmol) and potassium thiocyanate (1.46 g, 15 mmol) in 0 ºC methanol (30 mL). The reaction was left on for 1h, and then quenched by water. The methanol in the mixture was evaporated under reduced pressure, and the remaining aqueous phase was extracted with ethyl acetate. The combined organic layer was washed with brine, and water, dried with anhydrous $Na_2SO_4$, and eventually concentrated to afford an orange colored oil-like mixture. The crude mixture was purified via flash column chromatography (hexane/ethyl acetate: 5/1) to result in compound **23** as an orange colored solid (1.14 g, 5.4 mmol, 54% yield). $^{1}$H NMR (500 MHz, $CDCl_3$): $\delta$ 6.80 (s, 2H), 5.74 (s, 1H), 3.91 (s, 6H); $^{13}$C NMR (126 MHz, $CDCl_3$): $\delta$ 147.9, 136.9, 112.7, 111.6, 109.3, 56.8; MS (ESI) m/z calculated for $C_9H_{10}NO_3S$ $[M + H]^+$: 212.0, found 212.0.

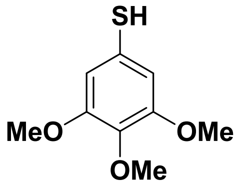

**12**

Potassium carbonate (276 mg, 2 mmol) and intermediate **23** (211 mg, 1 mmol) were dissolved in DMF (20 mL) and cooled to 0 ºC. Methyl iodide (125 µL, 2 mmol) was added dropwisely, after which the reaction was stirred at room temperature until TLC analysis confirmed the completion of reaction. The mixture was then diluted with ethyl acetate and water. After extraction and separation, the organic layer was washed with brine and water, dried with anhydrous sodium sulfate, and finally concentrated under reduced pressure to afford the methylated intermediate. The crude mixture was dissolved in THF (20 mL), added with the aqueous solution of lithium hydroxide (1M, 2 mL). After stirring at room temperature for 2h, the reaction was quenched by water, and the crude product was extracted by ethyl acetate. The organic layer was dried with anhydrous sodium sulfate, vacuum concentrated, and subsequently purified via flash column chromatography (hexane/ethyl acetate: 5/1) to afford compound **12** in yellow colored solid form (72% yield over two steps). $^{1}$H NMR (500 MHz, $CDCl_3$): $\delta$ 6.73 (s, 2H), 3.81(s, 3H), 3.80 (s, 6H); $^{13}$C NMR (126 MHz, $CDCl_3$): $\delta$ 153.4, 138.0, 132.0, 106.5, 60.9, 56.2; MS (ESI) m/z calculated for $C_9H_{13}O_3S$ $[M + H]^+$: 201.0, found 201.1.

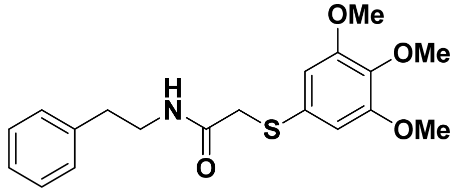


**46**

Prepared according to General Procedure B using 3,4,5-trimethoxybenzenethiol (**12**) (20 mg, 0.1 mmol) to give product **46** (16 mg, 88% yield) as white solid after flash column chromatography (hexane/ethyl acetate: 1/1). $^{1}$H NMR (500 MHz, $CDCl_3$): $\delta$ 7.25-7.17 (m, 3H), 7.07-7.05 (m, 2H), 6.77 (t, $J$ = 6.0 Hz, 1H), 6.45 (s, 1H), 3.82(s, 3H), 3.81 (s, 6H), 3.59 (s, 2H), 3.52 (q, $J$ = 7.0 Hz, 2H), 2.75 (t, $J$ = 7.0 Hz, 2H); $^{13}$C NMR (126 MHz, $CDCl_3$): $\delta$ 167.9, 153.7, 138.5, 129.5, 128.66, 128.64, 126.6, 105.37, 105.34, 61.0, 56.2, 41.0, 37.9, 35.6; HRMS (ESI) m/z calculated for $C_{19}H_{24}NO_4S$ $[M + H]^+$: 362.1421, found 362.1413.

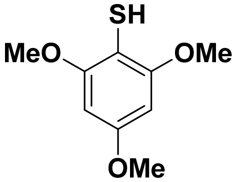


**13**

To a solution of 2,4,6-trimethoxybenzene **(24)** (504 mg, 3.0 mmol) in THF (10 mL), n-butyllithium in hexane (2.5 M, 1.2 mL, 3.0 mmol) was added at 0 °C, followed with a catalytic amount of tetramethylethylenediamine (TMEDA) (23 µL, 0.15 mmol). The reaction mixture was warmed up to room temperature, and stirred for 1 h until the suspension turned an orange color. Elemental sulfur (96 mg, 3 mmol) in warm toluene (3 mL) was added dropwisely, and the reaction mixture was stirred at room temperature. Upon reaction completion as monitored by TLC, 10 mL of water was added to quench the reaction, and the aqueous layer was acidified by 1 M HCl. The products were extracted by ethyl acetate, washed with water, brine, and dried over sodium sulfate. After vacuum concentration, the crude mixture was purified via flash column chromatography (hexane/ethyl acetate: 4/1) to afford the final product **13** as a crystalline yellow solid (360 mg, 60% yield). $^1$H NMR (500 MHz, $CDCl_3$): $\delta$ 6.17 (s, 2H), 3.87(s, 6H), 3.80 (s, 3H), 3.77 (s, 1H); $^{13}$C NMR (126 MHz, $CDCl_3$): $\delta$ 158.7, 156.2, 99.7, 91.2, 56.1, 55.5; MS (ESI) m/z calculated for $C_9H_{13}O_3S$ $[M + H]^+$: 201.1, found 201.1.

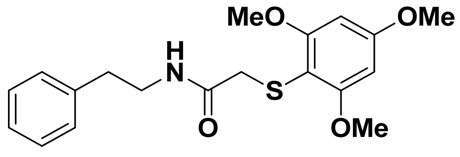


**47**

Prepared according to General Procedure B using 2,4,6-trimethoxybenzenethiol (**13**) (20 mg, 0.1 mmol) to generate product **47** (15 mg, 83% yield) as white solid after flash column chromatography (hexane/ethyl acetate: 2/1). $^1$H NMR (500 MHz, $CDCl_3$): $\delta$ 7.88 (s, 1H), 7.30-7.27 (m, 2H), 7.23-7.20 (m, 1H), 7.18-7.16 (m, 2H), 6.10 (s, 2H), 3.82(s, 3H), 3.78 (s, 6H), 3.47 (s, 2H), 3.44 (q, $J$ = 7.0 Hz, 2H), 2.76 (t, $J$ = 7.0 Hz, 2H); $^{13}$C NMR (126 MHz, $CDCl_3$): $\delta$ 169.1, 162.4, 161.5, 139.1, 128.7, 128.6, 126.4, 100.1, 91.2, 56.1, 55.5, 41.3, 38.6, 35.9; HRMS (ESI) m/z calculated for $C_{19}H_{24}NO_4S$ $[M + H]^+$: 362.1421, found 362.1415.

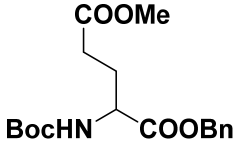


**26**

Boc-L-glutamic acid 1-benzyl ester **(25)** (6.0 g, 17.7 mmol) and sodium bicarbonate (3.68 g, 26.7 mmol) were dissolved in DMF (60 mL) and cooled to 0 ºC. Methyl iodide (2.21 mL, 35.6 mmol) was added dropwisely, after which the reaction was stirred at room temperature until TLC analysis confirmed reaction completion. Upon completion, the reaction mixture was diluted 10-fold with water and extracted with ethyl acetate. The organic layer was washed with a 10% sodium bicarbonate solution, followed by brine and was subsequently dried with anhydrous sodium sulfate. After vacuum concentration, the crude mixture was purified via flash column chromatography (hexane/ethyl acetate: 2/1) to afford compound **26** as a clear oil (6.25 g, quantitative yield). $^1$H NMR (500 MHz, $CDCl_3$): $\delta$ 7.38-7.31 (m, 5H), 5.16 (d, $J$ = 4.0 Hz, 2H), 5.12 (m, 1H), 4.37 (d, $J$ = 5.0 Hz, 1H), 3.66 (s, 3H), 2.44-2.31 (m, 2H), 2.23-2.17 (m, 1H), 2.00-1.92 (m, 1H), 1.42 (s, 9H); $^{13}$C NMR (126 MHz, $CDCl_3$): $\delta$ 173.2, 172.1, 155.4, 135.3, 128.7, 128.5, 128.3, 80.1, 67.3, 53.0, 51.8, 30.1, 28.3, 27.8; MS (ESI) m/z calculated for $C_{18}H_{25}NNaO_6$ $[M + Na]^+$: 374.2, found 374.2.

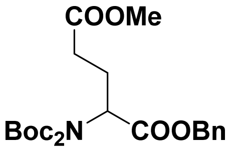

**27**

To a solution of intermediate **26** (6.25 g, 17.7 mmol) and 4-dimethylaminopyridine (DMAP)(435 mg, 3.5 mmol) in acetonitrile was added di-tert-butyl dicarbonate (7.76 g, 35.4 mmol). The reaction mixture was stirred overnight and directly vacuum concentrated upon completion as monitored by TLC. The concentrated crude mixture was purified via flash column chromatography (hexane/ethyl acetate: 4/1) to afford compound **27** as a clear oil (7.63 g, 95% yield). $^1$H NMR (500 MHz, $CDCl_3$): $\delta$ 7.33-7.27 (m, 5H), 5.14 (d, $J$ = 2.5 Hz, 2H), 4.97 (q, $J$ = 5.0 Hz, 1H), 3.66 (s, 3H), 2.53-2.46 (m, 1H), 2.43-2.35 (m, 2H), 2.24-2.16 (m, 1H), 1.44 (s, 18H); $^{13}$C NMR (126 MHz, $CDCl_3$): $\delta$ 173.1, 170.2, 152.0, 135.6, 128.5, 128.2, 128.0, 83.3, 66.9, 57.5, 51.7, 30.6, 27.9, 24.8; MS (ESI) m/z calculated for $C_{23}H_{33}NNaO_8$ $[M + Na]^+$: 474.2, found 474.2.

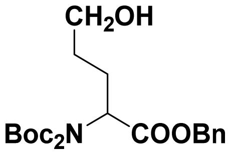

**28**

In a flame dried flask under a nitrogen atmosphere, a solution of intermediate **27** (7.63 g, 16.9 mmol) in THF (80 mL) was cooled to -80 °C. Diisobutylaluminum hydride solution (DIBAL)(1.0 M in hexanes) (33.8 mL) was added dropwisely over 30 min. The reaction mixture was stirred at -80 °C for at least 2 h. Upon completion as monitored by TLC, the reaction was quenched with a saturated Rochelle salt solution in water (200 mL), and was stirred at room temperature overnight. On the next day, the reaction mixture was diluted further with water (100 mL) and extracted with ethyl acetate. The organic layer was then dried with sodium sulfate and vacuum concentrated. The crude reaction mixture was purified via flash column chromatography (hexane/ethyl acetate: 2/1) to afford compound **28** as a clear oil (5.0 g, 70% yield). $^1$H NMR (500 MHz, $CDCl_3$): $\delta$ 7.34-7.29 (m, 5H), 5.14 (q, $J$ = 12.5 Hz, 2H), 4.91 (d, $J$ = 9.5, 5.5 Hz, 1H), 3.66 (t, $J$ = 6.5 Hz, 2H), 2.28-2.21 (m, 1H), 1.99-1.91 (m, 1H), 1.66-1.1.62 (m, 2H), 1.44 (s, 18H); $^{13}$C NMR (126 MHz, $CDCl_3$): $\delta$ 170.8, 152.3, 135.7, 128.5, 128.1, 128.0, 83.2, 66.8, 62.3, 58.0, 29.4, 27.9, 26.0; HRMS (ESI) m/z calculated for $C_{12}H_{18}NO_3$ (without di-Boc) $[M + H]^+$: 224.1281, found 224.1279.

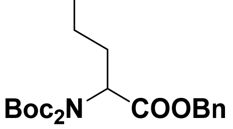

**29**

Intermediate **28** (5.0 g, 11.8 mmol), triphenylphosphine (4.64 g, 17.7 mmol), and imidazole (1.2 g, 17.7 mmol) were dissolved in DCM (60 mL) and stirred. Once dissolved, iodine (5.99 g, 23.6 mmol) was added, and the reaction mixture was stirred overnight. Upon completion, the reaction was quenched with saturated sodium sulfite (75 mL), and the organic products were extracted with dichloromethane. The organic layer was dried with anhydrous sodium sulfate, concentrated under reduced pressure; and the resulting crude mixture was purified using flash column chromatography (hexane/ethyl acetate: 4/1). Compound **29** was finally obtained as a clear oil (5.04 g, 80% yield). $^{1}$H NMR (500 MHz, $CDCl_3$): *δ* 7.39-7.27 (m, 5H), 5.15 (q, *J* = 12.5 Hz, 2H), 4.90 (q, *J* = 5.0 Hz, 1H), 3.27-3.14 (m, 2H), 2.27-2.19 (m, 1H), 2.09-2.01 (m, 1H), 1.95-1.85 (m, 2H), 1.46 (s, 18H); $^{13}$C NMR (126 MHz, $CDCl_3$): *δ* 170.4, 152.2, 135.6, 128.5, 128.2, 128.0, 83.3, 66.9, 57.2, 30.5, 30.2, 28.0, 5.7; HRMS (ESI) m/z calculated for $C_{22}H_{33}INO_6$ $[M + H]^+$: 534.1347, found 534.2258.

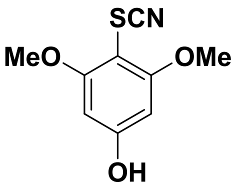


**31**

To a solution of N-chlorosaccharin (5.15 g, 23.7 mmol) in dichloromethane (90 mL) was added silver thiocyanate (3.93 g, 23.7 mmol). A white solid crashed out upon addition and the reaction mixture was kept stirring for 1 h. 3,5-dimethoxyphenol (compound **30**, 3.04 g ,19.6 mmol) was then added, and the reaction mixture was stirred for another 3 h, at which point the reaction was confirmed to be complete by TLC analysis. The dark red heterogeneous mixture was vacuum filtered, and the filtrate was vacuum concentrated to afford a dark red oil. The crude oil mixture was purified via flash column chromatography (hexane/ethyl acetate: 3/1) to afford compound **31** as an orange solid (2.50 g, 60% yield). $^{1}$H NMR (500 MHz, $CDCl_3$): *δ* 6.10 (s, 2H), 3.84 (s, 6H); $^{13}$C NMR (126 MHz, $CDCl_3$): *δ* 161.5, 160.9, 112.6, 92.9, 89.1, 56.4; MS (ESI) m/z calculated for $C_9H_{10}NO_3S$ $[M + H]^+$: 212.0, found 212.0.

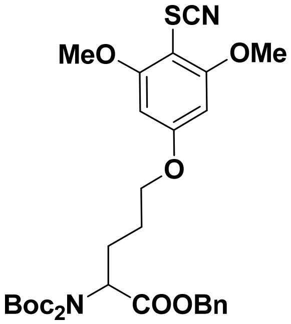


**32**

Intermediate **29** (5.0 g, 9.37 mmol), **31** (3.96 g, 18.7 mmol) and potassium carbonate (1.94 g, 14.0 mmol) were dissolved in DMF (50 mL) and stirred at room temperature for 8 h. Upon reaction completion, the mixture was diluted 10-fold with water, and the organic products were extracted by ethyl acetate. The organic layer was dried with sodium sulfate, vacuum concentrated; and the crude oil was purified via flash column chromatography (hexane/ethyl acetate: 3/1) to afford compound **32** as a yellowish oil (4.05 g, 70% yield). $^1$H NMR (500 MHz, $CDCl_3$): $\delta$ 7.34-7.28 (m, 5H), 6.13 (s, 1H), 5.15 (q, $J$ = 12.5 Hz, 2H), 4.95 (dd, $J$ = 9.5, 5.5 Hz, 1H), 4.00 (t, $J$ = 6.0 Hz, 2H), 3.89 (s, 6H), 2.38-2.30 (m, 1H), 2.12-2.05 (m, 1H), 1.91-1.84 (m, 2H), 1.45 (s, 18H); $^{13}$C NMR (126 MHz, $CDCl_3$): $\delta$ 170.6, 163.6, 161.4, 152.3, 135.6, 128.5, 128.2, 128.0, 111.9, 91.8, 83.3, 67.6, 66.9, 57.8, 56.4, 27.9, 26.0, 26.0; HRMS (ESI) m/z calculated for $C_{31}H_{41}N_2O_9S$ [M + H]$^+$: 617.2527, found 617.2522.

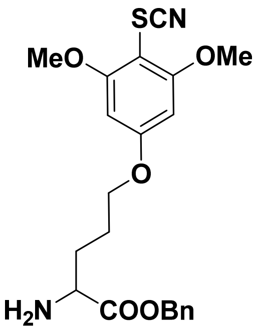


**33**

Intermediate **32** (4.0 g, 6.49 mmol) and triisopropylsilane (1.59 mL, 7.78 mmol) were dissolved in trifluoroacetic acid / dichloromethane (10 mL/10 mL) and stirred for 2 h. The reaction mixture was directly vacuum concentrated. The resulting crude oil was suspended in water and basified to pH=8 with saturated sodium bicarbonate (~ 40 mL). The organic products were extracted with ethyl acetate and dried with anhydrous sodium sulfate. The organic layer was concentrated under reduced pressure, and the crude oil was purified via flash column chromatography (dichloromethane/methanol: 50/1) to afford compound **33** are an orange oil (2.16 g, 80% yield). $^1$H NMR (500 MHz, $CDCl_3$): $\delta$ 7.30-7.27 (m, 5H), 6.05 (s, 2H), 5.09 (s, 2H), 3.91 (t, $J$ = 6.0 Hz, 2H), 3.81 (s, 6H), 3.51-3.48 (m, 1H), 1.89-1.71 (m, 4H); $^{13}$C NMR (126 MHz, $CDCl_3$): $\delta$ 175.7, 163.7, 161.5, 135.7, 128.8, 128.6, 128.4, 112.0, 91.9, 67.9, 66.9, 56.5, 543, 31.2, 25.5; HRMS (ESI) m/z calculated for $C_{21}H_{25}N_2O_5S$ [M + H]$^+$: 417.1479, found 417.1469.

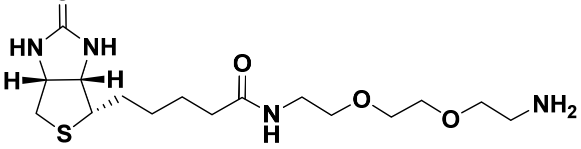


**35**

Intermediate **35** was synthesized according to published procedures.[5] Briefly, a solution of D-biotin (2.5 g, 10.2 mmol) (compound **34**) in DMF (60 mL) was stirred and heated at 60 ºC until fully dissolved. The coupling reagent 1,1′-Carbonyldiimidazole (CDI)(3.32 g, 20.5 mmol) was then added, and the reaction mixture was kept stirrring at 60 ºC for 3 h, after which the linker 2,2'-(Ethylenedioxy)bis(ethylamine) (5.96 mL, 40.9 mmol) was added. The reaction mixture was stirred overnight at room temperature, and then vacuum concentrated. The crude oil was purified via flash column chromatography (dichloromethane/methanol: 5/1, plus 1% triethylamine) to render compound **35** (3.65 g, 95% yield) as a yellowish oil. $^1$H NMR (500 MHz, $D_2O$): $\delta$ 4.49 (dd, $J$ = 8.0, 5.0 Hz, 1H), 4.30 (dd, $J$ = 7.5, 4.5 Hz, 1H), 3.59-3.50 (m, 7H), 3.41 (t, $J$ = 5.5 Hz, 1H), 3.37 (t, $J$ = 5.5 Hz, 1H), 3.23-3.19 (m, 1H), 2.93 (dd, $J$ = 13.0, 5.0 Hz, 1H), 2.86 (t, $J$ = 5.0 Hz, 2H), 2.70 (d, $J$ = 13.0 Hz, 1H), 2.22 (t, $J$ = 7.5 Hz, 2H), 1.78-1.56 (m, 4H), 1.47-1.41 (m, 2H). MS (ESI) m/z calculated for $C_{16}H_{31}N_4O_4S$ $[M + H]^+$: 375.2, found 375.2.

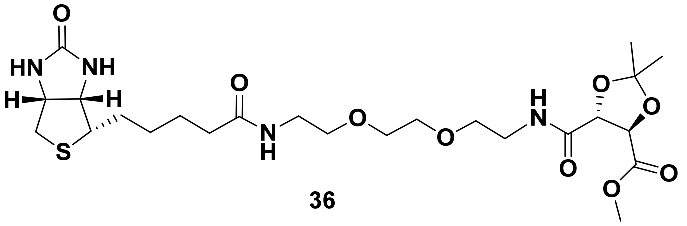


**36**

The coupling agent HATU (8.38 g, 22.0 mmol) was mixed with the commercially available building block (4R,5R)-5-(Methoxycarbonyl)-2,2-dimethyl-1,3-dioxolane-4-carboxylic acid (3.0 g, 14.7 mmol) in 40 mL DMF. Compound **35** (8.25 g, 22.0 mmol) was added, and the reaction mixture was stirred for 10 minutes, followed by the addition of N,N-Diisopropylethylamine (5.12 mL, 29.4 mmol) and the reaction mixture was stirred overnight. The reaction mixture was directly vacuum concentrated and purified via flash chromatography (dichloromethane/methanol: 20/1) to afford compound **36** as an orange oil (6.16 g, 75% yield). $^1$H NMR (500 MHz, $CD_3OD$) $\delta$ 4.72 (s, 2H), 4.51 (dd, $J$ = 8.0, 4.5 Hz, 1H), 4.32 (dd, $J$ = 8.0, 4.5 Hz, 1H), 3.82 (s, 3H), 3.67-3.62 (m, 4H), 3.61 (t, $J$ = 5.5 Hz, 2H), 3.56 (t, $J$ = 5.5 Hz, 2H), 3.53-3.47 (m, 1H), 3.46-3.41 (m, 1H), 3.38 (q, $J$ = 6.0 Hz, 2H), 3.25-3.21 (m, 1H), 2.94 (dd, $J$ = 13.0, 5.0 Hz, 1H), 2.72 (d, $J$ = 13.0 Hz, 1H), 2.24 (t, $J$ = 7.5 Hz, 2H), 1.80-1.58 (m, 4H), 1.50 (s, 3H), 1.47 (s, 3H), 1.12 (d, $J$ = 6.5 Hz, 4H); $^{13}$C NMR (126 MHz, $CD_3OD$) $\delta$ 174.8, 170.8, 170.5, 164.7, 113.3, 78.0, 77.2, 69.9, 69.9, 69.2, 69.0, 62.0, 60.2, 55.6, 51.8, 41.3, 39.6, 38.9, 38.6, 35.3, 28.4, 28.1, 25.5, 25.2, 22.1; HRMS (ESI) m/z calcd for $C_{24}H_{41}N_4O_9S$ $[M + H]^+$ 561.2589, found 561.2583.

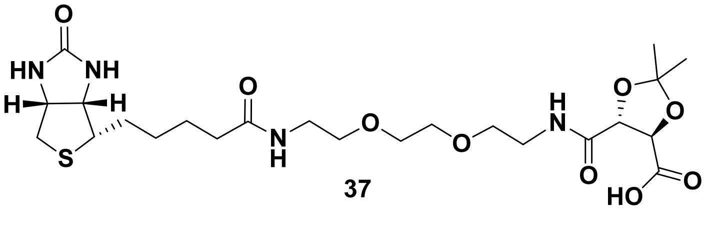


**37**

Lithium hydroxide (2 M, 3.2 mL) was slowly added into a solution of compound **36** (3.0 g, 5.35 mmol) in 20 mL methanol at 0 ºC. The reaction mixture was stirred for 2 h and acidified with 1 M HCl (~5 mL). The crude mixture was vacuum concentrated and purified via high-performance liquid chromatography (HPLC) to afford compound **37** (2.62 g, 90% yield) as a white solid. For HPLC purification (flow rate: 10 mL/min), solvent A is 0.1% TFA containing water while solvent B is 0.1% TFA containing acetonitrile. After the initial 5 min post sample injection, solvent B percentage was increased linearly to 100% within 35 min. The system was continuously flushed with 100% solvent B for another 5 min before the run stopped. Compound peak retention time on HPLC: ~ 21 min. $^{1}$H NMR (500 MHz, $CD_3OD$): $\delta$ 4.58 (dd, $J$ = 6.0 Hz, 2H), 4.39 (dd, $J$ = 8.0, 4.5 Hz, 1H), 4.21 (dd, $J$ = 8.0, 4.5 Hz, 1H), 3.54-3.50 (m, 4H), 3.49 (t, $J$ = 5.5 Hz, 2H), 3.44 (t, $J$ = 5.5 Hz, 2H), 3.41-3.36 (m, 1H), 3.34-3.29 (m, 1H), 3.26 (t, $J$ = 5.5 Hz, 2H), 3.13-3.09 (m, 1H), 2.83 (dd, $J$ = 13.0, 5.0 Hz, 1H), 2.60 (d, $J$ = 12.5 Hz, 1H), 2.13 (t, $J$ = 7.5 Hz, 2H), 1.68-1.47 (m, 4H), 1.38(s, 3H), 1.36 (s, 3H); $^{13}$C NMR (126 MHz, $CD_3OD$): $\delta$ 174.8, 172.2, 170.9, 164.7, 112.9, 77.9, 77.3, 69.92, 69.88, 69.2, 69.0, 62.0, 60.2, 55.6, 39.6, 38.9, 38.7, 35.3, 28.4, 28.1, 25.4, 25.2; HRMS (ESI) m/z calculated for $C_{23}H_{39}N_4O_9S$ $[M + H]^+$: 547.2432, found 547.2424.

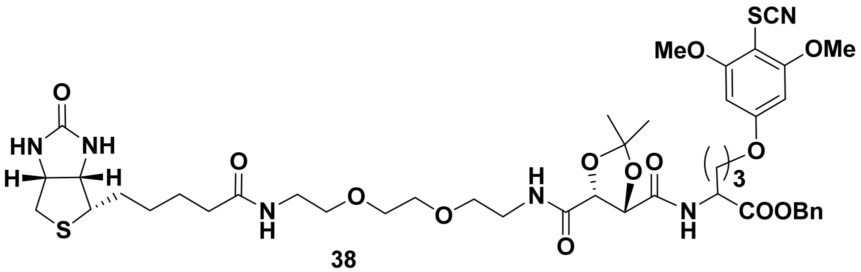


38

Compound **37** (2.0 g, 3.66 mmol) and HATU (2.1 g, 5.49 mmol) were dissolved in DMF (18 mL). Intermediate **33** (0.91 g, 2.19 mmol) was then added and the reaction mixture was stirred for 10 min, followed by the addition of N, N-diisopropylethylamine (DIPEA) (0.57 mL, 3.29 mmol). The reaction mixture was stirred overnight. Upon completion, the mixture was diluted 10-fold with water, and the organic products were extracted with ethyl acetate. The organic layer was dried with sodium sulfate, vacuum concentrated; and the crude mixture was eventually purified via flash column chromatography (dichloromethane/methanol: 30/1)) to afford compound **38** in light-orange solid form (1.64 g, 80% yield). $^{1}$H NMR (500 MHz, $CD_3OD$): $\delta$ 7.58 (d, $J$ = 8.5 Hz, 1H), 7.36-7.29 (m, 5H), 6.67 (t, $J$ = 5.5 Hz, 1H), 6.11 (s, 2H), 5.21-5.12 (m, 2H), 4.75-4.70 (m, 1H), 4.63 (d, $J$ = 6.5 Hz, 1H), 4.54 (d, $J$ = 6.5 Hz, 1H), 4.48-4.45 (m, 1H),4.29-4.26 (m, 1H), 3.97 (t, $J$ = 6.0 Hz, 2H), 3.87 (s, 6H), 3.59-3.50 (m, 8H), 3.42-3.37 (m, 2H), 3.13-3.08 (m, 1H), 2.88-2.84 (m, 1H), 2.69 (d, $J$ = 13.0 Hz, 1H), 2.20 (t, $J$ = 7.5 Hz, 2H), 2.14-2.08 (m, 2H), 1.95-1.88 (m, 1H), 1.86-1.76 (m, 2H), 1.73-1.61 (m, 5H), 1.47 (s, 3H), 1.45 (s, 3H); $^{13}$C NMR (126 MHz, $CD_3OD$): $\delta$ 173.8, 171.4, 170.2, 169.9, 164.0, 163.5, 161.4, 135.1, 128.7, 128.6, 128.4, 112.9, 111.9, 91.8, 70.1, 70.0, 69.6, 67.5, 67.3, 61.8, 60.3, 56.4, 55.5, 55.4, 53.5, 51.7, 50.7, 40.5, 39.2, 39.1, 35.8, 28.9, 28.1, 28.0, 26.2, 26.1, 25.6, 25.0; HRMS (ESI) m/z calculated for $C_{44}H_{61}N_6O_{13}S_2$ $[M + H]^+$: 945.3733, found 945.3722.

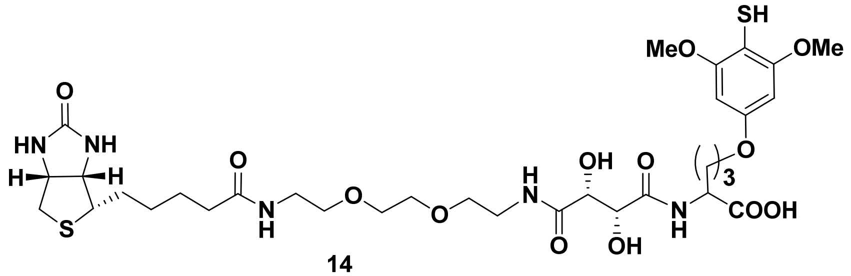

**14**

Compound **38** (1.0 g, 1.06 mmol) was dissolved in acetic acid/water (9 mL/1 mL) and refluxed for ~ 24 h, at which point the complete diol deprotection was confirmed by LC-MS analysis. The reaction mixture was then vacuum concentrated to afford the diol intermediate as a white solid. The diol intermediate was resuspended in THF (8 mL) at 0 ºC, and 2 M lithium hydroxide (2.09 mmol, ~ 1.1 mL) in water was added dropwisely. The reaction mixture was stirred for ~ 2 h, at which point the hydrolysis was confirmed complete by TLC. The mixture was acidified with 1 M HCl (~ 3 mL), vacuum concentrated, and finally purified by HPLC. The HPLC purification utilized water (0.1% TFA) as solvent A and acetonitrile (0.1% TFA) as solvent B. The flow rate was 10 mL/min. For the first 5 min post injection, 100% solvent A was used in the program. After that, solvent B percentage was linearly increased to 50% within 55 min. Then the HPLC flow was changed to 100% solvent B within the next 20 min, followed by additional 5 min flushing with solvent B percentage remaining at 100%. Compound peak retention time on HPLC was ~ 58 min. The eluted fraction was lyophilized to afford compound **14** as a white solid (334 mg, 40% yield). $^1$H NMR (500 MHz, DMSO-d$_6$): $\delta$ 7.84 (t, $J$ = 5.5 Hz, 1H), 7.80 (d, $J$ = 8.0 Hz, 1H), 7.64 (t, $J$ = 6.0 Hz, 1H), 6.29 (s, 2H), 4.35-4.28 (m, 3H), 4.24 (d, $J$ = 2.0 Hz, 1H), 4.12 (dd, $J$ = 7.5, 4.5 Hz, 1H), 3.97 (t, $J$ = 6.0 Hz, 1H), 4.48-4.45 (m, 1H),4.29-4.26 (m, 1H), 3.97 (t, $J$ = 6.0 Hz, 2H), 3.80 (s, 6H), 3.53-3.48 (m, 4H), 3.44 (t, $J$ = 6.0 Hz, 2H), 3.39 (t, $J$ = 6.0 Hz, 2H), 3.35-3.28 (m, 1H), 3.26-3.22 (m, 1H), 3.21-3.14 (m, 2H), 3.13-3.07 (m, 1H), 2.82 (dd, $J$ = 7.5, 5.0 Hz, 1H), 2.59-2.54 (m, 1H), 2.07-2.04 (m, 2H), 1.97-1.91 (m, 1H), 1.83-1.75 (m, 1H), 1.74-1.67 (m, 2H), 1.63-1.56 (m, 1H), 1.52-1.41 (m, 3H), 1.33-1.28 (m, 2H); $^{13}$C NMR (126 MHz, DMSO-d$_6$): $\delta$ 173.6, 172.6, 172.3, 172.3, 163.2, 158.0, 156.0, 92.5, 73.0, 72.8, 70.0, 69.7, 69.4, 67.7, 61.5, 59.7, 56.6, 55.9, 54.1, 51.8, 38.9, 38.8, 35.6, 28.7, 28.5, 25.7, 25.4, 18.6, 17.2; HRMS (ESI) m/z calculated for $C_{33}H_{52}N_5O_{13}S_2$ [M + H]$^+$: 790.2998, found 790.2988.

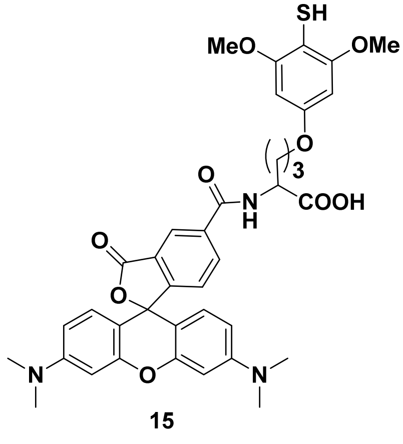

**15**

5-carboxytetramethylrhodamine (5-TAMRA, compound **39**, Thermo Fisher, 43 mg, 0.10 mmol) was dissolved in 2 mL DMF, and mixed with HATU (46.0 mg, 0.12 mmol). The reaction mixture was stirred for 10 min to ensure it was fully dissolved. The previously prepared **33** (42.0 mg, 0.10 mmol) was added to the solution, followed by DIPEA (53.0 μL, 0.30 mmol). The resulting mixture was stirred at RT for 3 h and was vacuum concentrated upon completion as confirmed by TLC. The crude mixture was purified via flash column chromatography (dichloromethane/methanol: 20/1) to afford the conjugated intermediate as purple oil, which was directly dissolved in THF (2 mL) and cooled down to 0 °C. Lithium hydroxide (2 M, 0.06 mL) in water was added dropwisely, and the solution was stirred at RT for 2 hours at which point the hydrolysis was near complete as confirmed by TLC. The mixture was then acidified with 1M HCL (~ 200 μL), and vacuum concentrated. The crude mixture was purified via HPLC that implemented water (0.1% TFA) as solvent A and acetonitrile (0.1% TFA) as solvent B, with a flow rate of 10 mL/min. The purification program was run as the following: 5.0% solvent B for the first minute, followed by a liner progression to 70.0% solvent B for the next 39 minutes, ending with the last 5 minutes of 100.0% solvent B for a total of 45 minutes per HPLC run. The product peak came out at 40 min, and lyophilization of the collected fraction rendered compound **15** (39.6 mg, 56% yield) as a pink solid. $^1$H NMR (500 MHz, $CD_3OD$): $\delta$ 8.76-8.74 (m, 1H), 8.23-8.19 (m, 1H), 7.48 (d, $J$ = 8.0 Hz, 1H), 7.14-7.10 (m, 2H), 7.06-7.03 (m, 2H), 6.97-6.93 (m, 2H), 6.25 (s, 2H), 4.78-4.73 (m, 1H), 4.10-4.04 (m, 2H), 3.80 (s, 6H), 3.28 (s, 12H), 2.30-2.22 (m, 1H), 2.12-2.03 (m, 1H), 2.00-1.94 (m, 2H); $^{13}$C NMR (126 MHz, $CD_3OD$): $\delta$ 175.3, 168.4, 167.3, 160.7, 159.2, 159.1, 159.0, 157.3, 138.3, 137.3, 132.8, 132.5, 132.0, 132.0, 131.9, 131.6, 115.6, 114.7, 101.2, 97.4, 93.1, 68.5, 56.6, 54.3, 40.9, 29.2, 27.1; MS (ESI) m/z calculated for dimerized probe $C_{76}H_{78}N_6O_{18}S_2$ $[M + 2H]^{2+}$: 713.2402, found 713.2416.

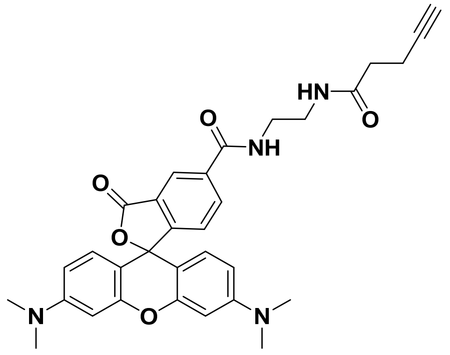


**16**

Compound **39** (43.0 mg, 0.10 mmol) was mixed with HATU (46.0 mg, 0.12 mmol) in 2 mL DMF, and the mixture was stirred for 10 min. N-(2-aminoethyl)pent-4-ynamide (17.0 mg, 0.12 mmol) was then added, followed by 53.0 μL of N,N-diisopropylethylamine (0.30 mmol). The reaction mixture was stirred for 3 h and was vacuum concentrated upon completion by TLC. The crude reaction mixture was purified by HPLC using water (0.1% TFA) as solvent A and acetonitrile (0.1% TFA) as solvent B. The flow at a rate of 10 mL/min used the program same as that used for purifying compound **15**. Compound **16** (38.6 mg, 70% yield) was eventually obtained as a pink solid with a retention time of 30 min. $^1$H NMR (500 MHz, $CD_3OD$): $\delta$ 8.67 (d, $J$ = 1.5 Hz, 1H),

8.15 (dd, $J$ = 8.0, 1.5 Hz, 1H), 7.43 (d, $J$ = 8.0 Hz, 1H), 7.04 (d, $J$ = 9.5 Hz, 2H), 6.96 (dd, $J$ = 9.5, 2.5 Hz, 2H), 6.88 (d, $J$ = 2.0 Hz, 2H), 3.50-3.47 (m, 2H), 3.42-3.38 (m, 2H), 3.21 (s, 12H), 2.40-2.36 (m, 2H), 2.35-2.31 (m, 2H), 2.16 (t, $J$ = 2.5 Hz, 1H); $^{13}$C NMR (126 MHz, $CD_3OD$): $\delta$ 174.7, 168.5, 167.5, 160.7, 159.1, 159.0, 138.1, 137.6, 133.1, 132.3, 131.94, 131.91, 131.3, 115.6, 114.7, 97.4, 83.5, 70.3, 41.3, 40.9, 39.9, 36.1, 15.7; MS (ESI) m/z calculated for $C_{32}H_{33}N_4O_5$ $[M + H]^+$: 553.2445, found 553.2446.

#### *Biological Experiments*

##### Antibody-Based Global Profiling of Acetylation

HEK293 cells (American Type Culture Collection) were maintained in 10 cm cell culture dishes with DMEM media and 10% FBS, 100 IU/mL penicillin, and 100 µg/mL streptomycin. Deacetylase inhibitor cocktail (100x, APExBIO) was added to the media overnight, after which the cells were washed with PBS and lysed in the CelLytic M buffer (Sigma Aldrich) that was pre-mixed with protease inhibitor cocktail (EDTA-free, Roche) and the deacetylase inhibitor cocktail (APExBIO). The cell mixture was sonicated for 18s (3s on, 7s off, 20% amplitude) on ice, and subsequently centrifuged at 15000 rpm at 4°C for 10 min. The resulting supernatants were collected and the protein concentration was determined by BCA assay (Pierce, Thermo Fisher) to be ~ 3 mg/mL. Approximately 50 µg of cell lysates were loaded for each lane and separated by 4-12% Bis-Tris SDS PAGE (Thermo Fisher), which were then transferred to PVDF membranes using a semi-dry blotting apparatus (Bio-Rad). Each lane on the membrane was carefully cut and blocked with 3% BSA in TBST (with 0.1% Tween-20) for 1 h. The piece of the membrane that contained one sample lane was then incubated with a specific anti-acetyl lysine antibody (from Ab21623, Ab190479, Ab80178, Ab61257, Abcam; or CST9441, CST9814, Cell Signaling Technology) overnight at 4 C, followed by washings and subsequent incubation with the IRDye® 680RD secondary antibody (Li-Cor). After extensive washing, protein bands were detected via near-infrared fluorescence on the LI-COR Odyssey FC Imaging System (700 nm channel scanning, Ex 685 nm/Em 730 nm).

##### Enzymatic Peptide Substrate Modifications with Fluorine

The PCAF assay cocktail was prepared by mixing 4 µL of 5x histone acetyltransferase (HAT) assay buffer (250 mM Tris-HCl, pH 8.0, 0.5 mM EDTA, 5 mM DTT), 1 µL of 2 mM histone H3-20 peptide (AnaSpec), 4.7 µL of 2.1 mM acetyl-CoA (Fisher Scientific) or acetyl CoA analogs, and 5.3 µL $H_2O$, in a total volume of 15 µL. After 5 µL of 1.3 µM PCAF enzyme (Cayman Chemical) was added, the reaction mixture was incubated at 30°C for 3h. The sample was then subjected to high resolution LC-MS analysis.

The MYST2 assay cocktail was prepared by mixing 4 μL of 5x HAT assay buffer, 0.5 μL of 2 mM histone H4-20 peptide solution (AnaSpec), 4.7 μL of 2.1 mM acetyl-CoA (Fisher Scientific) or acetyl CoA analogs, and 0.8 μL $H_2O$, at a total volume of 10 μL. After the addition of 10 μL 0.9 μM KAT7 enzyme (SignalChem), the reaction mixture was incubated at 30°C for 3h. The sample was then subjected to high resolution LC-MS analysis.

The TIP60 assay cocktail was prepared by mixing 0.5 μL of 2 mM histone H4-20 peptide (AnaSpec), 4.7 μL of 2.1 mM acetyl-CoA or related analogs, and 2.5 μL Tris buffer (50 mM, pH 8.0). Approximately 2.3 μL of 4.3 μM Tip60 (Cayman Chemical) in the stock buffer (50 mM Tris-HCl, pH 7.5, 100 mM NaCl, 10% glycerol) was then added and the reaction mixture was incubated at 30°C for 5h. The sample was finally analyzed by high resolution LC-MS.

##### Hydrolysis of Acetyl CoA and Fluoroacetyl CoA

The hydrolysis rate of acetyl-CoA and fluoroacetyl CoA were measured in a similar manner to reported procedures.[6-7] Briefly, 10 μM of the acetyl or fluoroacetyl CoA was incubated in 100 mM Tris buffer mixed with 0.5 mM DTNB, pH 7.2. The increase in the solution's absorbance at 412 nm was recorded, as a result of the reaction between DTNB and the free thiol in the released CoA hydrolysis product. A CoA standard curve was generated by measuring the absorbance of serially diluted CoA stock (200 μM) in the same assay buffer.

##### Non-Enzymatic Acetylation on Bovine Serum Albumin

Following the reported procedures for acyl-CoAs,[7] bovine serum albumin as a model protein (1 mg/mL) was dissolved in 50 mM HEPES, 150 mM NaCl, pH 8.0 or 7.0. Acetyl-CoA or fluoroacetyl CoA at the desired final concentrations, the buffer as a negative control, or the Sulfo-NHS-acetate (Pierce, Thermo Fisher) as a positive control were added to separate solutions, with the final pH adjusted to pH 8.0 or 7.0. Next, the reaction mixtures were incubated at 37 ºC for 6 h. The reaction samples (2 μL each) were then separated by SDS-PAGE, transferred to PVDF membrane, blocked and washed the same way as the aforementioned anti-acetylation western blot assays. The blot was probed with the MultiMab™ antibody (Ac-K-100, CST9814, Cell Signaling) that can recognize both acetyl-lysine and F-acetyl lysine.

##### Stability and Reactivity of Model Substrates and Probes in Cell Lysates

The cell lysate experiments were performed similarly to those reported for other previously established bio-orthogonal reactions.[8] In general, 5 μL of substrate compound 3, 3-Cl, or the probe 13 mixed with their corresponding internal standards (3' or 13') in DMSO (10 mM stock concentration) were added to 25 μL HEK293 cell lysates (~ 3 mg/mL protein concentration). TCEP (5 μL, 60 mM stock concentration) was added to each reaction group (A, B, or C, respectively, Figure S10A) to maintain a reducing environment. The final reaction pH was adjusted to 8.5, and water was added to make a final volume of 50 μL. After incubation at 37 °C for 14h, the reaction mixture's pH was adjusted to ~ 6.

The solution was extracted with ethyl acetate three times and concentrated in vacuo. The resulting residues were redissolved in methanol and analyzed by high resolution LC-MS (Wistar Institute) on a ThermoFisher Scientific Q Exactive HF-X mass spectrometer equipped with a HESIII probe and coupled to a ThermoFisher Scientific Vanquish Horizon UHPLC system. Compounds were separated on a Synergi™ Polar RP column (4 μm, 150 x 1 mm, Phenomenex). After LC-MS analysis, the peak area for each compound was integrated, and the percent recovery yield was calculated versus the internal standard. The FTDR reaction in cell lysate (Figure S11) was carried out using the same procedure except that the incubation time was 5 h, and the extracts were analyzed by LC-MS/MS analysis.

**Biotinylation of Labelled Peptides based on the Fluorine-Thiol Displacement Reaction**

The histone peptide H3-20 that previously underwent fluoroacetylation was lyophilized and resuspended in water (~ 400 μM). About 1 μL of this stock solution was mixed with 1 μL Tris buffer (1M) and 4 μL $H_2O$. Then, 1 μL of the Biotin-SH probe stock in DMF (40 mM) was added, along with 1 μL of 50 mM TCEP aqueous solution. After adjusting the pH to 8.5 (with 2 μL of 1 M NaOH solution), the final concentrations of H3-20 substrate, Biotin-SH probe, and TCEP were 40 μM, 4 mM, and 5 mM, respectively. The reaction was incubated at 37 °C overnight, and the sample was subjected to high resolution LC-MS analysis.

**Fluorination and Biotinylation of Known Protein Substrates by Acetyltransferase Assay and FTDR**

The protein substrates of acetylation such as human histone H3.1 (New England Biolabs) or histone H4 (New England Biolabs) was dissolved in the aforementioned HAT buffer at a final concentration of 15 μM, and was mixed with acetyl-CoA analogs (450 μM) and the corresponding acetyl transferase (1.3 μM) (PCAF (Cayman Chemical) for histone H3.1; MYST2 (SignalChem) for histone H4). The reaction mixture was incubated at 30 °C, pH 7.2 for 6h, and was quenched by lyophilization.

For biotinylation, the fluorinated protein mixture was redissolved in water (final concentration ~ 35 μM), added with 1.75 mM Biotin-SH probe and 2 mM TCEP. The pH was adjusted to 8.5 and the mixture was incubated at 37 °C for indicated time period (1 h, 3 h, or 6 h). Samples after this reaction were then loaded on gels for SDS-PAGE analysis. The control protein mixture (previously treated with 4PY-CoA) was reacted with biotin-azide probe (Thermo Fisher) following the reported copper-catalyzed azide-alkyne cycloaddition (CuAAC) procedures,[4] and was analyzed by SDS-PAGE in parallel to other samples.

For in-gel fluorescent imaging, the separated proteins on the PAGE gels were fixed with 50% isopropanol/45% water/5% acetic acid for 15 min at RT. The gel was washed, and incubated for

1 h at RT in biotin-free Casein blocking buffer (Sigma Aldrich). The blocked gel was then probed with streptavidin-IRDye 680RD (LI-COR) at a dilution of 1/2000 for 45 min at RT, followed by multiple washings with PBST (containing 0.1% Tween-20). Those biotinylated proteins were finally visualized by near-infrared fluorescence detection through the LI-COR Odyssey FC Imaging System (700 nm channel, Ex 685 nm/Em 730 nm).

**Removal of Fluoroacetylation on Protein Substrates by Histone Deacetylases**

A 5x histone deacetylase assay buffer was prepared, containing 250 mM Tris-HCl (pH 8.0), 685 mM NaCl, 5 mM $MgCl_2$, 13.5 mM KCl, and 5 mM DTT. The fluoroacetylated histone substrate (~ 20 µM final concentration) was mixed with the corresponding histone deacetylase (SIRT1 (R&D Systems) for fluoroacetylated H3; HDAC1, HDAC2, HDAC3, or SIRT2 (BPS Bioscience) for fluoroacetylated H4), and 2 µL of the 5x assay buffer. For assays involving sirtuins, nicotinamide adenine dinucleotide (NAD+, Fisher Scientific) was added at a final concentration of 1 mM. The reaction mixture's pH was adjusted to 8.0, with water added to make a final volume of 10 µL. The mixture was incubated at 37 ºC for 6h, and subsequently treated with Biotin-SH to initiate FTDR reaction on any remaining fluoroacetylation. All the samples were finally separated on SDS-PAGE, transferred to PVDF membranes, and probed by streptavidin-IRDye 680RD (LI-COR) as mentioned before.

**Expression, Purification and Characterization of EZH2 Protein**

The plasmid expressing GST-fused EZH2 N-terminal domain (1-500) was constructed based on the parent plasmid pGEX-EZH2 that was a gift from Prof. Min-Chie Hung (Addgene plasmid # 28060).[9] The C-terminal sequence (501-745) was deleted using Q5® Site-Directed Mutagenesis Kit (New England Biolabs) and customized primers (Integrated DNA Technologies). The sequences of the resulting pGEX expression vector were confirmed by DNA sequencing (GENEWIZ). The final vector was transformed into BL21 competent cells via electroporation. The resulting cells were recovered in SOC medium and plated onto an ampicillin-containing LB agar plate. After overnight incubation at 37 ºC, the colonies were picked, amplified and mixed with 50% glycerol as cell stock for EZH2 (1-500) protein expression. On the day of expression, BL21 stock was inoculated in LB medium supplemented with 100 µg/mL ampicillin, and shaken at 250 rpm, 37 ºC (MaxQ 8000, Thermo Scientific). When $OD_{600}$ reached 0.8, 0.5 mM IPTG was added to induce protein expression. The cell culture was left for ~ 12 h at 25 °C, 250 rpm (MaxQ 8000, Thermo Scientific). Cells were then harvested and frozen at -80 °C overnight.

The bacteria pellets were lysed in lysis buffer (50 mM Tris-HCl, pH 8.0, 150 mM NaCl, 2 mM EDTA, 20% sucrose, 0.2% TritonX-100, and 5% glycerol) that also contained 1 mg/mL lysozyme (VWR) and protease inhibitor cocktail (Roche cOmplete). After shaking at 250 rpm, 25 ºC for 1.5 h, the cell lysate were centrifuged at 9000 rpm for 40 min (FX6100 Rotor, Beckman Coulter), and was filtered through a 0.2 µm filter to remove debris. The filtrate was passed through affinity columns packed with glutathione resin (GenScript), which was later washed with 20 mL PBS buffer (0.02% tween-20). Finally, 15 mL of elution buffer (10 mM glutathione, 50 mM Tris-HCl, pH 8.0) was added, and the eluted EZH2 (1-500) protein was buffer exchanged into storage buffer (50 mM Tris-HCl, pH 8.0, 10% glycerol, 1 mM DTT, 0.1 mM EDTA) by Amicon® Ultra Centrifugal Filters (MilliporeSigma). The protein purity and identity was confirmed by SDS-PAGE.

The fluorination and biotinylation on EZH2 protein were performed following the aforementioned procedures (acetyltransferase assay and FTDR using Biotin-SH probe) for histone substrates. The acetyltransferase used for EZH2 was PCAF (Cayman Chemical). To prepare the positive control for CuAAC ("C"), azidoacetic acid NHS ester ($N_3$-NHS) was synthesized according to published procedures.[10] Briefly, 30 µL of EZH2 protein (0.65 mg/mL in DPBS with 0.25 mM TCEP added, pH 7.2) was first reacted with 2.5 µL of $N_3$-NHS (1mM, DPBS buffer, pH 7.2) at room temperature for 1h. To the resulting EZH2 mixture with lysines conjugated by azide, 1 µL of THPTA (10 mM), 1 µL of sodium ascorbate (50 mM), 0.5 µL of biotin-alkyne linker (5 mM), and 1 µL of $CuSO_4$ (5 mM) were added to initiate the CuAAC-based modification of EZH2 by biotin. The reaction mixture was incubated at room temperature for 2h, before loading to SDS PAGE for analysis in parallel to other samples.

**Cell Cytotoxicity Studies of Cofactor Analogs**

HeLa cells were obtained from American Type Culture Collection (ATCC), and were cultured similarly to the aforementioned HEK293 cell line in DMEM medium supplemented with 10% FBS, 100 IU/mL penicillin, and 100 µg/mL streptomycin. Both cells were maintained in a cell-culture incubator at 37 ºC, with 5% $CO_2$. On the night before the assay, these cell lines were plated into 96-well cell culture plates (white, flat bottom, Costar) at 5,000 cells, 90 µL media per well. The next day, cofactor analogs (azido-ethyl acetate, fluoro-ethyl acetate, or DMSO carrier control) were serially diluted in culture media as 10x stock solutions, and then added to the pre-plated cells at 10 µL/well. The final treatment concentrations are 62.5, 125, 250, 500, 1000, and 2000 µM. After thorough mixing, the samples were incubated in the cell culture incubator for 12h at 37 ºC, under 5% $CO_2$. At the end of the treatment, the plates were taken out and cooled down to room temperature. Cell viability was measured by CellTiter Glo assay (Promega) following the published procedure.[11] Data were processed and plotted using Graphpad Prism (GraphPad Software). Viability of the cells treated with DMSO control was adopted as 100% viability control.

**Cellular Metabolism Study of Ethyl Fluoroacetate**

HEK293 cells were cultured in a 100 mm x 20 mm tissue culture dish (Corning Costar) up to ~ 80% confluence. The pro-metabolite ethyl fluoroacetate was added at a final concentration of 1 mM, and the culture was incubated at 37 ºC for 2 h. The acyl-CoA extraction was performed following the reported procedures.[12] Briefly, the cells were gently scraped down into the media, and spun down at 1,000 g. The cells were resuspended in 1 mL ice-cold extraction solution (10% trichloroacetic acid in milliQ water), and sonicated on ice for 30s (1 pulse per second). The resulting cell lysate was centrifuged at 15,000 g, 4 ºC for 5 min to precipitate the debris. In the meanwhile, Oasis HLB SPE columns (Waters) were conditioned by 1 mL methanol and equilibrated with 1 mL water. The supernatant of the cell lysate was loaded to the column, with subsequent washing of the column with water. The potential CoA extracts were eluted by 0.5 mL elution solution (25 mM ammonium acetate in methanol), dried in vacuo, and resuspended in 50 uL 5% 5-sulfosalicyilic acid for LC-MS/MS analysis. As a control, the HEK293 cell lysate was heated at 90 ºC for 10 min, and then mixed with 1 mM ethyl fluoroacetate at 37 ºC for 2 h. The follow-up extraction and LC-MS/MS analysis were carried out the same as mentioned above.

**Intracellular Fluorination of Protein Substrates**

HeLa or HEK293 cells were seeded at a density of 20,000 cells/well in a 12-well flat bottom cell culture plate (Corning Costar). When they reached ~ 80% confluence, cells were incubated with 1 mM ethyl azidoacetate (for "click chemistry"), ethyl fluoroacetate (for fluorination), or DMSO control at 37 ºC in standard medium for 6 h, similar to reported procedures.[13] The cells were then ready for follow-up intracellular imaging.

**Intracellular Dye Labeling and Imaging of Protein Substrates**

After incubation with azido- or fluoro-modified ethyl acetate precursors, cells were rinsed for three times with DPBS buffer, and fixed with 3.2% paraformaldehyde for 10 min at RT. With another around of rinses, cells were permeabilized with 0.1% Triton-100 in PBS buffer for 10 min at RT, after which point the intracellular proteins became ready for labeling by TAMRA dyes.

The "click chemistry" labeling on control groups was performed according to published procedures.[14] Cells previously treated with azidoacetate or DMSO were incubated with 100 μM TAMRA-alkyne probe (compound **16**) in the standard medium that also contained 200 μM $CuSO_4$, 500 μM BTTES, and 2.5 mM freshly prepared sodium ascorbate. The mixture was reacted at 37 ºC for 1 h, followed by PBS washing, and a subsequent nucleus staining with 1 μg/mL Hoechst 33342 (Fisher Scientific). For fluorine-thiol displacement labeling, cells previously treated with fluoroacetate or DMSO were washed twice with PBS buffer (pH 8.0) that contained 5 mM TCEP. Each round of washing consisted of a 10 min incubation period. The cells were then incubated with 1 mM TAMRA-SH probe (compound **15**), and 5 mM TCEP (pH 8.5) in the standard medium to ensure the complete displacement of fluorinated intracellular proteins. After an incubation of 6 h at 37 ºC, cell samples were washed with TCEP containing PBS buffer for three times, and then stained with Hoechst 33342 dye at RT for 10min. All cell samples were briefly washed after nucleus staining, and examined using the ZOE fluorescent microscope imager (Bio-Rad Laboratories).

**In-Gel Fluorescent Imaging of Protein Substrates Labelled from Cell Lysates**

HEK293 cells were seeded at 20,000 cells/dish in a 100 mm x 20 mm cell culture dish (Corning). Upon ~ 80% confluence, cells were incubated with 1 mM ethyl azidoacetate (for "click chemistry"), ethyl fluoroacetate (for fluorination), or DMSO as a control at 37 ºC for 6 h. For

HAT inhibition assay, cells were preincubated with 10 μM anacardic acid and 10 μM MG149 (Selleckchem) for 6h before the addition of pro-metabolite ethyl fluoroacetate for fluorination. After these treatments, cells were rinsed three times with DPBS buffer. Each cell sample was immediately lysed, centrifuged, with supernatants collected and the protein concentration measured as mentioned before.

For "click chemistry" labeling of the azido-modified proteins in cell lysates, approximately 300 μL supernatant of each cell lysate (~ 2 mg/mL) was incubated with 1 mM TCEP, 100 μM TBTA, 1 mM $CuSO_4$, and 100 μM TAMRA-alkyne probe (compound **16**) according to published procedures.[13, 15] To prepare the positive control ("C") for "click chemistry" labeling, BSA control protein (3 mg/mL in PBS, pH 8.0) was mixed with 30 equivalents of the aforementioned N3-NHS ester. The reaction mixture was rotated at room temperature for 4h, and the BSA protein was purified by methanol precipitation twice. The protein pellet was redissolved in PBS buffer and modified with TAMRA-alkyne probe (**16**) following the similar CuAAC reaction conditions as mentioned above. For fluorine-thiol displacement labeling, approximately 300 μL supernatant of each cell lysate (~ 2 mg/mL) was treated with 5 mM TCEP and 2 mM TAMRA-SH probe (compound **15**) at pH 8.5 for 6 h to ensure the complete displacement. All the protein samples after reaction each had 1 mL of cold acetone added to precipitate out proteins. Proteins were then resuspended in PBS buffer, re-precipitated with methanol, and pelleted by centrifuging at 15,000g, 4 °C for 10 min. Each protein pellet was redissolved in 100 μL PBS buffer that comprised of 1% SDS and 10% glycerol. Approximately ~ 30 μg of each protein sample was mixed with SDS-PAGE sample buffer, and separated by 4%-12% gradient SDS-PAGE analysis. Approximately 1 μg of the BSA positive control "C" was loaded in parallel. In-gel fluorescence detection of the labelled TAMRA dye was achieved by scanning the gel with the LI-COR Odyssey FC Imaging System (700 nm channel, Ex 685 nm/Em 730 nm).

**FTDR-Based Labeling and Imaging of Protein Substrates with Concurrent HDAC Inhibition**

HEK293 cells were cultured and treated with the pro-metabolite ethyl fluoroacetate, or DMSO control as mentioned above. To probe the effect of HDAC inhibition, 1x deacetylase inhibitor cocktail (APExBIO) was added to the medium concurrently with the addition of the pro-metabolite and was incubated for the indicated time (6h and 12h, respectively). The cell samples were then lysed, with the proteins labelled by the TAMRA-SH probe based on FTDR. Subsequently, each sample was purified, loaded to SDS-PAGE, and imaged the same as mentioned above.

**Histone Extraction**

HEK293 cell samples were prepared as mentioned above. For the competition assay, 10 mM ethyl acetate was added to the media concurrently with the pro-metabolite ethyl fluoroacetate. After cell lysis and FTDR, histones were extracted with the EpiQuik Total Histone Extraction kit (Epigentek) following the manufacturer's instructions. Approximately 3 μg of each histone extract was mixed with the sample loading buffer and separated on 12% SDS-PAGE, followed by subsequent in-gel fluorescent imaging and CBB staining as mentioned before.

**FTDR-Based Pull Down of Protein Substrates Followed by Western Blot Analysis**

HEK293 cells after step 1 treatment (Figure 5A) was lysed and the resulting proteome (~ 2 mg/mL, pH 8.5) was added with 100 mM TCEP, 4 mM Biotin-SH probe (compound 14). The reaction mixture was incubated at 37℃ for 6h, with the unconjugated probes removed by subsequent methanol precipitation twice. The protein pellet was redissovled in PBS buffer (0.1% SDS, pH 7.2). Approximately 100 μL of the proteins (1 mg/ml) were mixed with 100 μL streptavidin magnetic beads (New England BioLabs) at room temperature for 1h. The beads were sequentially washed with PBS buffer (0.1% SDS, pH 7.2), PBS buffer (0.2% SDS and 4M urea, pH 7.2), and PBS buffer (0.2% SDS, pH 7.2). After the final washing with pure PBS buffer three times, the beads were incubated with the elution buffer (10 mM sodium periodate in PBS) for 30 min in the dark. The elution was repeated three times and the combined elute was concentrated on a lyophilizer. The eluted protein samples were finally separated by 4-12% Bis-Tris SDS-PAGE and transferred onto PVDF membranes. The blot was blocked with 3% BSA in TBST for 1h, followed by the incubation with the anti-Histone H3 antibody (HRP Conjugate, CST#12648), anti-Histone H4 antibody (HRP Conjugate, abcam#ab197517), and the anti-alpha-Tubulin antibody (CST#2144) at 4℃. After overnight incubation, the blot was washed with TBST for three times and was further incubated with an anti-rabbit secondary antibody (HRP conjugated) for the detection of alpha tubulin. After 30 min incubation at room temperature, the blot was washed with TBST and imaged using the Clarity Max Western ECL substrate.

**Proteomics Study of Fluoroacetylated Histones and Alpha-tubulin**

HEK293 cells were treated with ethyl fluoroacetate (Figure 6A), with subsequent histone extraction carried out the same as mentioned above. The histone proteins were desalted using the C4 columns (The Nest Group), lyophilized, and redissolved in the incubation buffer (50 mM Tris-HCl, 5 mM $CaCl_2$, 2 mM EDTA, pH 7.6-7.9) for in-solution digestion. After the addition of Arg-C (Promega), the digestion was activated with the 10x activation buffer (50 mM Tris-HCl, 60 mM DTT, 2 mM EDTA, pH 7.6-7.9) that was added to a final concentration of 1x. The reaction mixture was incubated at 37 ℃ for 7 h, followed by purification using C18 columns (The Nest Group). The elusions are collected, lyophilized, and redissolved in water to be injected into LC-MS/MS system. For immuno-enrichment of endogenous alpha tubulin, 100 μL of each cell lysate was gently mixed with 10 μL of anti-alpha-tubulin antibody (Cell Signaling, CST3873S) and incubated overnight at 4 ℃. Approximately 50 μL of Dynabeads Protein G (Invitrogen) was then added and the mixture was incubated at 4 ℃ for 1 h. The Dynabeads were pulled down and washed with 300 μL PBS buffer (0.05% Tween-20) three times. The targeted proteins were eluted by heating the beads at 90 ℃ for 10 min within the LDS samples loading buffer (GenScript). The resulting supernatants were loaded onto a 10% SurePAGE Bis-Tris gel (GenScript). After staining by Coomassie brilliant blue R-250, the bands at 50 kDa were cut off for in-gel digestion by Glu-C

(Promega). Liquid chromatography tandem mass spectrometry (LC-MS/MS) analysis was performed as previously described[16] using a Q Exactive HF mass spectrometer (ThermoFisher Scientific) coupled with a Nano-ACQUITY UPLC system (Waters). Peptide sequences were identified using MaxQuant v1.6.15.0.[17] MS/MS spectra were searched against a UniProt human protein database (10/10/2019) using full enzyme specificity with up to two missed cleavages, static carboxamidomethylation of Cys, and variable Met oxidation (+15.9949 Da), Asn deamidation (0.9840 Da), Lys acetylation (+42.0106 Da) and Lys F-acetylation (+60.0011 Da). Consensus identification lists were generated with false discovery rates of 1% at protein, peptide and site levels.

### Supplemental Figures

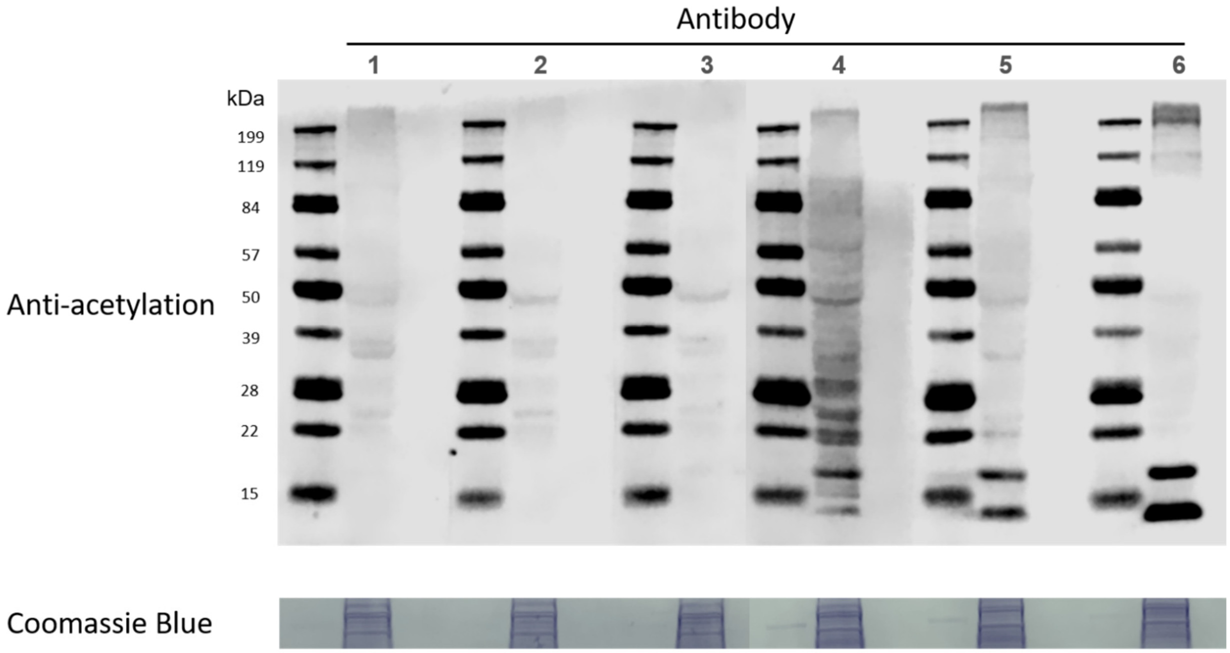


**Figure S1.** Global profiling of lysine acetylation using anti-acetyl lysine antibodies that are commercially available from different vendors (#1-6). Top panel: Western blots of lysine acetylation in HEK 293 cell lysates with different anti-acetyl lysine antibodies (#1-6); Bottom panel: Coomassie blue staining of the 120 kDa area as loading controls.

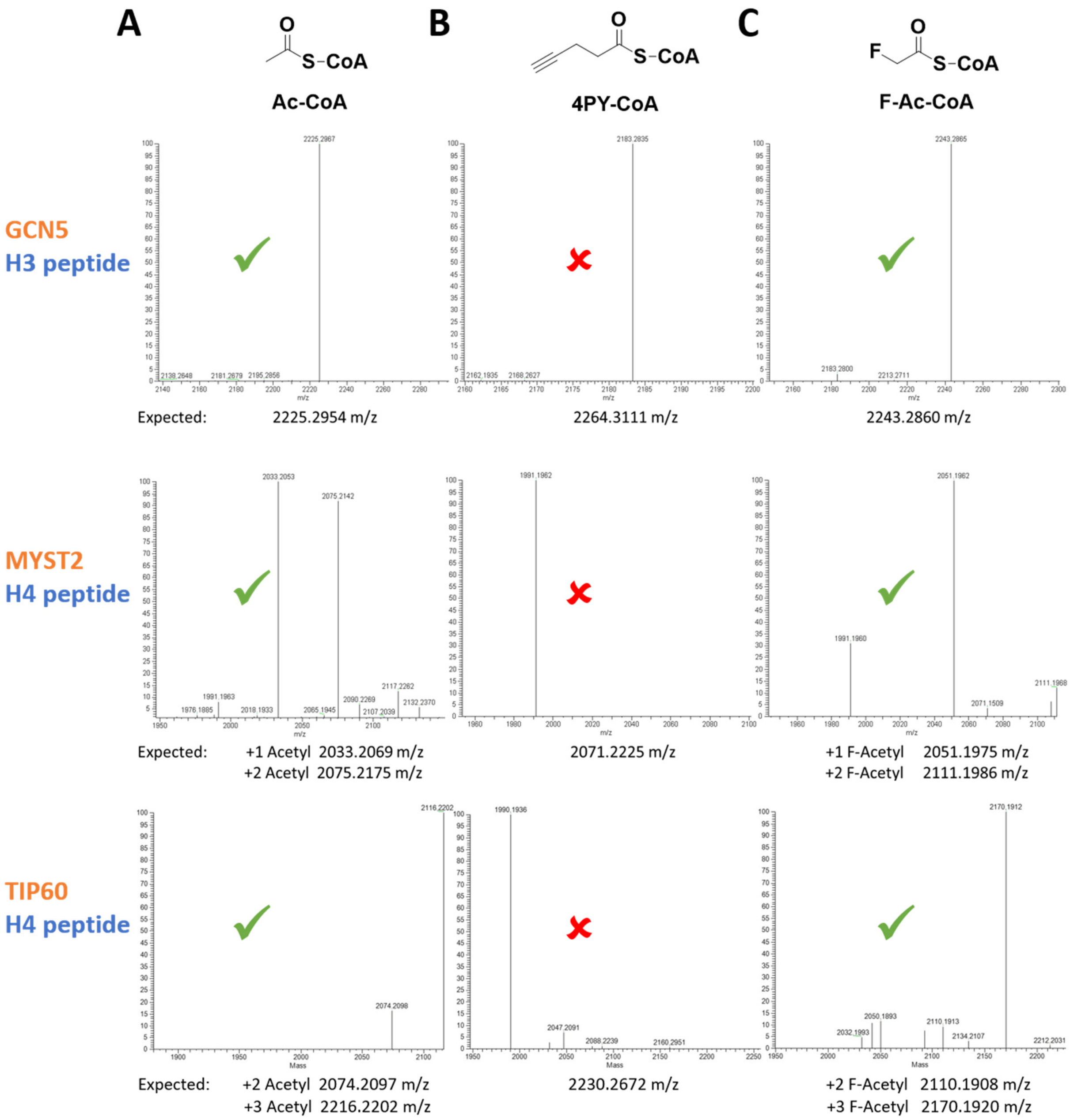


**Figure S2.** ESI-MS results for using acetyl-CoA analogs to label KAT substrates. (A) Acetyl-CoA, (B) 4PY-CoA, and (C) F-Ac-CoA were each mixed with corresponding acetyl transferases (GCN5, MYST2, or TIP60) and peptide substrates (H3-20: N-terminal 20-aa H3 peptide, exact mass 2182.2771 m/z; or H4-20: N-terminal 20-aa H4 peptide, exact mass 1990.1885 m/z). Top row of MS spectra: results with GCN5 KAT and H3 (1-20) peptide. Middle row: results with MYST2 KAT and H4 (1-20) peptide. Bottom row: results with TIP60 KAT and H4 (1-20) peptide. Expected theoretical m/z were shown under each MS spectrum. "√" indicates the observed m/z matches to the expected values; "×" indicates not, which is the case for the assays with 4PY-CoA that only resulted in unmodified wild type substrates.

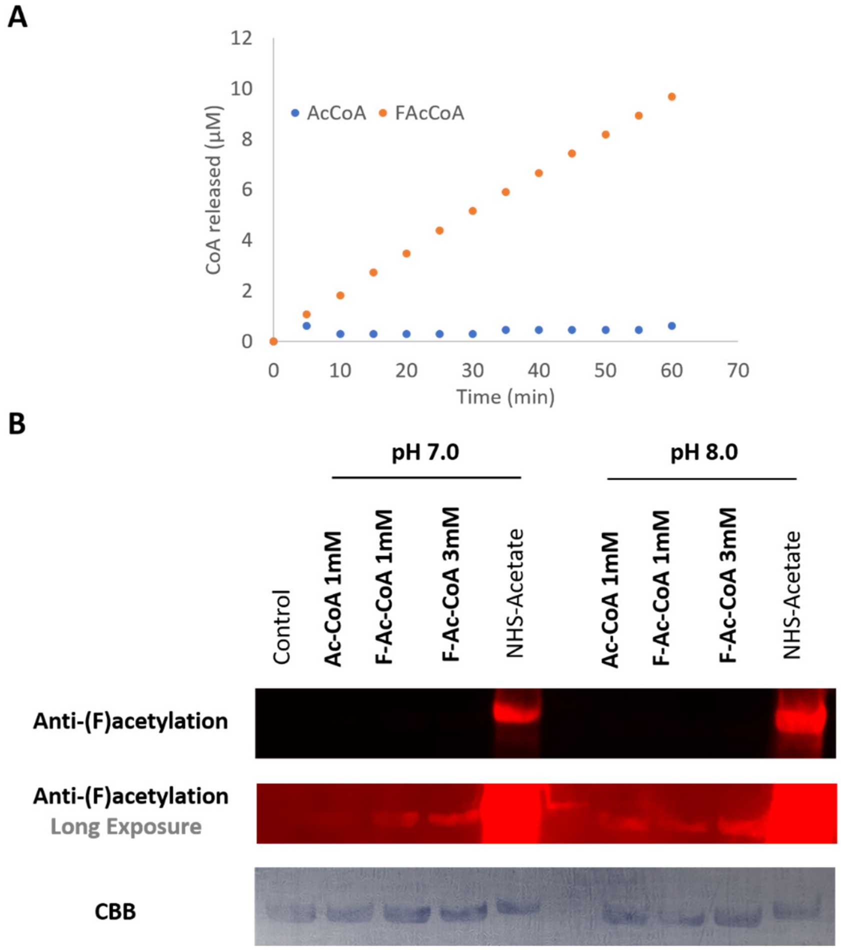


**Figure S3.** Chemical reactivity of fluoroacetyl-CoA (F-Ac-CoA) and acetyl-CoA (Ac-CoA). (A) Comparison of rates of hydrolysis of 10 μM F-Ac-CoA and 10 μM Ac-CoA at 100 mM Tris buffer (pH 7.2); The pseudo-first-order rate constants were determined to be 7.5 x $10^{-3}$ $s^{-1}$ for F-Ac-CoA, and 2.0 x $10^{-4}$ $s^{-1}$ for Ac-CoA, respectively. (B) The non-enzymatic acetylation of F-Ac-CoA and Ac-CoA to the model protein bovine serum albumin (BSA) at pH 7.0 or pH 8.0, 37ºC; The positive control, NHS-acetate, was tested in parallel, which was supposed to readily modify lysines of BSA with acetates. Top row: western blot for acetyl-lysine or F-acetyl lysine residues on BSA; Middle row: long exposure of the western blot; Bottom row: coomassie brilliant blue (CBB) staining as loading controls. The MultiMab™ antibody (Ac-K-100, Cell Signaling) used for these western blots can recognize both acetyl-lysine and F-acetyl lysine.

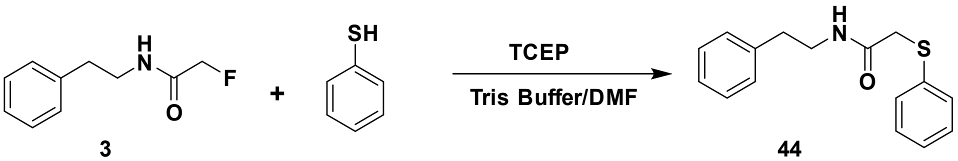


3 44

A.

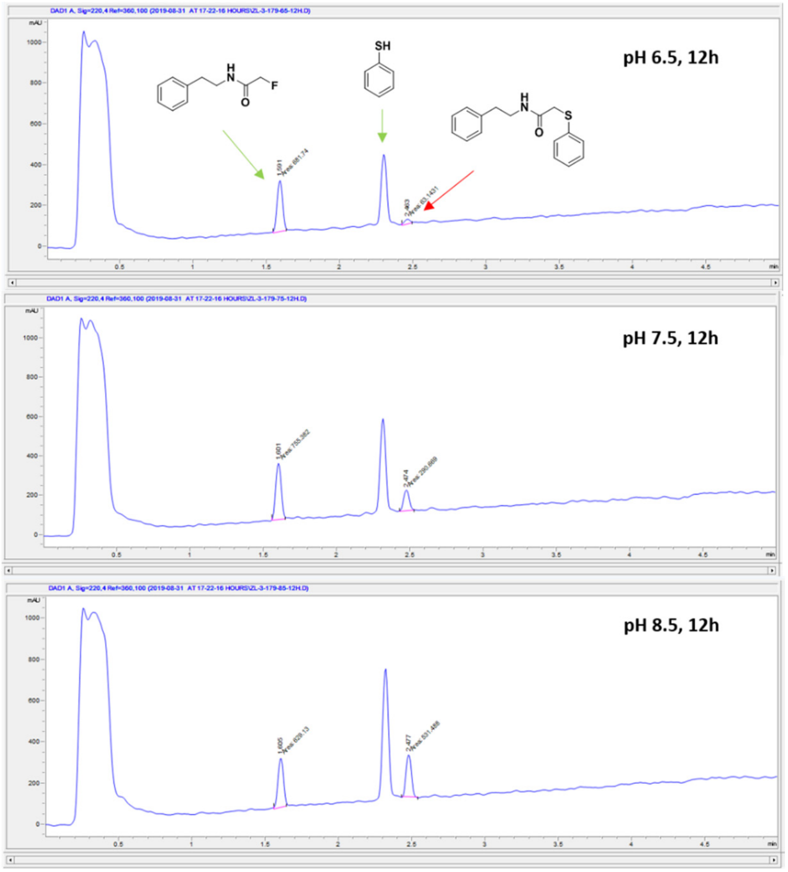


B.

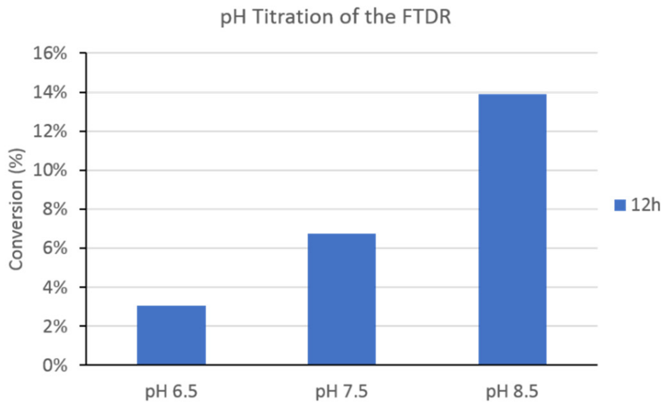


**Figure S4.** Optimization of the fluorine-thiol displacement reaction (FTDR) by titrating the effects of reaction pH values. Standard nucleophile (thiophenol) and substrate **3** (2-Fluoro-N-phenethylacetamide) were used. (A) Representative LC-MS spectra of reaction mixtures after 12 h of reaction under varied pH's. (B) Summary plot of the product yields at different reaction pH's.

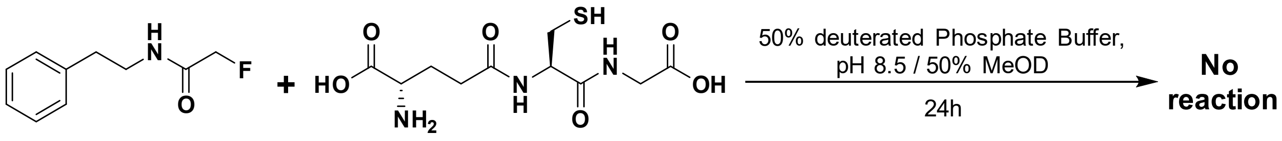


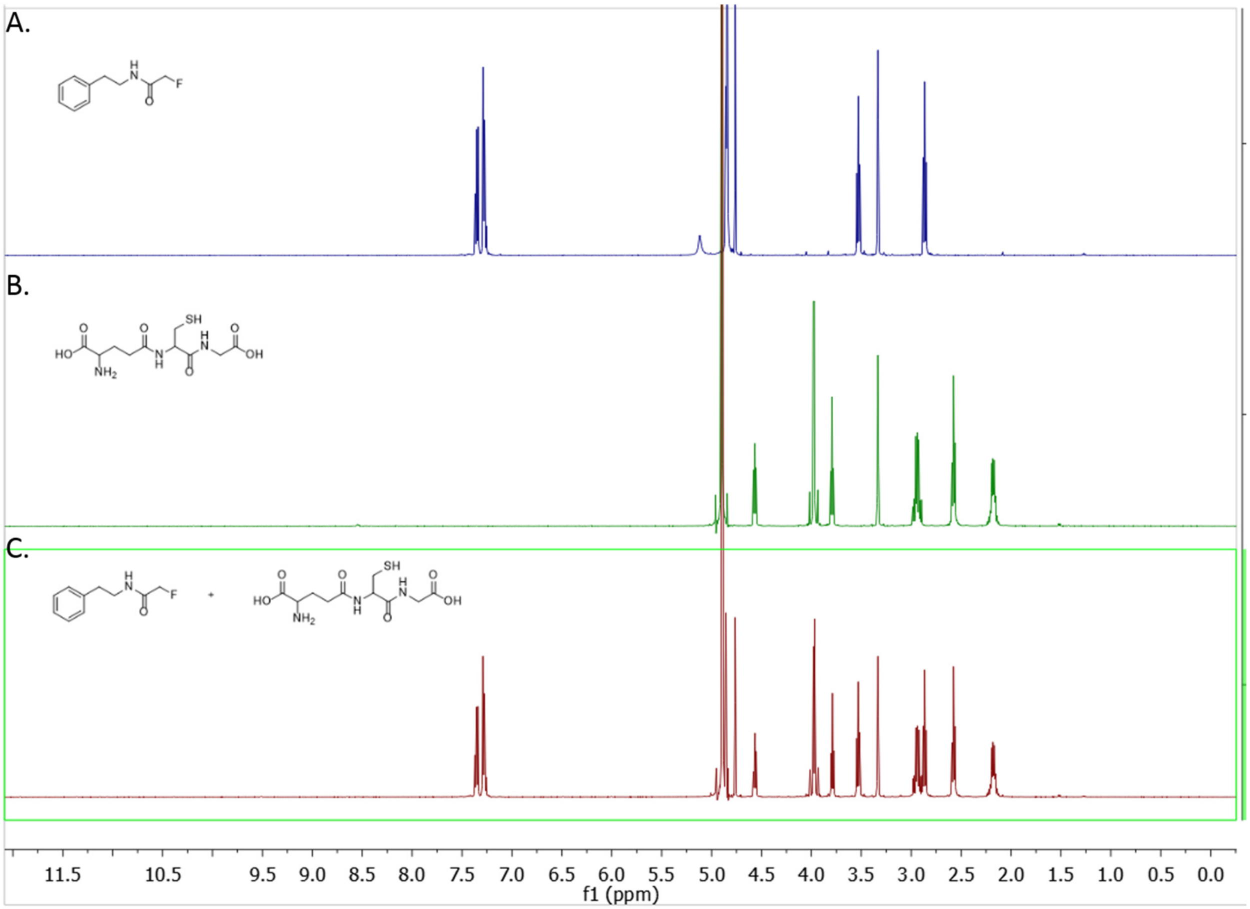


**Figure S5.** Substrate **3** (2-Fluoro-N-phenethylacetamide) is stable upon incubation with glutathione. (A) $^1$H-NMR spectra of **3** (25 mM in 1:1 mix of deuterated sodium phosphate buffer and MeOD, pH 8.5). (B) $^1$H-NMR spectra of reduced glutathione (GSH) (25 mM in 1:1 mix of deuterated sodium phosphate buffer and MeOD, pH 8.5). (C) $^1$H-NMR spectra of the mixture of **3** (25 mM) and GSH (25 mM) in the 1:1 mixed deuterated sodium phosphate buffer and MeOD, pH 8.5, after 24 h of incubation at 37 ºC under the nitrogen atmosphere.

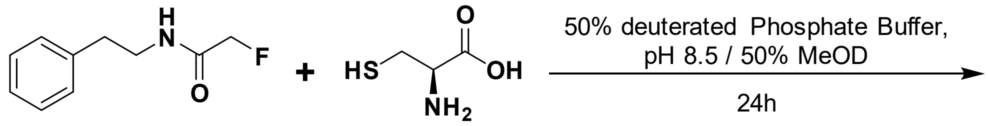


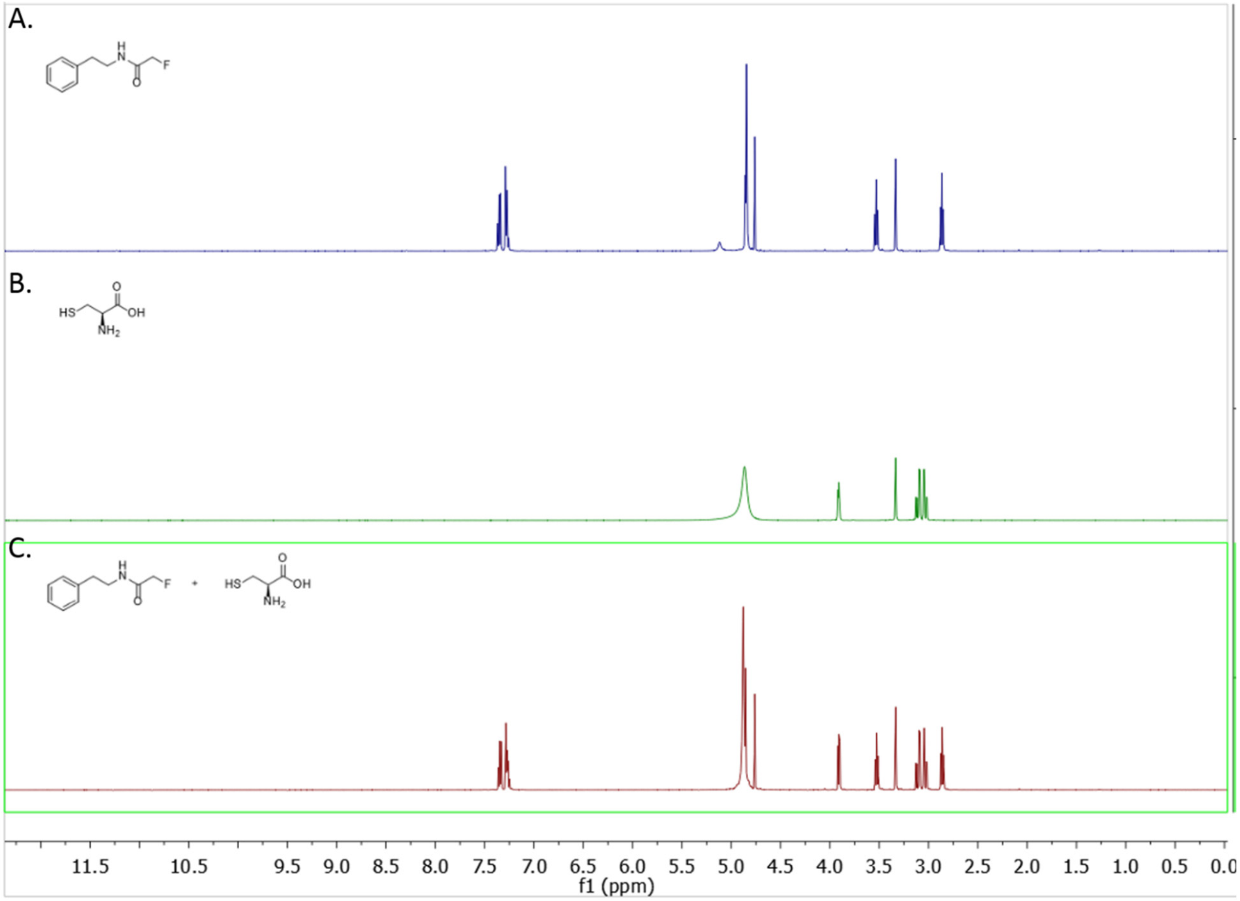


**Figure S6.** Substrate **3** (2-Fluoro-N-phenethylacetamide) is stable upon incubation with cysteine. (A) $^1$H-NMR spectra of **3** (25 mM in 1:1 mix of deuterated sodium phosphate buffer and MeOD, pH 8.5). (B) $^1$H-NMR spectra of reduced glutathione (GSH) (25 mM in 1:1 mix of deuterated sodium phosphate buffer and MeOD, pH 8.5). (C) $^1$H-NMR spectra of the mixture of **3** (25 mM) and GSH (25 mM) in the 1:1 mixed deuterated sodium phosphate buffer and MeOD, pH 8.5, after 24 h of incubation at 37 ºC under the nitrogen atmosphere.

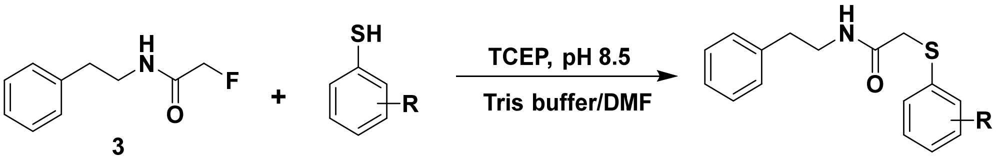


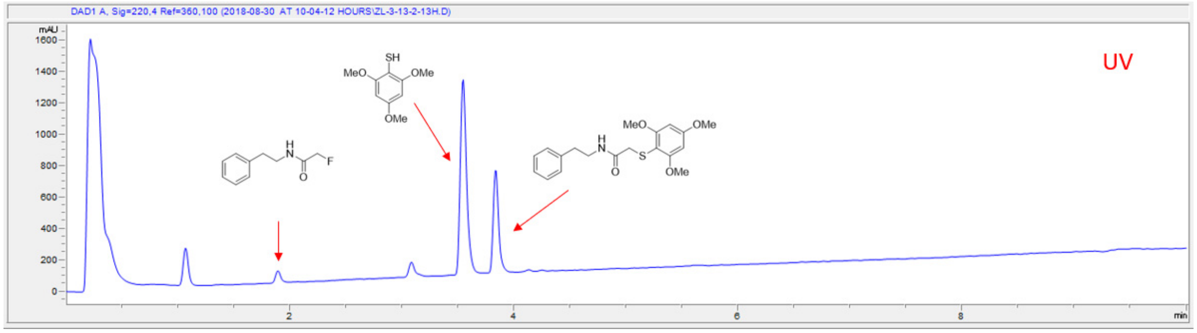


**Figure S7.** Representative LC-MS spectra for analysis of the reaction mixtures undergoing general procedure B. The UV trace was for the reaction between substrate **3** and the nucleophile 2,4,6-trimethoxybenzenethiol (**13**) after 13 h of incubation at 37 °C. Identities of the peaks were confirmed by the corresponding ESI-MS analysis.

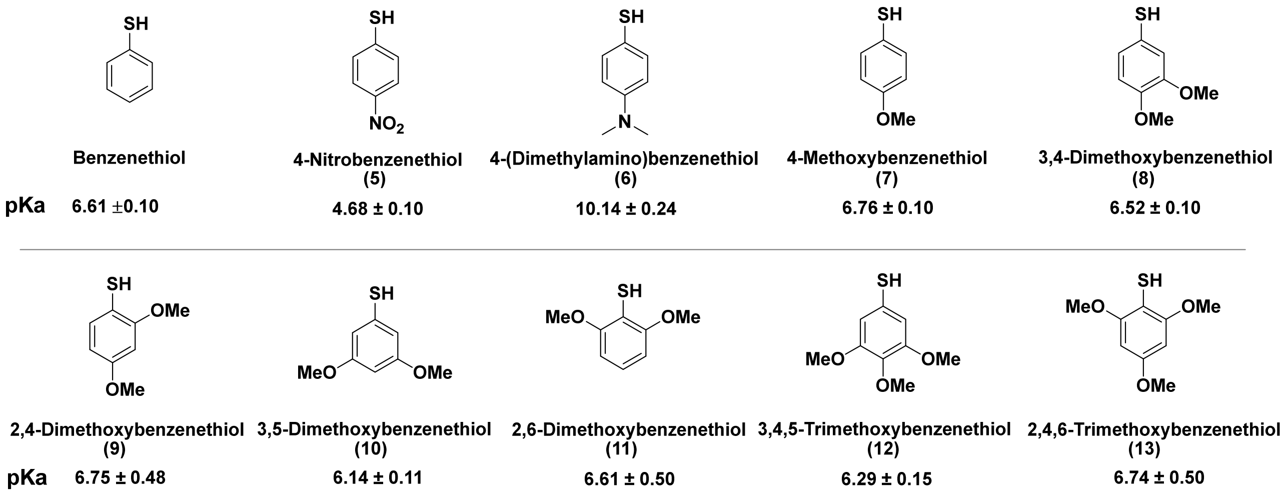


**Figure S8.** Summary of the pKa values of the benzenethiol derivatives. The data was referred from the SciFinder database.

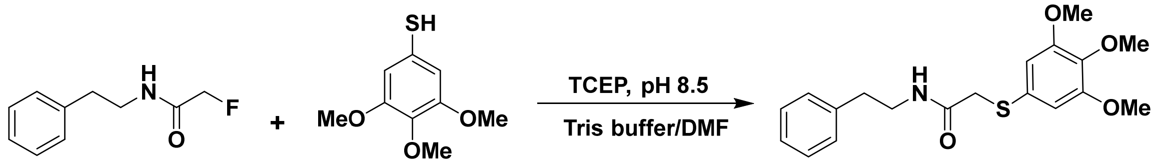


3 12 46

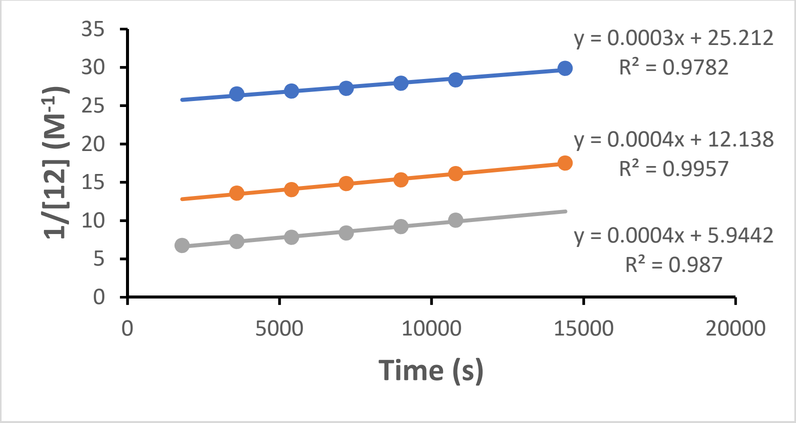


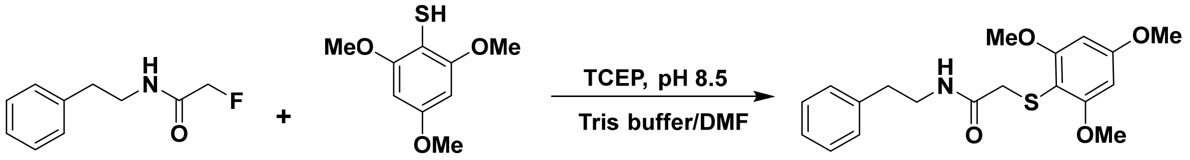


3 13 47

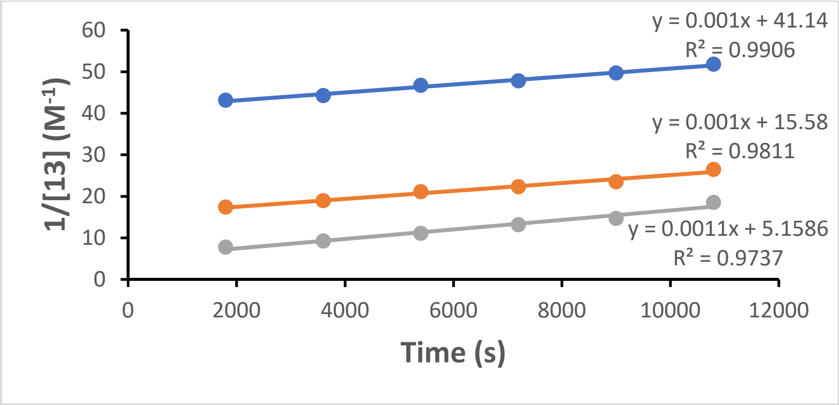


**Figure S9.** Evaluating the second order rates of FTDR reactions between **3** and **12**, **3** and **13**, respectively. Equal concentrations of both reactants were used and the assays were repeated independently at three different concentrations (40 mM, 80 mM, and 160 mM). Plotting $1/[X]_t$ (concentration of either reactant at time t) against time yielded the rate constant ((0.37 ± 0.06) x $10^{-3}$ $M^{-1}$ $s^{-1}$ for **3** and **12**, (1.03 ± 0.06) x $10^{-3}$ $M^{-1}$ $s^{-1}$ for **3** and **13**). The reported values represented an average of the three independent experiments as mentioned above.

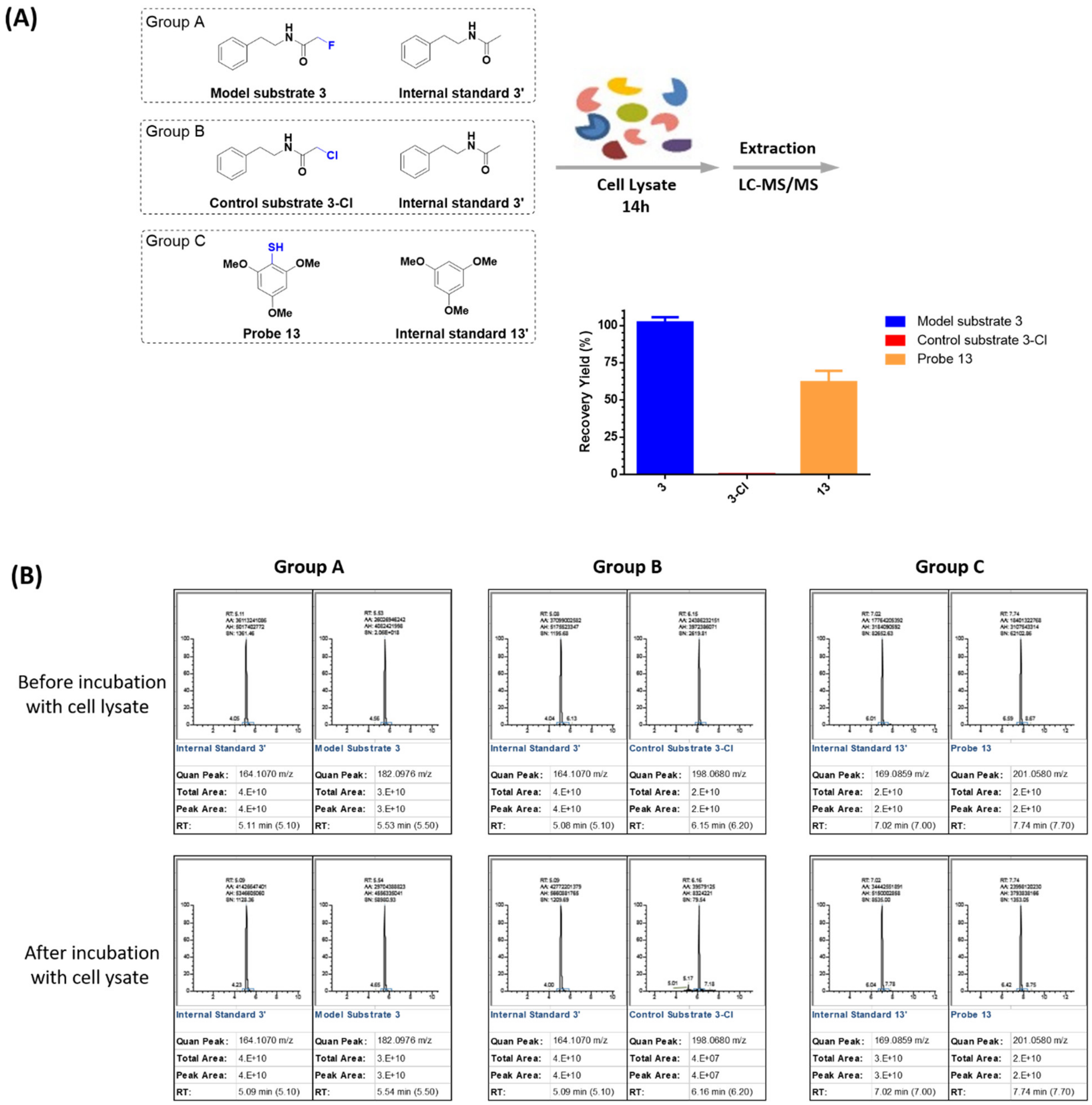


**Figure S10.** The stability and reactivity evaluations of the halo-acetamide tags and the probe **13** in mammalian cell lysates. (A) The model substrate **3**, control **3-Cl**, or the probe **13** was each mixed with the corresponding internal standard, and incubated within the HEK293 cell lysates for 14 h under the same FTDR reaction conditions, respectively. The compounds were then recovered by extraction and analyzed by LC-MS, generating a summary plot of recovery yields on the basis of triplicate repeats. (B) Representative results of the peak area and m/z information on LC-MS for each group analyzed before and after incubation with the cell lysates. The assays outlined in (A) and (B) were repeated in triplicate.

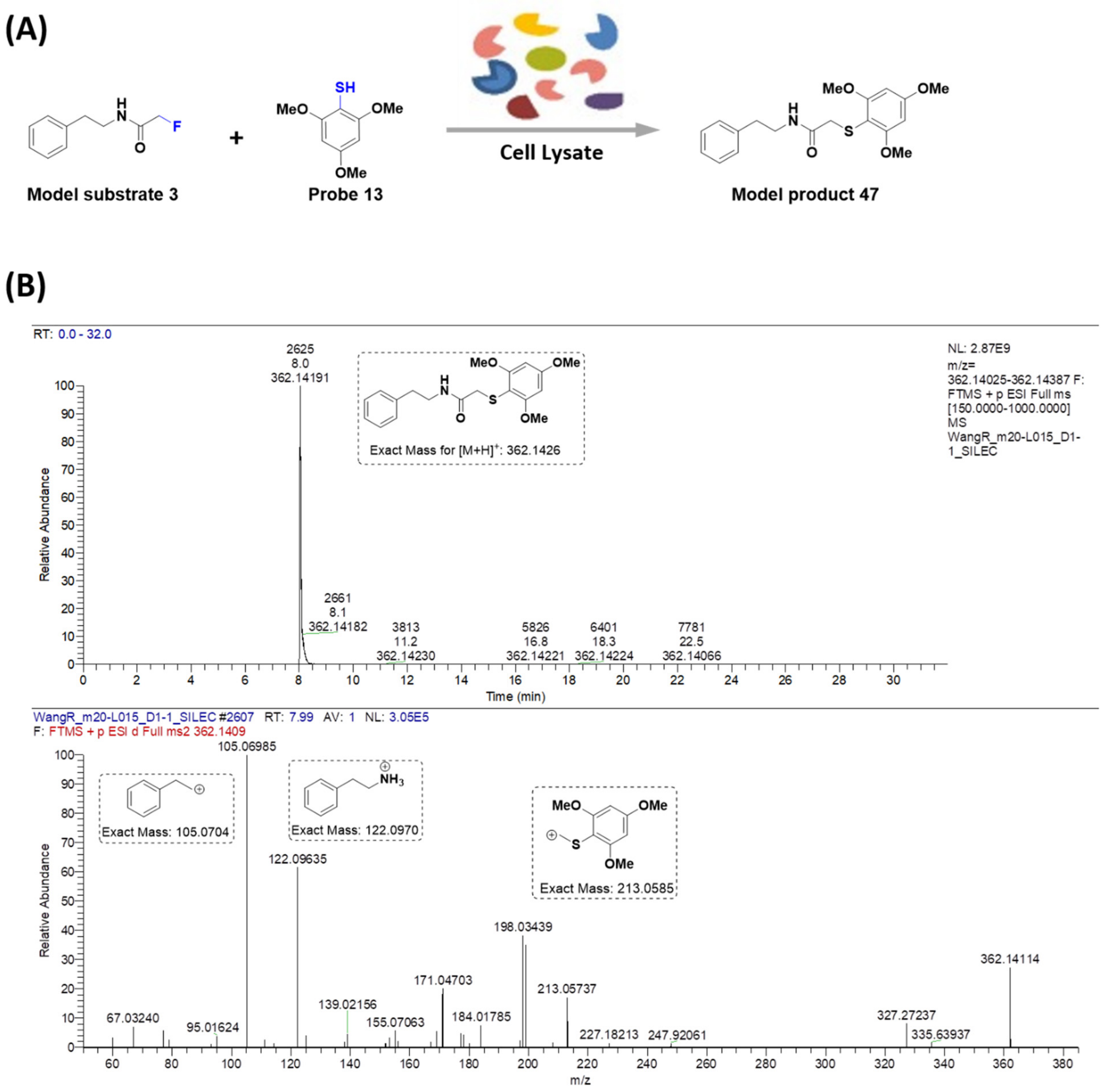


**Figure S11.** The FTDR reaction in mammalian cell lysates. (A) The model substrate **3** and probe **13** were mixed in the presence of HEK293 cell lysates. After incubation at 37ºC for 5h, the mixture was extracted and analyzed by LC-MS/MS. (B) LC-MS (top panel) and MS/MS (bottom panel) analysis of the cell lysate mixture identified the FTDR reaction product **47**.

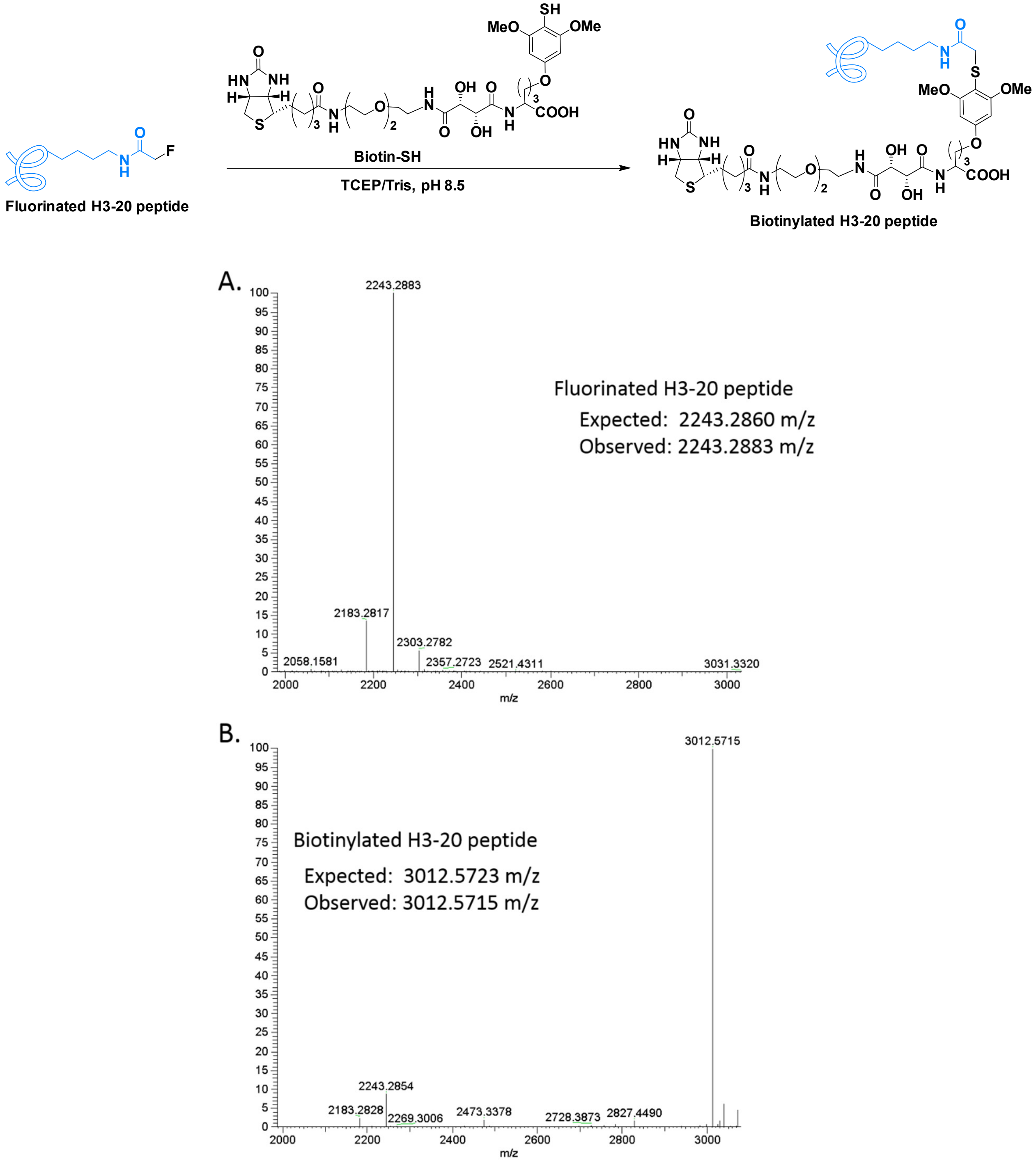


**Figure S12.** Biotinylation of fluorinated H3-20 peptide based on the Fluorine-Thiol Displacement Reaction. Shown below the reaction scheme is the ESI-MS spectra of the fluorinated H3-20 peptide (A) before and (B) after reacting with Biotin-SH during the process of biotinylation. H3-20 peptide sequence is "$NH_2$-Ala-Arg-Thr-Lys-Gln-Thr-Ala-Arg-Lys-Ser-Thr-Gly-Gly-Lys-Ala-Pro-Arg-Lys-Gln-Leu-COOH".

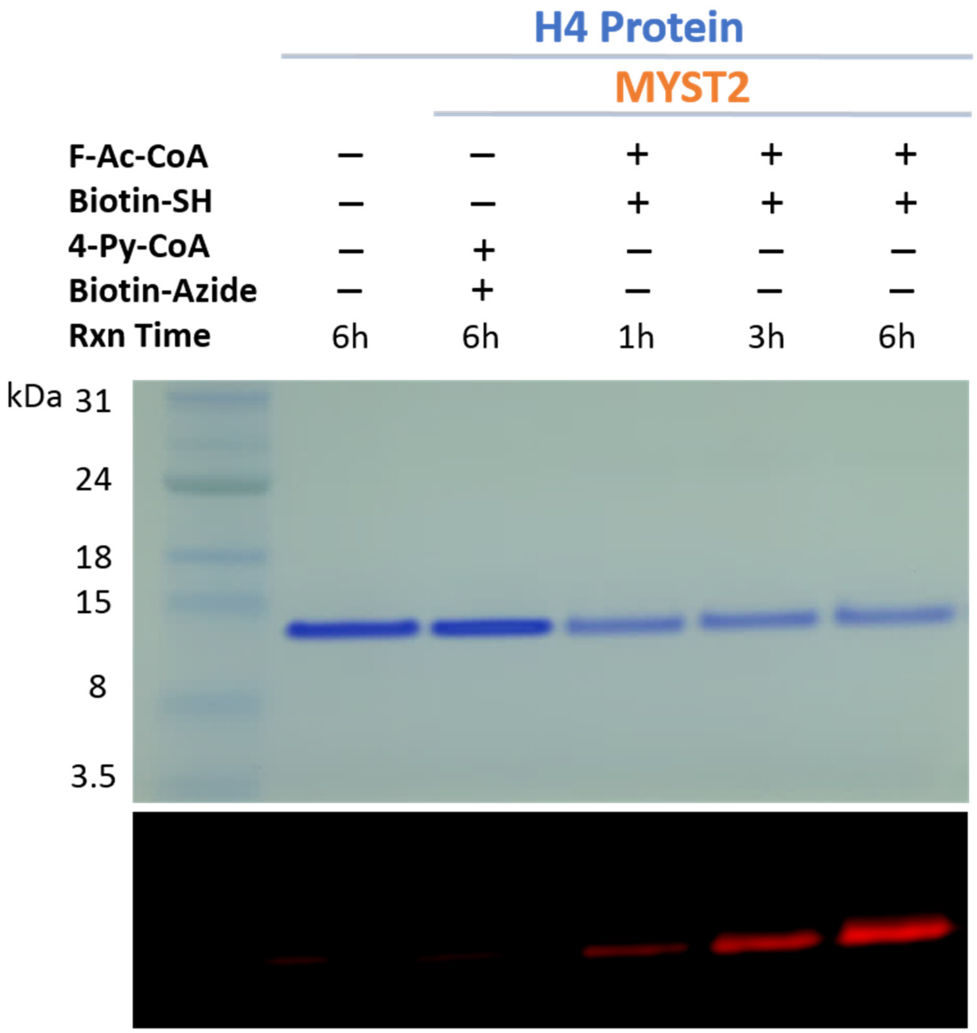


**Figure S13.** Biotinylation of H4 protein substrates by MYST2 enzyme. Biotin was detected by streptavidin-IRDye 680RD under near infrared fluorescence scanning (Ex 685 nm/Em 730 nm).

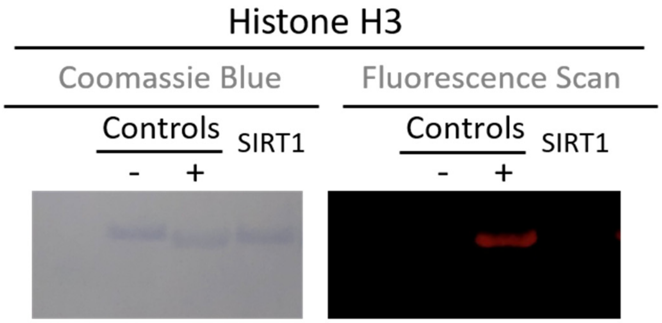


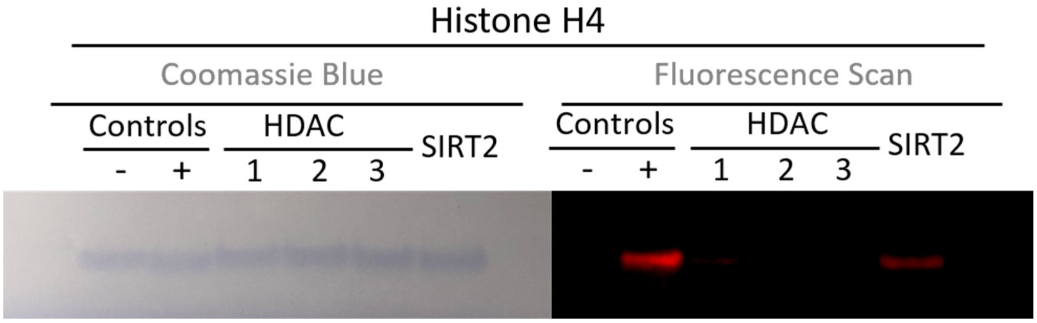


**Figure S14.** Western blot assays to evaluate the removal of F-acetyl groups on histone substrate by different histone deacetylases (SIRT1 for fluoroacetylated histone H3, HDACs 1-3 and SIRT2 for fluoroacetylated histone H4). Negative control "-": intact wild type H3 or H4; Positive control "+": fluoroacetylated H3 or H4 after treatment with F-Ac-CoA and the corresponding acetyltransferases. Other samples lanes are the mixture of fluoroacetylated H3 or H4 with different histone deacetylases.

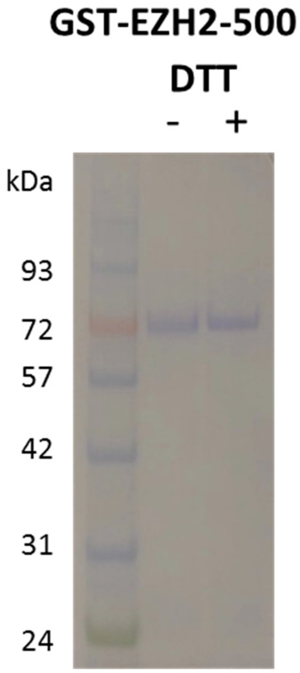


**Figure S15.** Characterization of EZH2 (1-500) protein by SDS-PAGE analysis.

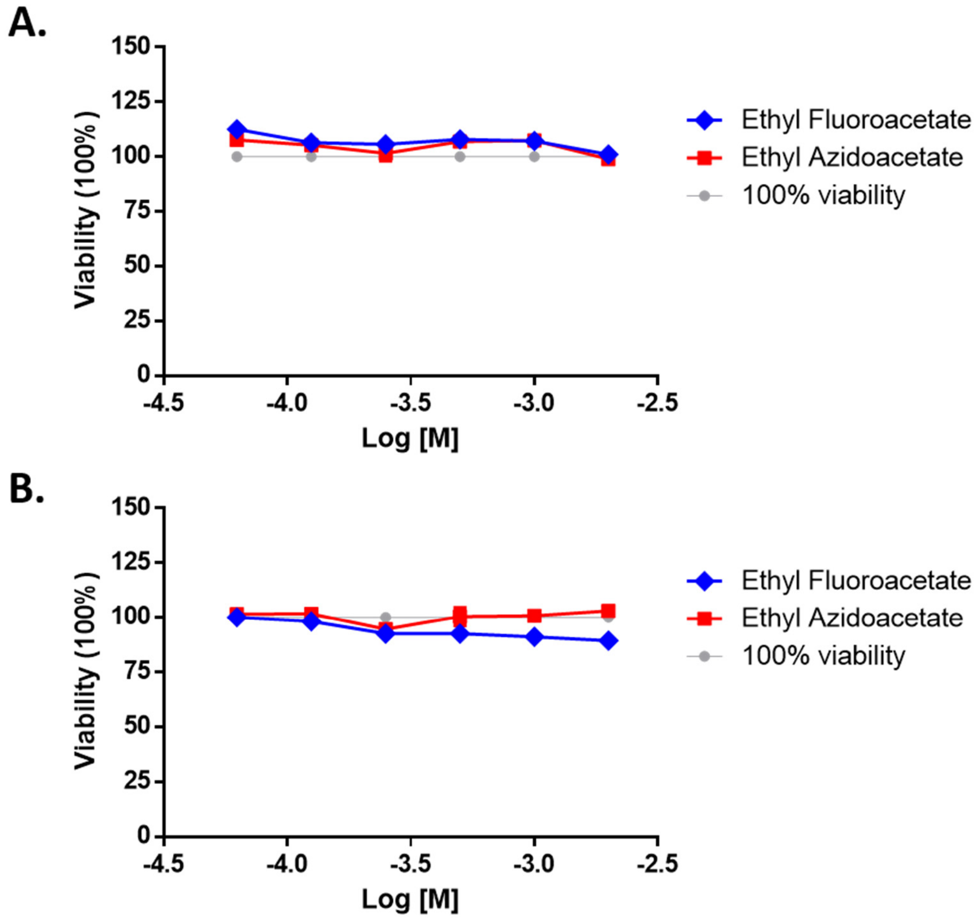


**Figure S16.** *In vitro* cell cytotoxicity assay to confirm the nontoxicity of ethyl ester pro-metabolites. Relative cytotoxicity on (A) HeLa cells and (B) HEK293T cells by ethyl fluoroacetate or the control ethyl azidoacetate after 12 h of incubation were plotted. Error bars represent SD of three replicates.

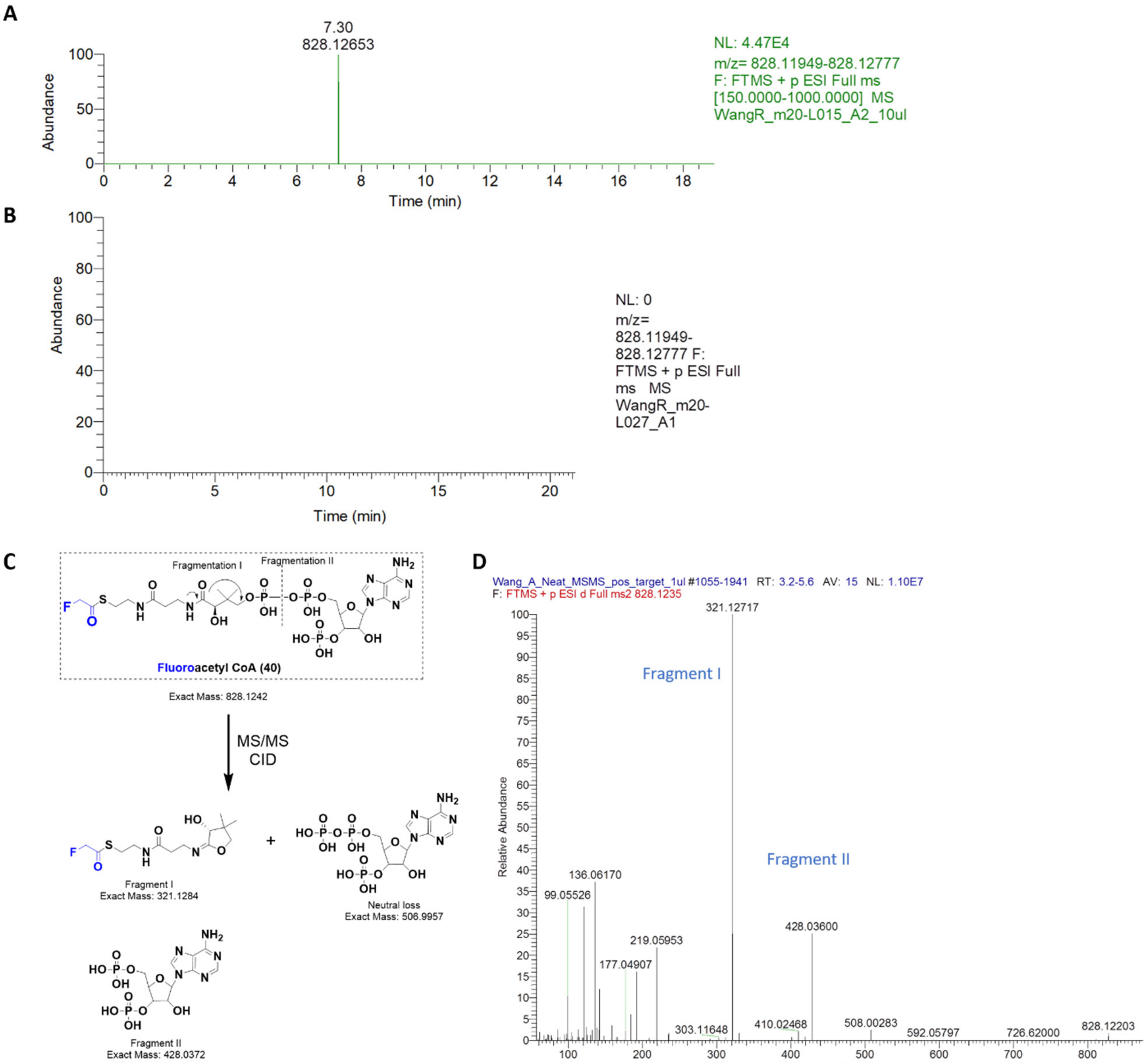


**Figure S17.** LC-MS/MS analysis of fluoroacetyl-CoA formation in HEK293 cells treated with the pro-metabolite fluoroacetate. (A) LC-MS of HEK293 cell extract demonstrating the eluted peak specific to fluoroacetyl-CoA (828.1265 m/z). The cells were treated with 1 mM ethyl fluoroacetate for 2 h. (B) LC-MS of extracts from preheated HEK293 lysates that was incubated with 1 mM ethyl fluoroacetate for 2h. (C) Product ions derived from fluoroacetyl-CoA for MS/MS analysis. (D) LC-MS/MS fragmentation analysis of the eluted fluoroacetyl-CoA peak.

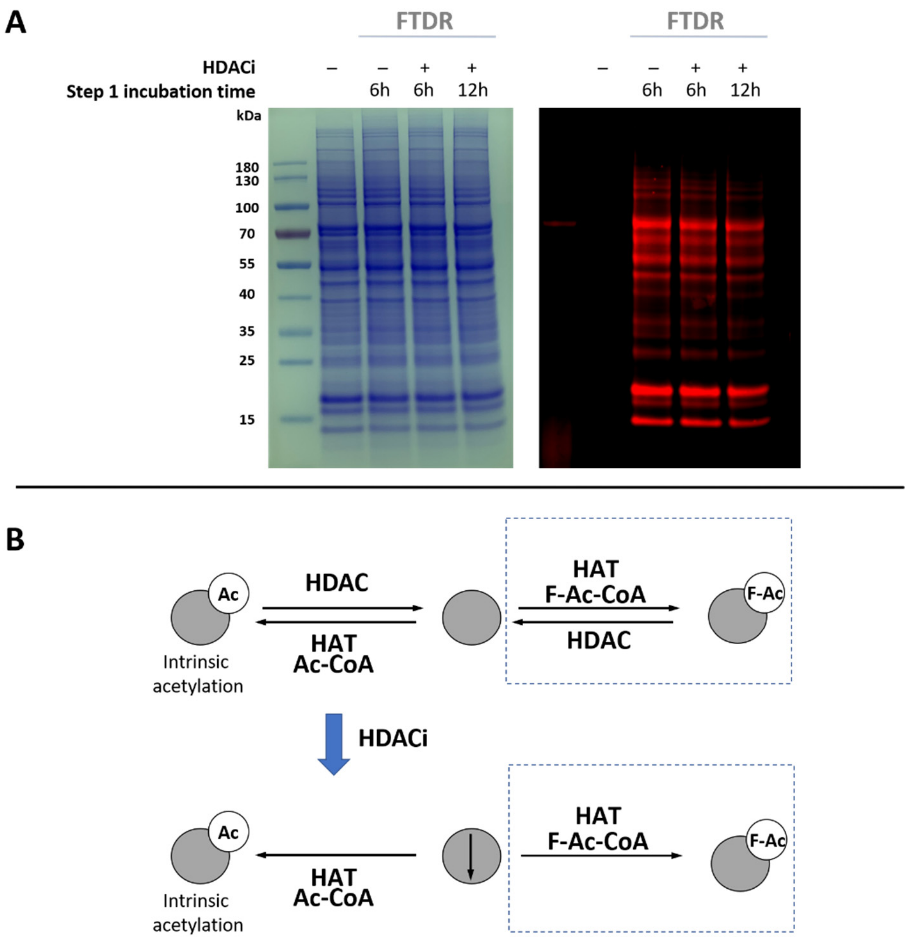


**Figure S18.** FTDR-based imaging of acetylation after concurrent HDAC inhibition. (A) HEK293 cells treated with the pro-metabolite ethyl fluoroacetate with or without the presence of the HDAC inhibitor cocktail (APExBIO) for 6h or 12h, before lysis and FTDR reaction. Left: CBB staining of all the cell lysate samples after FTDR reaction; Right: Fluorescent imaging. (B) The proposed hijacking of intrinsic acetylation. Under the dynamic acetylation-deacetylation equilibrium, the acetylated sites on protein substrates could be deacetylated to allow for modification by F-Ac (top row). With the equilibrium blocked by HDACi, the intrinsic acetylation may compete with F-acetylation.

**Database:** 20200916_WangR.fasta
**Protein ID:** C00004
**Monoisotopic Mass:** 18,029.73

**Source:** YZ-3-5 (non-Razor peptides included)
**Protein Name:** __CUST Histone H2B
**Amino Acid Coverage:** 99/166 = 59%

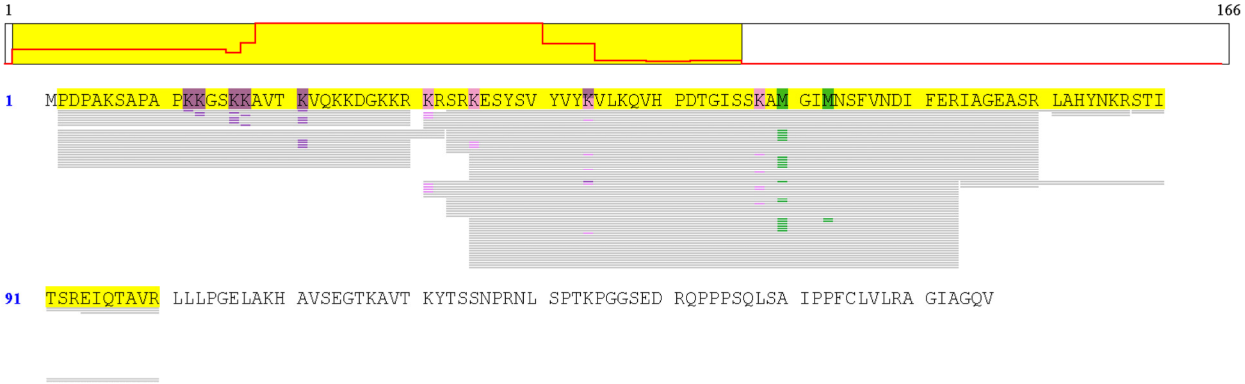


**Legends:**
+15.99 : Oxidation +42.01 : Acetyl +60.00 : F-Acetyl

| Peptide: | Count: | Position: |
|---|---|---|
| EIQTAVR | 1 | 94-100 |
| IAGEASR | 1 | 74-80 |
| IAGEASRLAHYNKRSTITSREIQTAVR | 2 | 74-100 |
| KESYSVYVYKVLKQVHPDTGISSKAMGIMNSFVNDIFER | 2 | 35-73 |
| KESYSVYVYKVLKQVHPDTGISSKAMGIMNSFVNDIFER | 4 | 35-73 |
| KESYSVYVYKVLKQVHPDTGISSKAMGIMNSFVNDIFER | 1 | 35-73 |
| KESYSVYVYKVLKQVHPDTGISSKAMGIMNSFVNDIFER | 14 | 35-73 |
| KESYSVYVYKVLKQVHPDTGISSKAMGIMNSFVNDIFERIAGEASR | 1 | 35-80 |
| KESYSVYVYKVLKQVHPDTGISSKAMGIMNSFVNDIFERIAGEASR | 3 | 35-80 |
| KESYSVYVYKVLKQVHPDTGISSKAMGIMNSFVNDIFERIAGEASR | 1 | 35-80 |
| KESYSVYVYKVLKQVHPDTGISSKAMGIMNSFVNDIFERIAGEASR | 1 | 35-80 |
| KESYSVYVYKVLKQVHPDTGISSKAMGIMNSFVNDIFERIAGEASR | 3 | 35-80 |
| KRSRKESYSVYVYKVLKQVHPDTGISSKAMGIMNSFVNDIFER | 1 | 31-73 |
| KRSRKESYSVYVYKVLKQVHPDTGISSKAMGIMNSFVNDIFER | 1 | 31-73 |
| KRSRKESYSVYVYKVLKQVHPDTGISSKAMGIMNSFVNDIFER | 2 | 31-73 |
| KRSRKESYSVYVYKVLKQVHPDTGISSKAMGIMNSFVNDIFER | 1 | 31-73 |
| KRSRKESYSVYVYKVLKQVHPDTGISSKAMGIMNSFVNDIFER | 2 | 31-73 |
| KRSRKESYSVYVYKVLKQVHPDTGISSKAMGIMNSFVNDIFERIAGEASR | 3 | 31-80 |
| KRSRKESYSVYVYKVLKQVHPDTGISSKAMGIMNSFVNDIFERIAGEASR | 1 | 31-80 |

| Peptide: | Count: | Position: |
|---|---|---|
| KRSRKESYSVYVYKVLKQVHPDTGISSKAMGIMNSFVNDIFERIAGEASR | 3 | 31-80 |
| LAHYNKR | 2 | 81-87 |
| PDPAKSAPAPKKGSKKAVTKVQKKDGKKR | 1 | 2-30 |
| PDPAKSAPAPKKGSKKAVTKVQKKDGKKR | 1 | 2-30 |
| PDPAKSAPAPKKGSKKAVTKVQKKDGKKR | 1 | 2-30 |
| PDPAKSAPAPKKGSKKAVTKVQKKDGKKR | 3 | 2-30 |
| PDPAKSAPAPKKGSKKAVTKVQKKDGKKR | 1 | 2-30 |
| PDPAKSAPAPKKGSKKAVTKVQKKDGKKR | 4 | 2-30 |
| PDPAKSAPAPKKGSKKAVTKVQKKDGKKR | 8 | 2-30 |
| PDPAKSAPAPKKGSKKAVTKVQKKDGKKRKR | 4 | 2-32 |
| SRKESYSVYVYKVLKQVHPDTGISSKAMGIMNSFVNDIFER | 2 | 33-73 |
| SRKESYSVYVYKVLKQVHPDTGISSKAMGIMNSFVNDIFER | 1 | 33-73 |
| SRKESYSVYVYKVLKQVHPDTGISSKAMGIMNSFVNDIFER | 5 | 33-73 |
| SRKESYSVYVYKVLKQVHPDTGISSKAMGIMNSFVNDIFERIAGEASR | 5 | 33-80 |
| SRKESYSVYVYKVLKQVHPDTGISSKAMGIMNSFVNDIFERIAGEASR | 3 | 33-80 |
| SRKESYSVYVYKVLKQVHPDTGISSKAMGIMNSFVNDIFERIAGEASR | 2 | 33-80 |
| SRKESYSVYVYKVLKQVHPDTGISSKAMGIMNSFVNDIFERIAGEASRLAHYNKR | 1 | 33-87 |
| STITSREIQTAVR | 2 | 88-100 |

**Figure S19.** The proteomics analysis of histone H2B which was extracted from HEK293 cells after treatment with pro-metabolite ethyl fluoroacetate. Pink legend: F-acetylation; Purple legend: wild type acetylation. The N-terminal amino acid is proline.

**Database:** 20200916_WangR.fasta
**Protein ID:** C00002
**Monoisotopic Mass:** 11,360.39

**Source:** ZL-2-5 (non-Razor peptides included)
**Protein Name:** __CUST Histone H4
**Amino Acid Coverage:** 96/103 = 93%

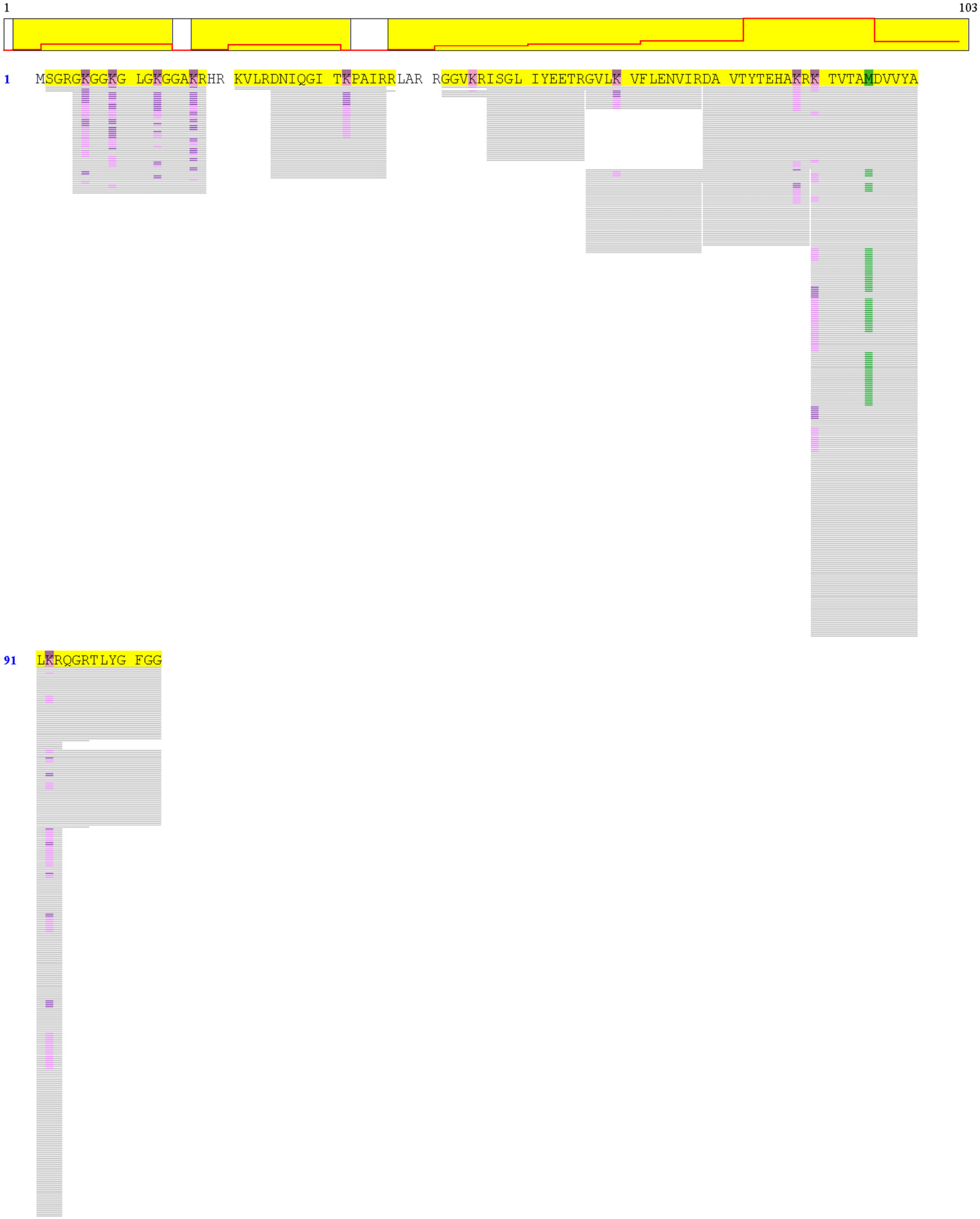


**Legends:**
+15.99 : Oxidation    +42.01 : Acetyl    +60.00 : F-Acetyl

**Legends:**
■ +15.99 : Oxidation ■ +42.01 : Acetyl ■ +60.00 : F-Acetyl

| Peptide: | Count: | Position: |
|---|---|---|
| DAVTYTEHAKR | 3 | 69-79 |
| DAVTYTEHAKR | 8 | 69-79 |
| DAVTYTEHAKR | 22 | 69-79 |
| DAVTYTEHAKRKTVTAMDVVYALKR | 1 | 69-93 |
| DAVTYTEHAKRKTVTAMDVVYALKR | 2 | 69-93 |
| DAVTYTEHAKRKTVTAMDVVYALKR | 1 | 69-93 |
| DAVTYTEHAKRKTVTAMDVVYALKRQGR | 1 | 69-96 |
| DAVTYTEHAKRKTVTAMDVVYALKRQGRTLYGFGG | 3 | 69-103 |
| DAVTYTEHAKRKTVTAMDVVYALKRQGRTLYGFGG | 1 | 69-103 |
| DAVTYTEHAKRKTVTAMDVVYALKRQGRTLYGFGG | 9 | 69-103 |
| DAVTYTEHAKRKTVTAMDVVYALKRQGRTLYGFGG | 2 | 69-103 |
| DAVTYTEHAKRKTVTAMDVVYALKRQGRTLYGFGG | 4 | 69-103 |
| DAVTYTEHAKRKTVTAMDVVYALKRQGRTLYGFGG | 19 | 69-103 |
| DNIQGITKPAIR | 7 | 25-36 |
| DNIQGITKPAIR | 17 | 25-36 |
| DNIQGITKPAIR | 21 | 25-36 |
| DNIQGITKPAIRR | 1 | 25-37 |
| GGVKRISGLIYEETR | 1 | 42-56 |
| GGVKRISGLIYEETR | 3 | 42-56 |
| GGVKRISGLIYEETRGVLKVFLENVIR | 1 | 42-68 |
| GKGGKGLGKGGAKR | 4 | 5-18 |
| GKGGKGLGKGGAKR | 2 | 5-18 |
| GKGGKGLGKGGAKR | 1 | 5-18 |
| GKGGKGLGKGGAKR | 2 | 5-18 |
| GKGGKGLGKGGAKR | 1 | 5-18 |
| GKGGKGLGKGGAKR | 2 | 5-18 |
| GKGGKGLGKGGAKR | 2 | 5-18 |
| GKGGKGLGKGGAKR | 2 | 5-18 |
| GKGGKGLGKGGAKR | 1 | 5-18 |
| GKGGKGLGKGGAKR | 1 | 5-18 |
| GKGGKGLGKGGAKR | 2 | 5-18 |
| GKGGKGLGKGGAKR | 1 | 5-18 |
| GKGGKGLGKGGAKR | 3 | 5-18 |
| GKGGKGLGKGGAKR | 2 | 5-18 |
| GKGGKGLGKGGAKR | 2 | 5-18 |
| GKGGKGLGKGGAKR | 1 | 5-18 |
| GKGGKGLGKGGAKR | 1 | 5-18 |
| GKGGKGLGKGGAKR | 2 | 5-18 |
| GKGGKGLGKGGAKR | 1 | 5-18 |
| GKGGKGLGKGGAKR | 1 | 5-18 |
| GKGGKGLGKGGAKR | 2 | 5-18 |
| GKGGKGLGKGGAKR | 2 | 5-18 |
| GKGGKGLGKGGAKR | 1 | 5-18 |
| GKGGKGLGKGGAKR | 2 | 5-18 |

| Peptide: | Count: | Position: |
|---|---|---|
| GKGGKGLGKGGAKR | 2 | 5-18 |
| GKGGKGLGKGGAKR | 2 | 5-18 |
| GKGGKGLGKGGAKR | 1 | 5-18 |
| GKGGKGLGKGGAKR | 2 | 5-18 |
| GKGGKGLGKGGAKR | 2 | 5-18 |
| GKGGKGLGKGGAKR | 3 | 5-18 |
| GVLKVFLENVIR | 4 | 57-68 |
| GVLKVFLENVIR | 6 | 57-68 |
| GVLKVFLENVIR | 37 | 57-68 |
| GVLKVFLENVIRDAVTYTEHAKR | 1 | 57-79 |
| GVLKVFLENVIRDAVTYTEHAKR | 3 | 57-79 |
| GVLKVFLENVIRDAVTYTEHAKR | 3 | 57-79 |
| ISGLIYEETR | 33 | 47-56 |
| ISGLIYEETRGVLKVFLENVIR | 1 | 47-68 |
| KTVTAMDVVYALKR | 1 | 80-93 |
| KTVTAMDVVYALKR | 6 | 80-93 |
| KTVTAMDVVYALKR | 2 | 80-93 |
| KTVTAMDVVYALKR | 11 | 80-93 |
| KTVTAMDVVYALKR | 3 | 80-93 |
| KTVTAMDVVYALKR | 1 | 80-93 |
| KTVTAMDVVYALKR | 2 | 80-93 |
| KTVTAMDVVYALKR | 18 | 80-93 |
| KTVTAMDVVYALKR | 2 | 80-93 |
| KTVTAMDVVYALKR | 8 | 80-93 |
| KTVTAMDVVYALKR | 28 | 80-93 |
| KTVTAMDVVYALKR | 7 | 80-93 |
| KTVTAMDVVYALKR | 4 | 80-93 |
| KTVTAMDVVYALKR | 13 | 80-93 |
| KTVTAMDVVYALKR | 19 | 80-93 |
| KTVTAMDVVYALKR | 77 | 80-93 |
| KTVTAMDVVYALKRQGR | 1 | 80-96 |
| KTVTAMDVVYALKRQGRTLYGFGG | 2 | 80-103 |
| KTVTAMDVVYALKRQGRTLYGFGG | 2 | 80-103 |
| KTVTAMDVVYALKRQGRTLYGFGG | 1 | 80-103 |
| KTVTAMDVVYALKRQGRTLYGFGG | 2 | 80-103 |
| KTVTAMDVVYALKRQGRTLYGFGG | 5 | 80-103 |
| KTVTAMDVVYALKRQGRTLYGFGG | 2 | 80-103 |
| KTVTAMDVVYALKRQGRTLYGFGG | 3 | 80-103 |
| KTVTAMDVVYALKRQGRTLYGFGG | 4 | 80-103 |
| KTVTAMDVVYALKRQGRTLYGFGG | 19 | 80-103 |
| KVLRDNIQGITKPAIR | 2 | 21-36 |
| SGRGKGGKGLGKGGAKR | 1 | 2-18 |
| SGRGKGGKGLGKGGAKR | 2 | 2-18 |

**Figure S20.** The proteomics analysis of histone H4 which was extracted from HEK293 cells after treatment with pro-metabolite ethyl fluoroacetate. Pink legend: F-acetylation; Purple legend: wild type acetylation. The N-terminal amino acid is serine.

**Database:** UP20191010_Human_9606.fasta
**Protein ID:** Q71U36
**Monoisotopic Mass:** 50,103.61

**Source:** C (non-Razor peptides included)
**Protein Name:** Tubulin alpha-1A chain OS=Homo sapiens OX=9606 GN=TUBA1A PE=1 SV=1
**Amino Acid Coverage:** 209/451 = 46%

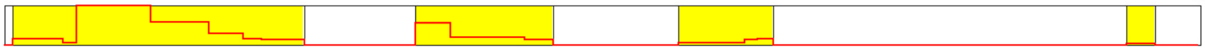


1 MRECISIHVG QAGVQIGNAC WELYCLEHGI QPDGQMPSDK TIGGGDDSFN TFFSETGAGK HVPRAVFVDL EPTVIDEVRT GTYRQLFHPE

91 QLITGKEDAA NNYARGHYTI GKEIIDLVLD RIRKLADQCT GLQGFLVFHS FGGGTGSGFT SLLMERLSVD YGKKSKLEFS IYPAPQVSTA

181 VVEPYNSILT THTTLEHSDC AFMVDNEAIY DICRRNLDIE RPTYTNLNRL IGQIVSSITA SLRFDGALNV DLTEFQTNLV PYPRIHFPLA

271 TYAPVISAEK AYHEQLSVAE ITNACFEPAN QMVKCDPRHG KYMACCLLYR GDVVPKDVNA AIATIKTKRT IQFVDWCPTG FKVGINYQPP

361 TVVPGGDLAK VQRAVCMLSN TTAIAEAWAR LDHKFDLMYA KRAFVHWYVG EGMEEGEFSE AREDMAALEK DYEEVGVDSV EGEGEEEGEE

451 Y

**Legends:**
+0.98 : Deamidation +15.99 : Oxidation +42.01 : Acetyl +60.00 : F-Acetyl

| Peptide: | Count: | Position: | Peptide: | Count: | Position: |
|---|---|---|---|---|---|
| CISIHVGQAGVQIGNACWE | 5 | 4-22 | KAYHEQLSVAE | 1 | 280-290 |
| DAANNYARGHYTIGKE | 1 | 98-113 | KAYHEQLSVAE | 3 | 280-290 |
| DMAALEKDYEE | 1 | 424-434 | LYCLEHGIQPDGQMPSDKTIGGGDDSFNTFFSE | 2 | 23-55 |
| FQTNLVPYPRIHFPLATYAPVISAE | 2 | 255-279 | QLITGKEDAANNYARGHYTIGKE | 3 | 91-113 |
| FSIYPAPQVSTAVVEPYNSILTTHTTLE | 1 | 169-196 | QLSVAE | 1 | 285-290 |
| FSIYPAPQVSTAVVEPYNSILTTHTTLE | 5 | 169-196 | RLSVDYGKKSKLE | 1 | 156-168 |
| HGIQPDGQMPSDKTIGGGDDSFNTFFSE | 6 | 28-55 | RLSVDYGKKSKLE | 1 | 156-168 |
| HGIQPDGQMPSDKTIGGGDDSFNTFFSE | 9 | 28-55 | RLSVDYGKKSKLE | 2 | 156-168 |
| HGIQPDGQMPSDKTIGGGDDSFNTFFSE | 1 | 28-55 | RLSVDYGKKSKLE | 13 | 156-168 |
| HGIQPDGQMPSDKTIGGGDDSFNTFFSE | 1 | 28-55 | TGAGKHVPRAVFVDLEPTVIDE | 1 | 56-77 |
| HGIQPDGQMPSDKTIGGGDDSFNTFFSE | 10 | 28-55 | TGAGKHVPRAVFVDLEPTVIDE | 12 | 56-77 |
| HGIQPDGQMPSDKTIGGGDDSFNTFFSETGAGKHVPRAVFVDLEPTVIDE | 2 | 28-77 | TGAGKHVPRAVFVDLEPTVIDEVRTGTYRQLFHPE | 3 | 56-90 |
| HSDCAFMVDNE | 2 | 197-207 | VRTGTYRQLFHPE | 4 | 78-90 |
| HSDCAFMVDNE | 2 | 197-207 | VRTGTYRQLFHPEQLITGKE | 2 | 78-97 |

**Figure S21.** The proteomics analysis of alpha-tubulin which was immunoenriched from HEK293 cells after treatment with pro-metabolite ethyl fluoroacetate. Pink legend: F-acetylation; Purple legend: wild type acetylation.

#### $^{1}$H-, $^{13}$C-, and $^{19}$F NMR Spectra for New Compounds

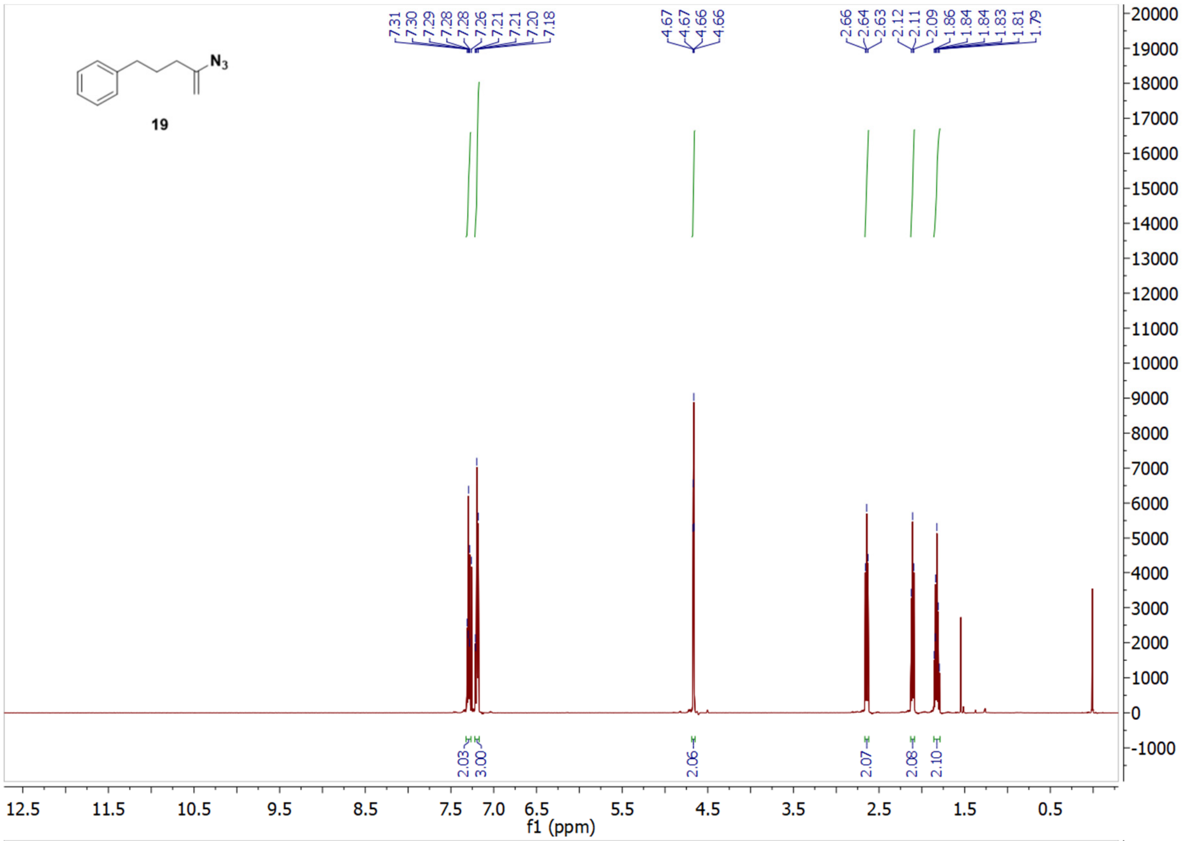


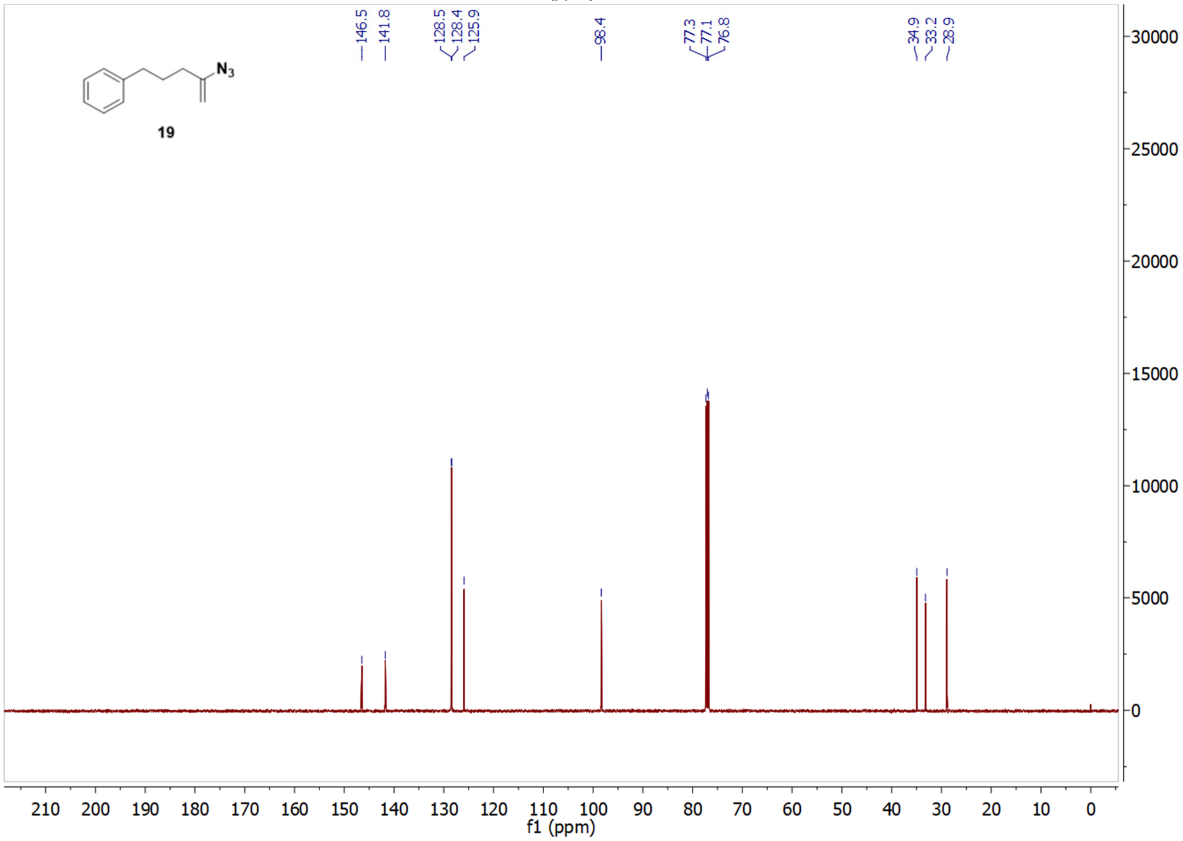

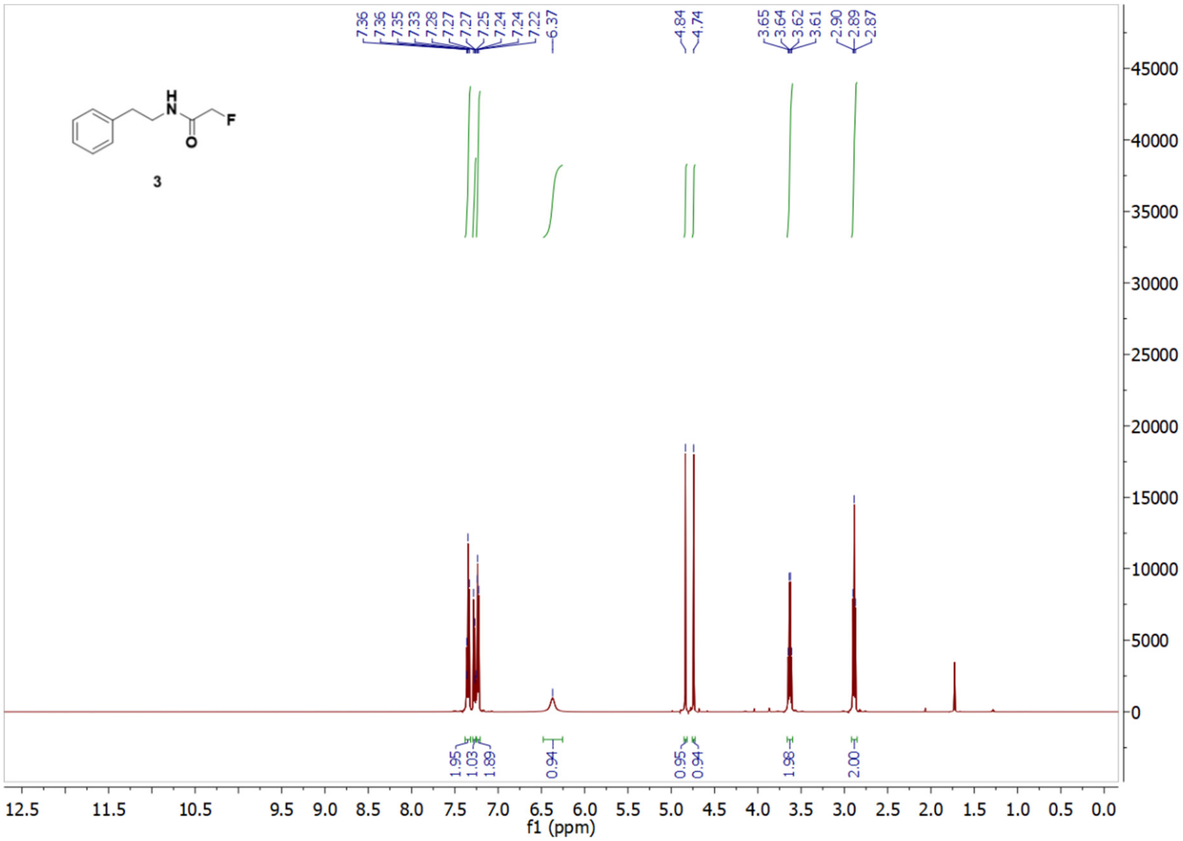

7.36
7.36
7.35
7.33
7.28
7.27
7.27
7.25
7.24
7.24
7.22
6.37
4.84
4.74
3.65
3.64
3.62
3.61
2.90
2.89
2.87
H
N
F
O
3
1.95
1.03
1.89
0.94
0.95
0.94
1.98
2.00
f1 (ppm)


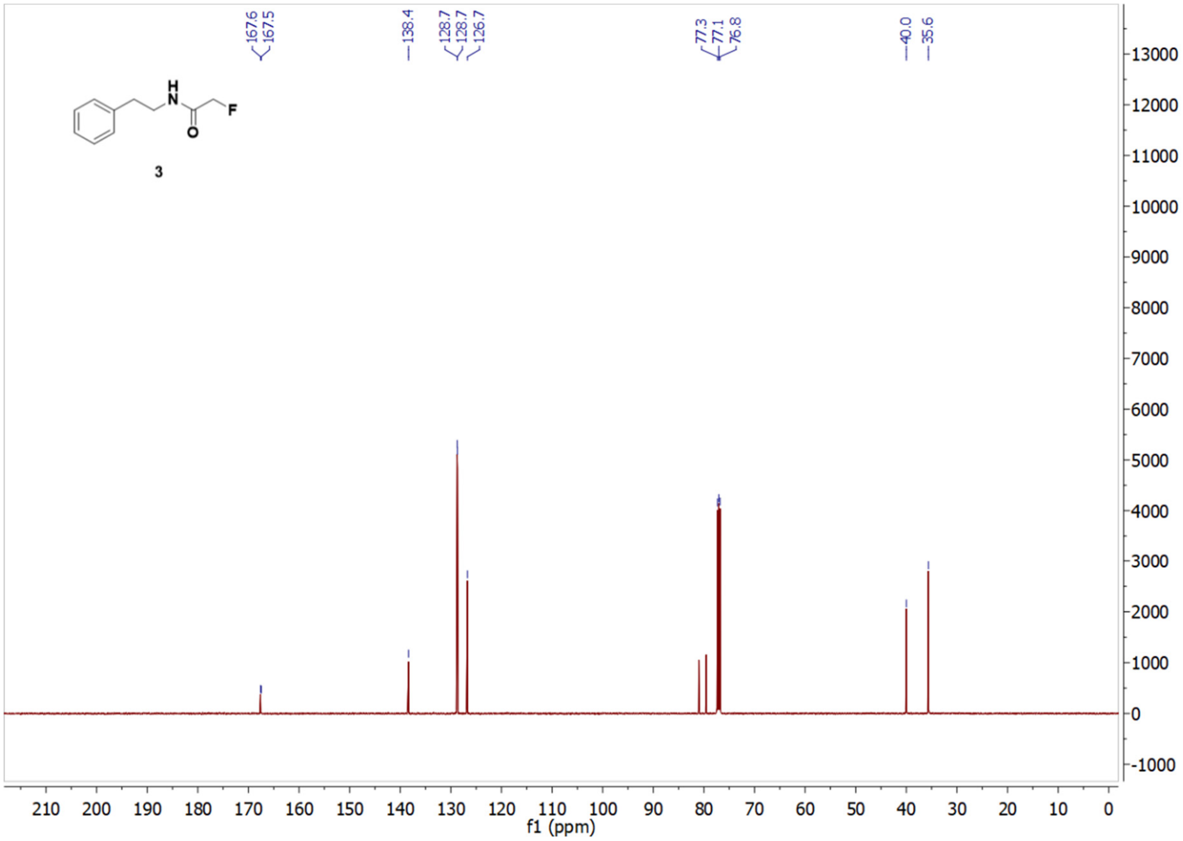

167.6
167.5
138.4
128.7
128.7
126.7
77.3
77.1
76.8
40.0
35.6
H
N
F
O
3
f1 (ppm)

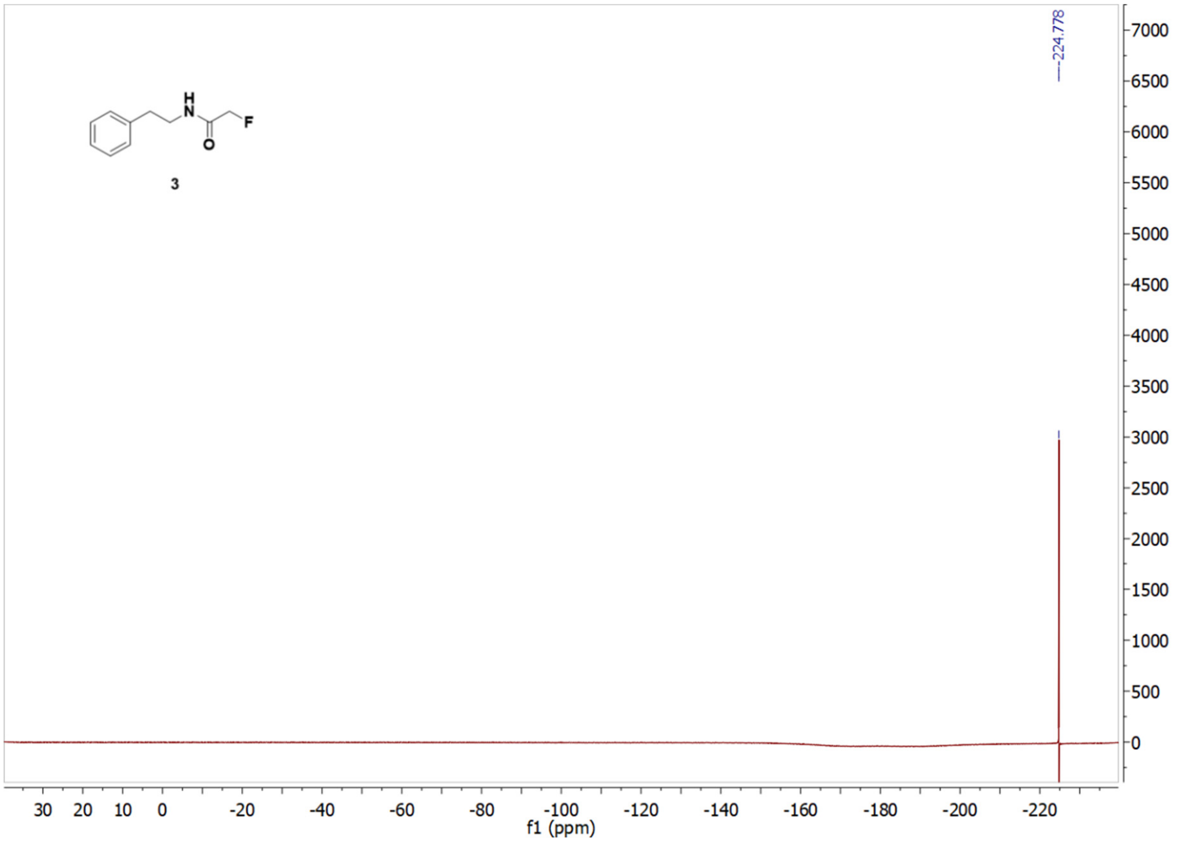

3
-224.778
f1 (ppm)


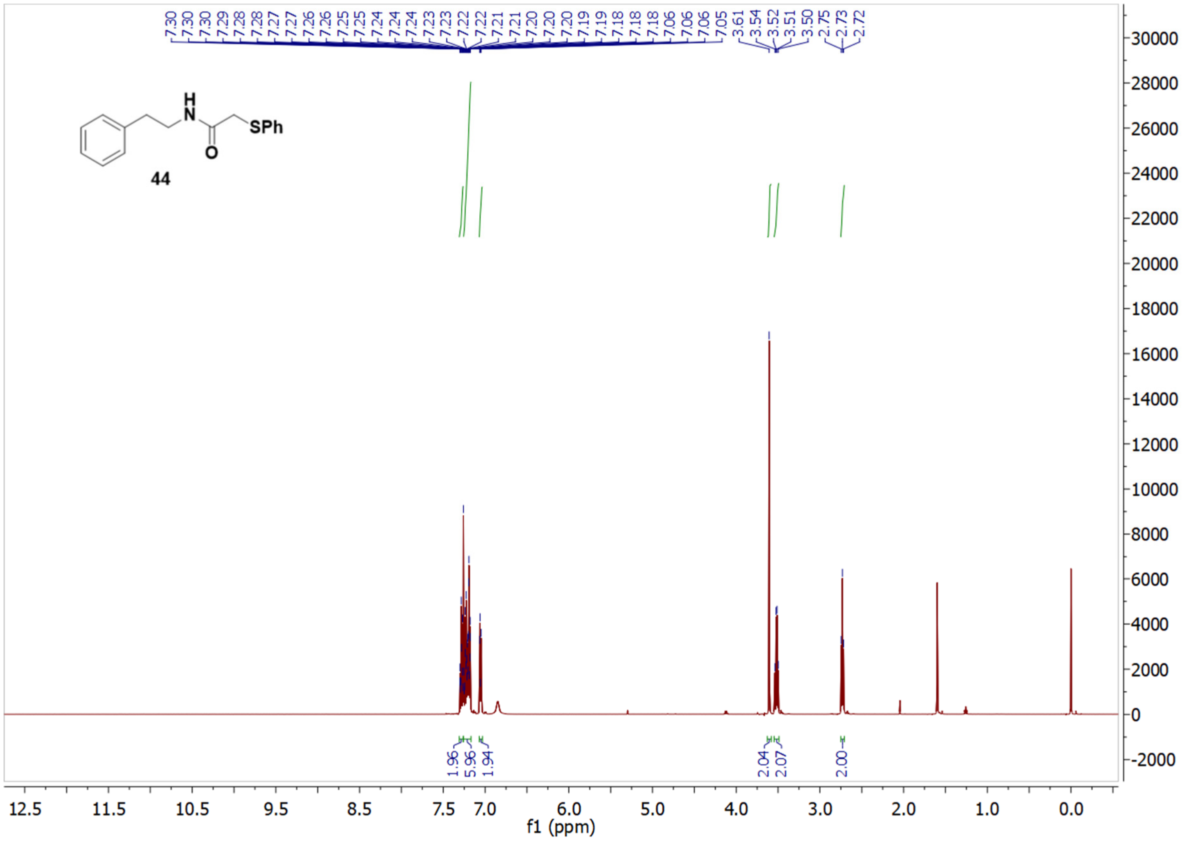

44
SPh
f1 (ppm)

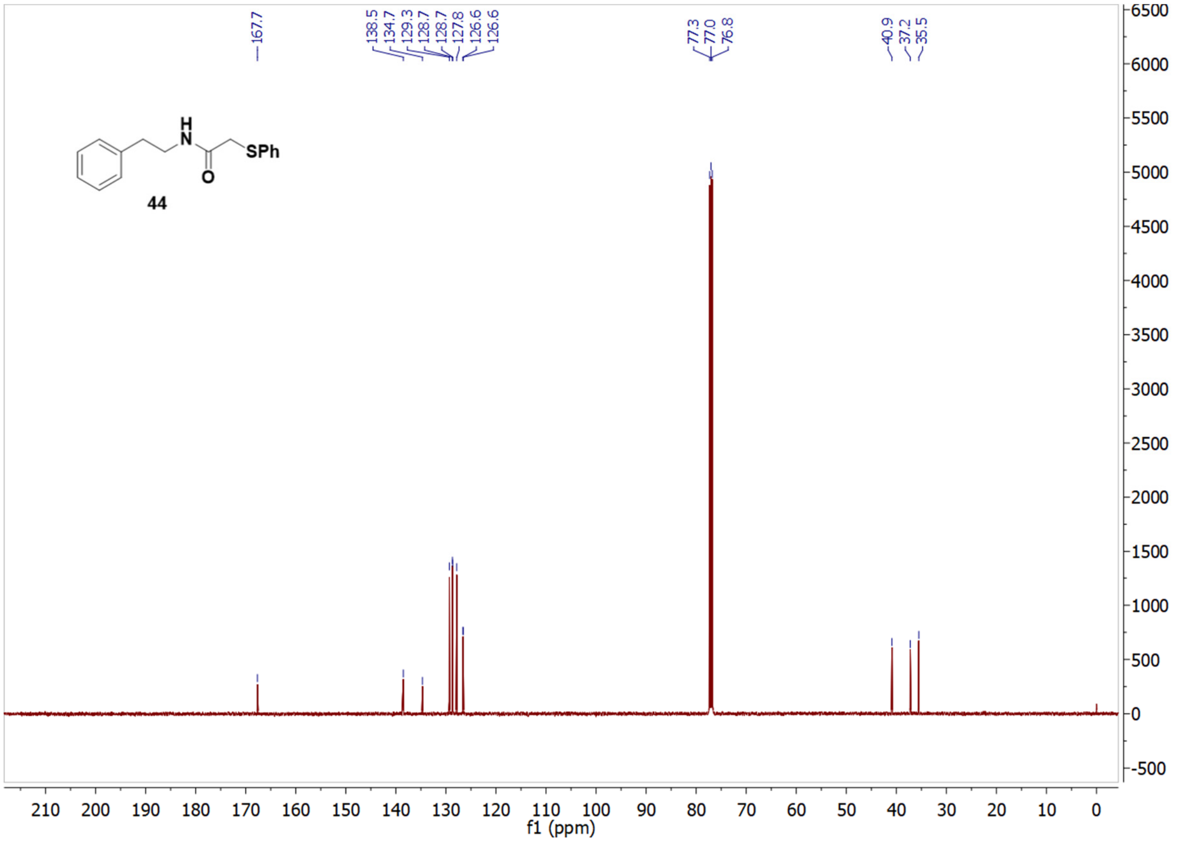
SPh
O
44
167.7
138.5
134.7
129.3
128.7
128.7
127.8
126.6
126.6
77.3
77.0
76.8
40.9
37.2
35.5
f1 (ppm)

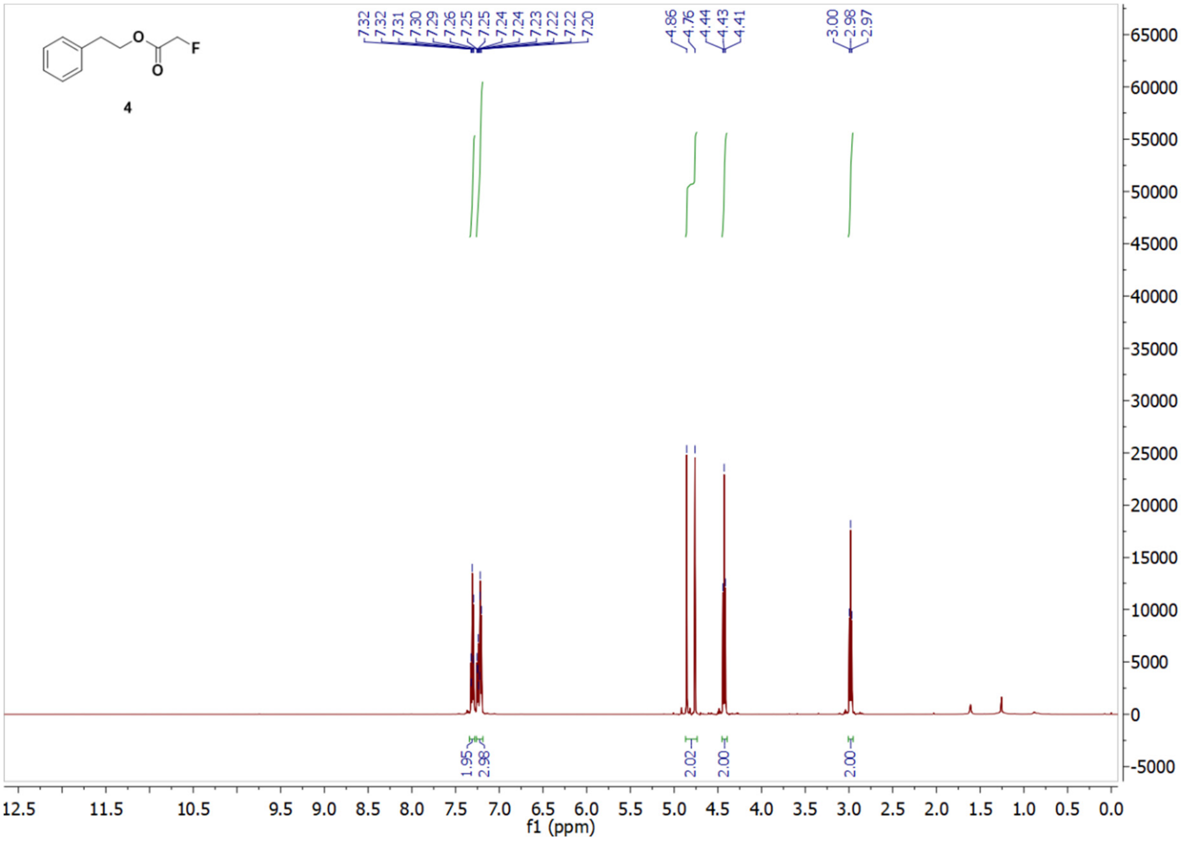
O
F
4
7.32
7.32
7.31
7.30
7.29
7.26
7.25
7.25
7.24
7.24
7.23
7.22
7.22
7.20
4.86
4.76
4.44
4.43
4.41
3.00
2.98
2.97
1.95
2.98
2.02
2.00
2.00
f1 (ppm)

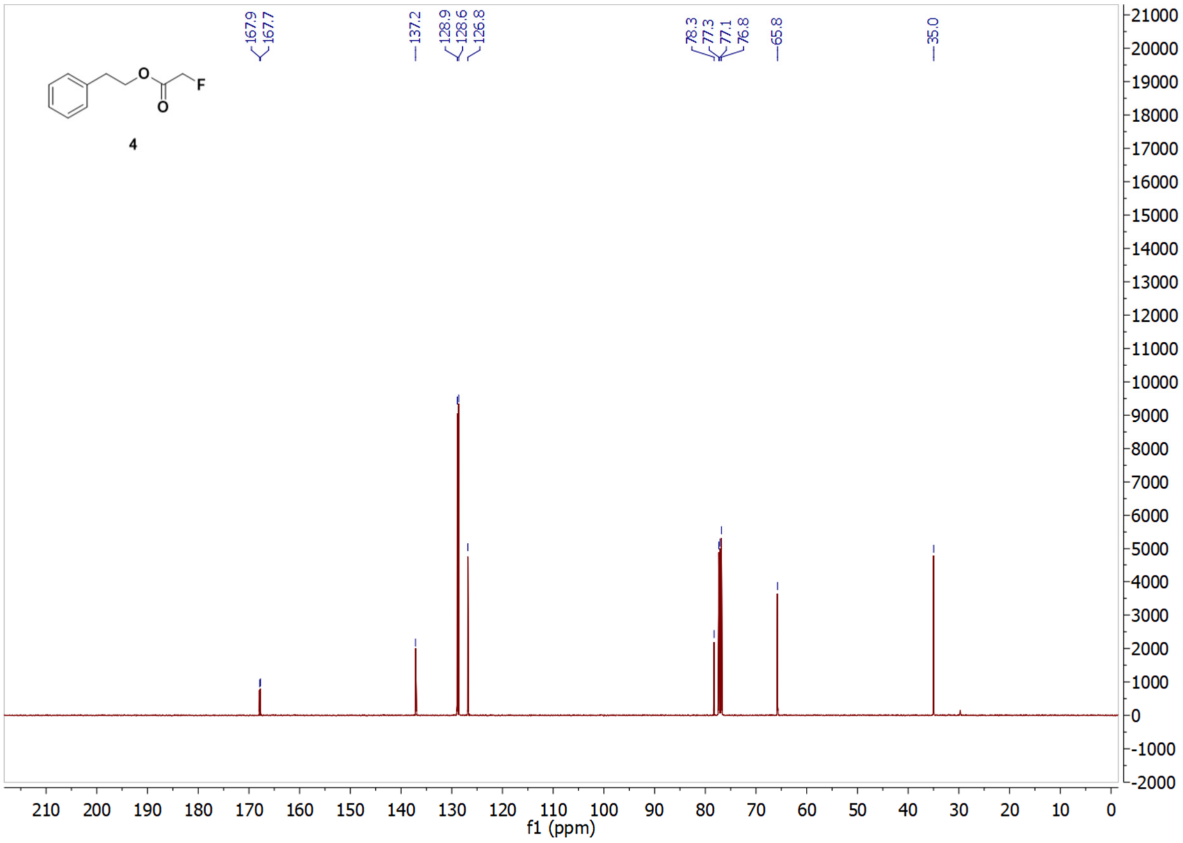

167.9
167.7
137.2
128.9
128.6
126.8
78.3
77.3
77.1
76.8
65.8
35.0
O
F
O
4
f1 (ppm)


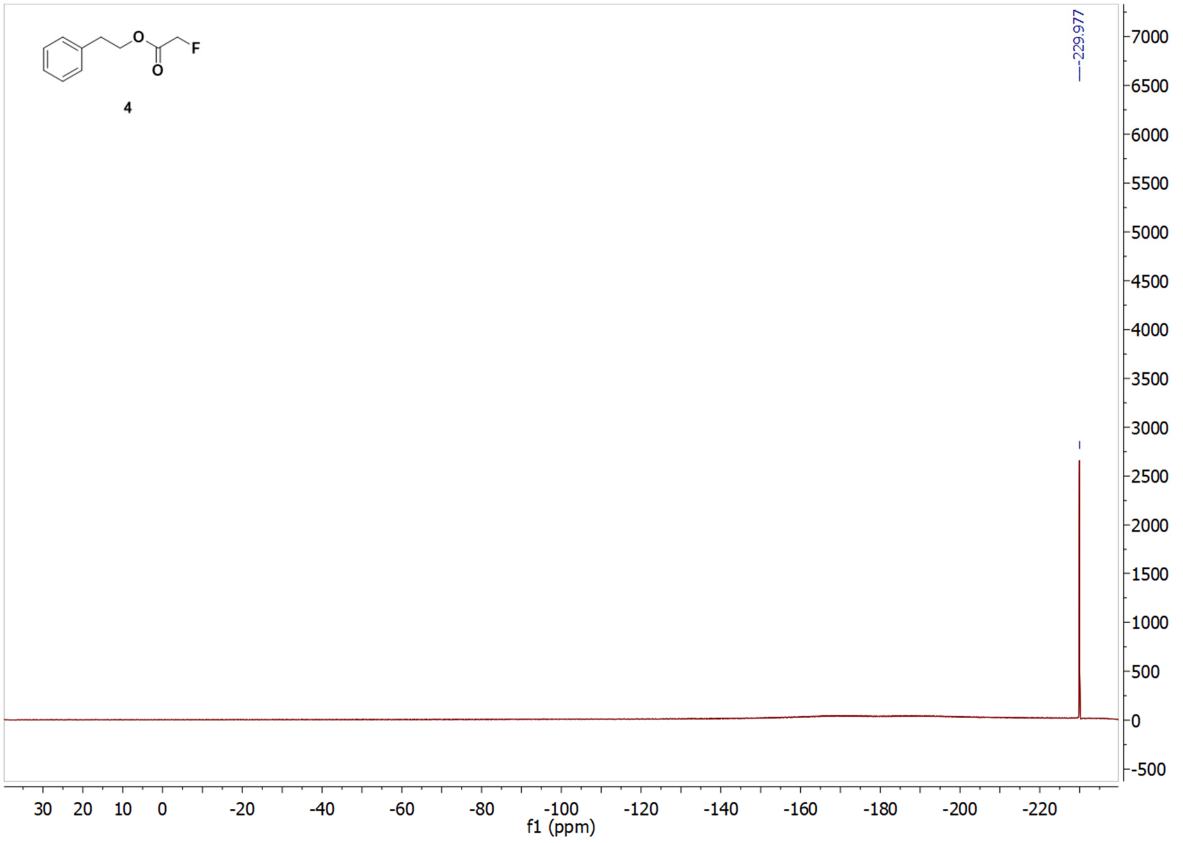

-229.977
O
F
O
4
f1 (ppm)

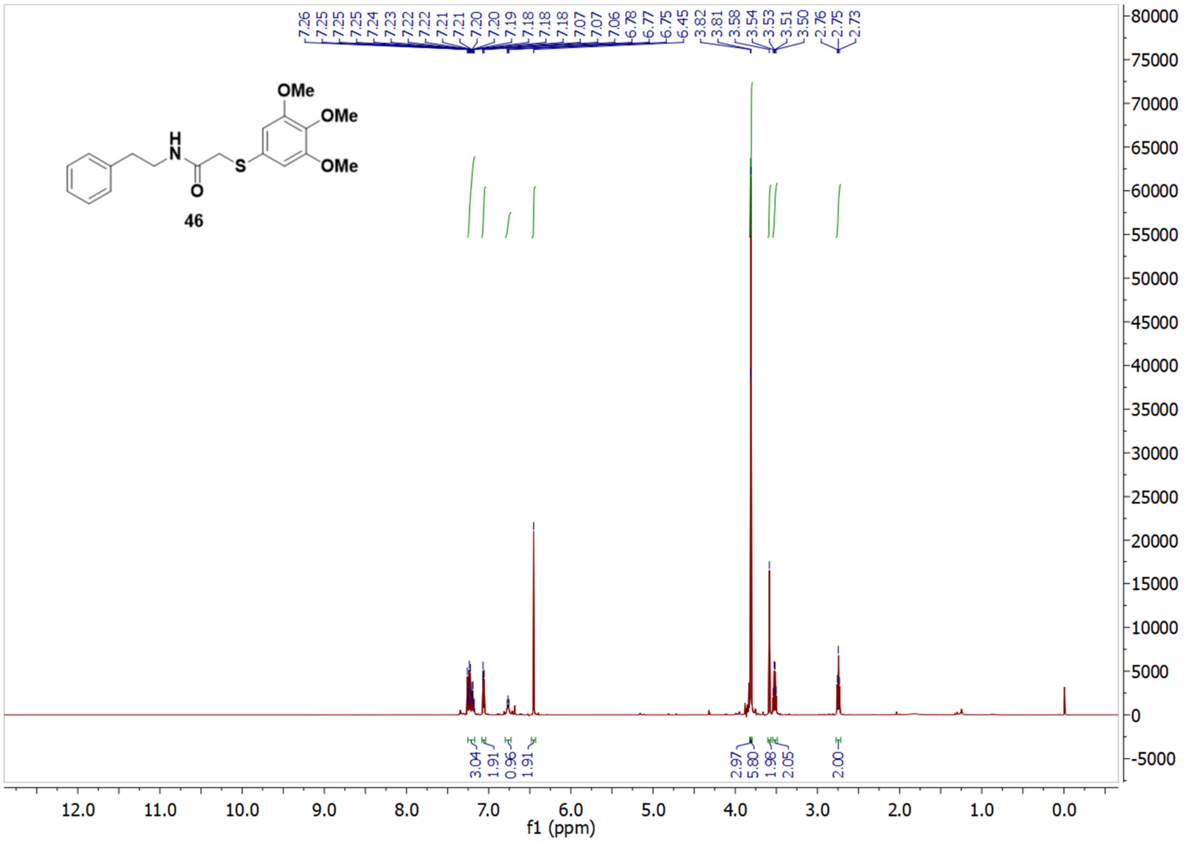

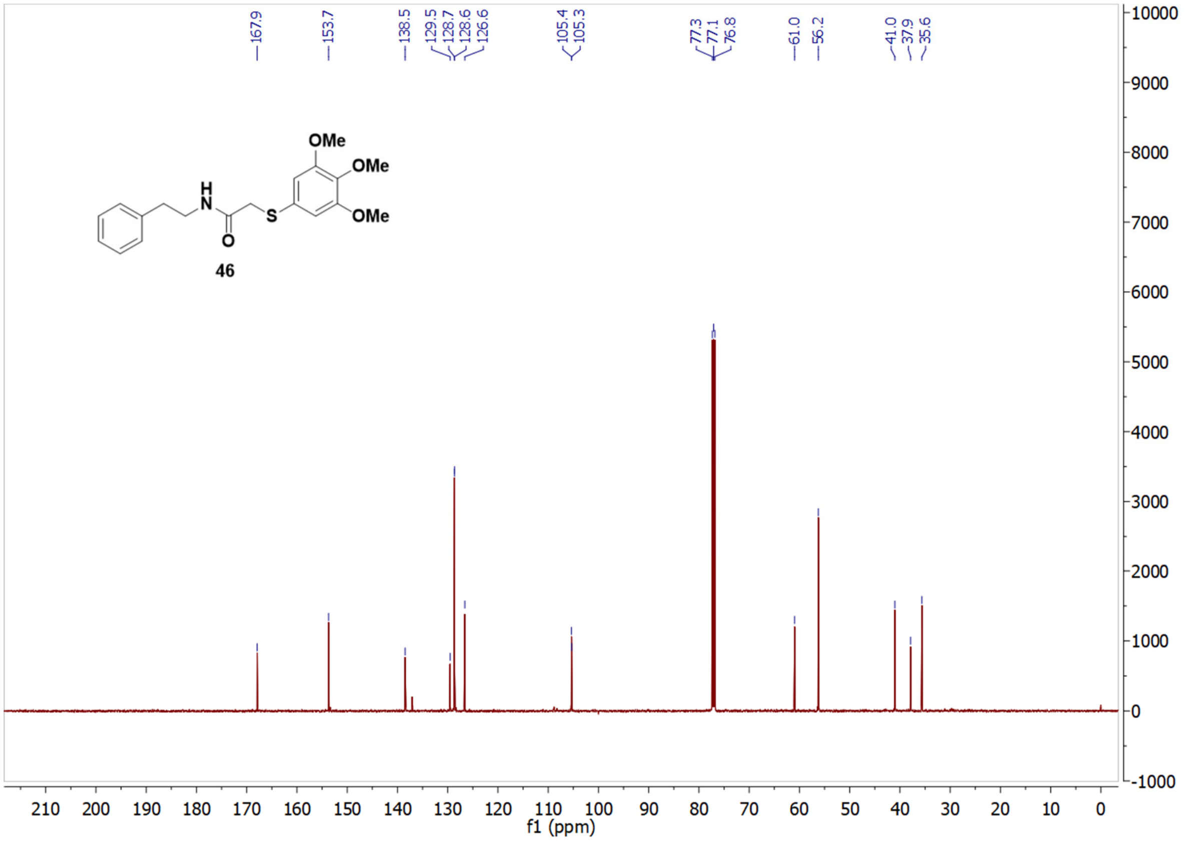

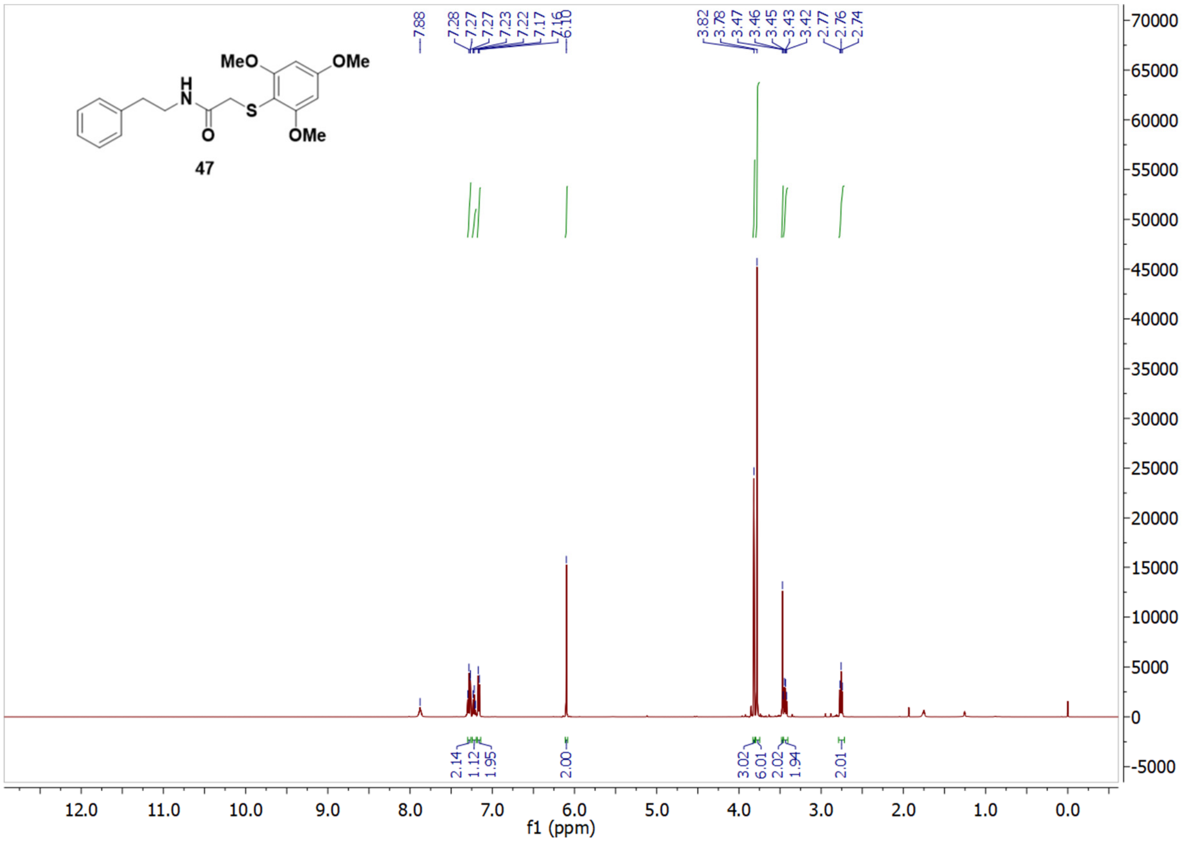
7.88
7.28
7.27
7.27
7.23
7.22
7.17
7.16
6.10
3.82
3.78
3.47
3.46
3.45
3.43
3.42
2.77
2.76
2.74
MeO
OMe
OMe
H
N
O
S
47
2.14
1.12
1.95
2.00
3.02
6.01
2.02
1.94
2.01
f1 (ppm)

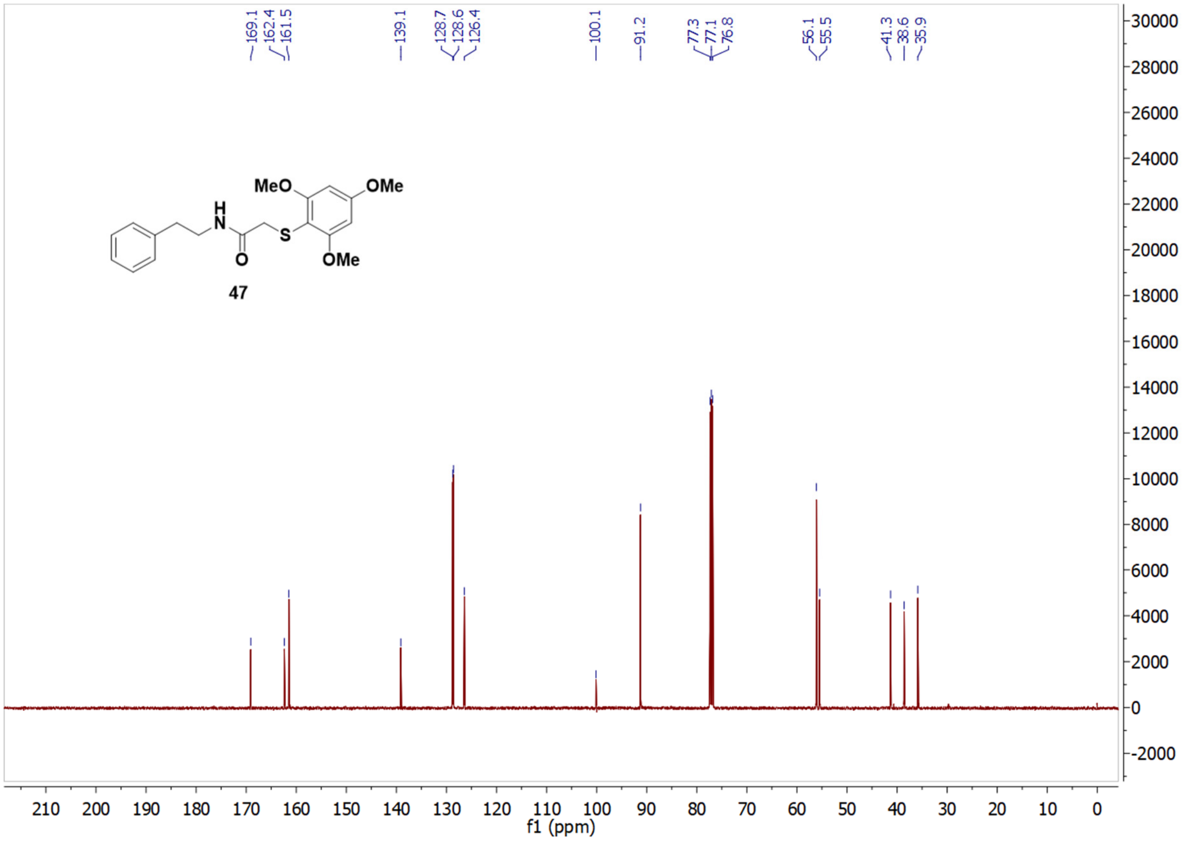
169.1
162.4
161.5
139.1
128.7
128.6
126.4
100.1
91.2
77.3
77.1
76.8
56.1
55.5
41.3
38.6
35.9
MeO
OMe
OMe
H
N
O
S
47
f1 (ppm)

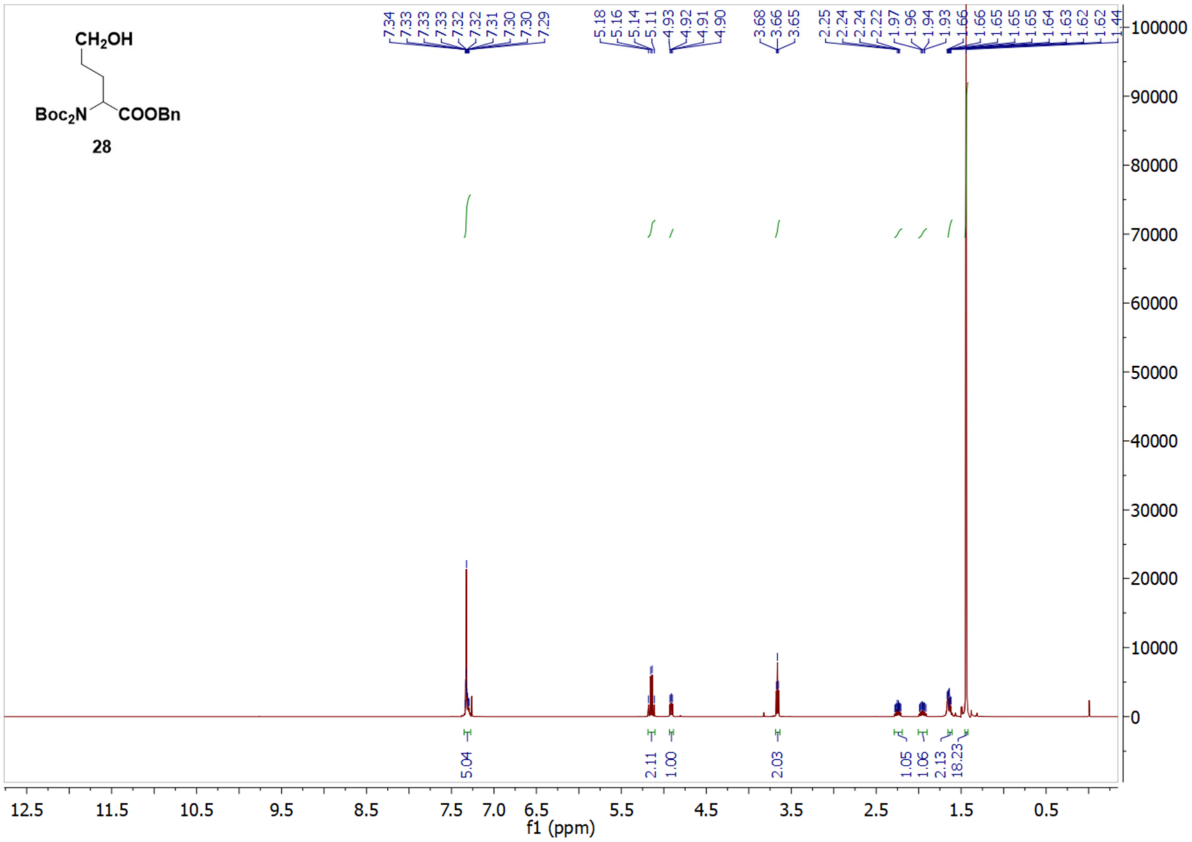
CH2OH
Boc2N
COOBn
28
7.34 7.33 7.33 7.33 7.32 7.32 7.31 7.30 7.30 7.29
5.18 5.16 5.14 5.11 4.93 4.92 4.91 4.90
3.68 3.66 3.65
2.25 2.24 2.24 2.22 1.97 1.96 1.94 1.93 1.66 1.66 1.65 1.65 1.65 1.64 1.63 1.62 1.62 1.44
5.04
2.11
1.00
2.03
1.05
1.06
2.13
18.23
f1 (ppm)

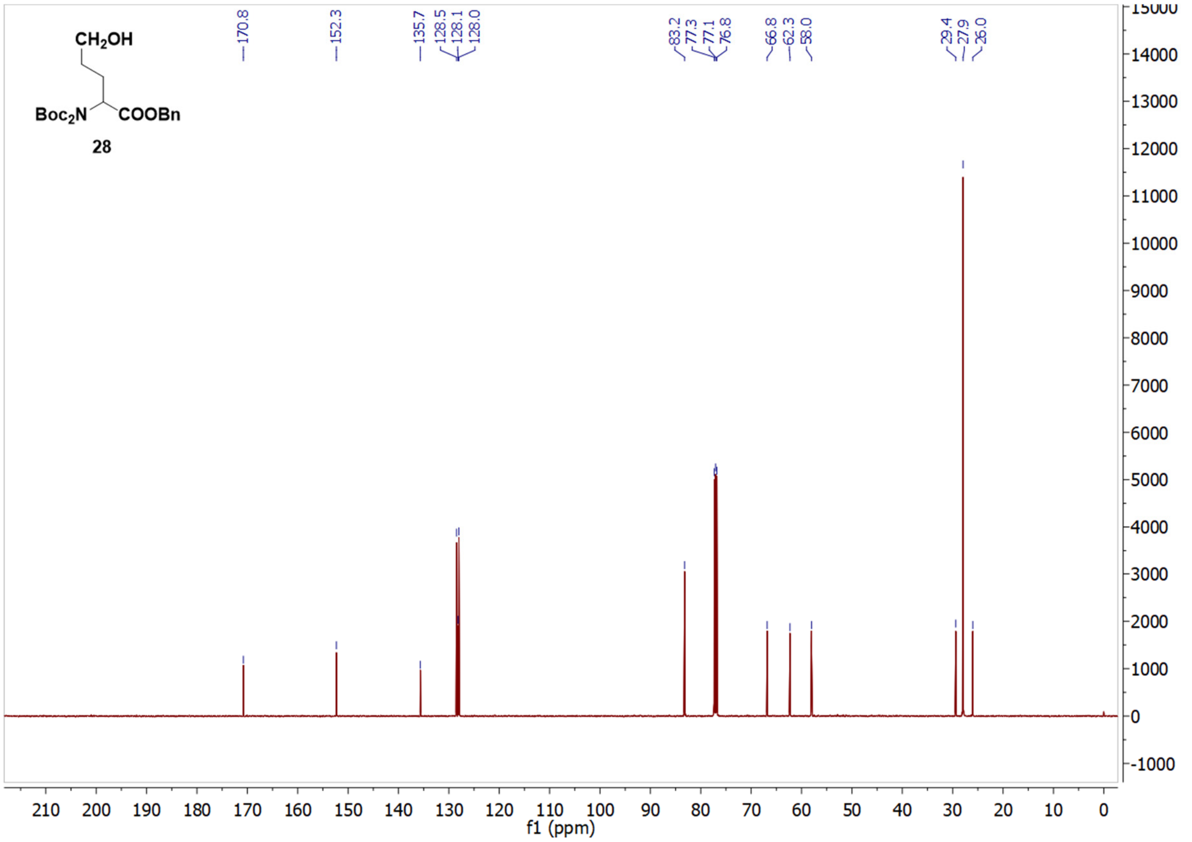
CH2OH
Boc2N
COOBn
28
170.8
152.3
135.7 128.5 128.1 128.0
83.2 77.3 77.1 76.8
66.8 62.3 58.0
29.4 27.9 26.0
f1 (ppm)

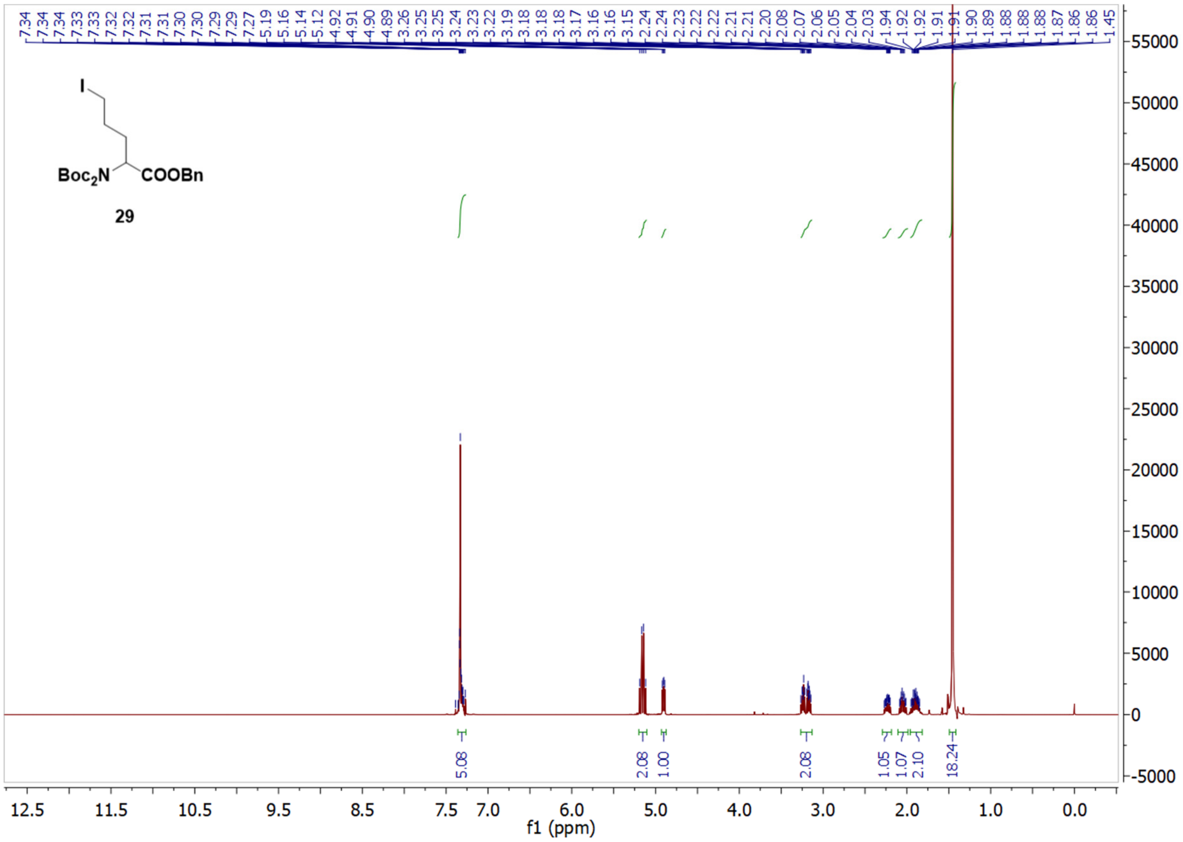

Boc$_2$N
COOBn
29
5.08
2.08
1.00
2.08
1.05
1.07
2.10
18.24
f1 (ppm)


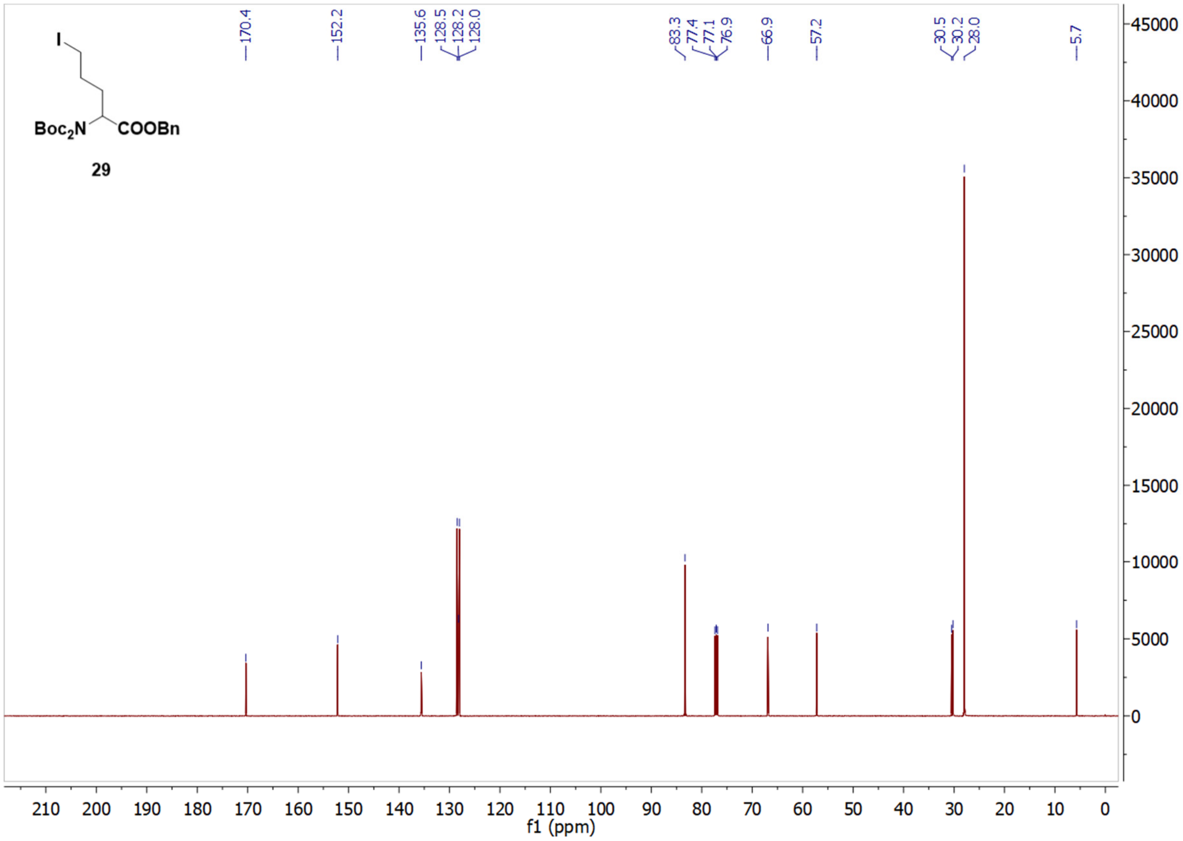

170.4
152.2
135.6
128.5
128.2
128.0
83.3
77.4
77.1
76.9
66.9
57.2
30.5
30.2
28.0
5.7
Boc$_2$N
COOBn
29
f1 (ppm)

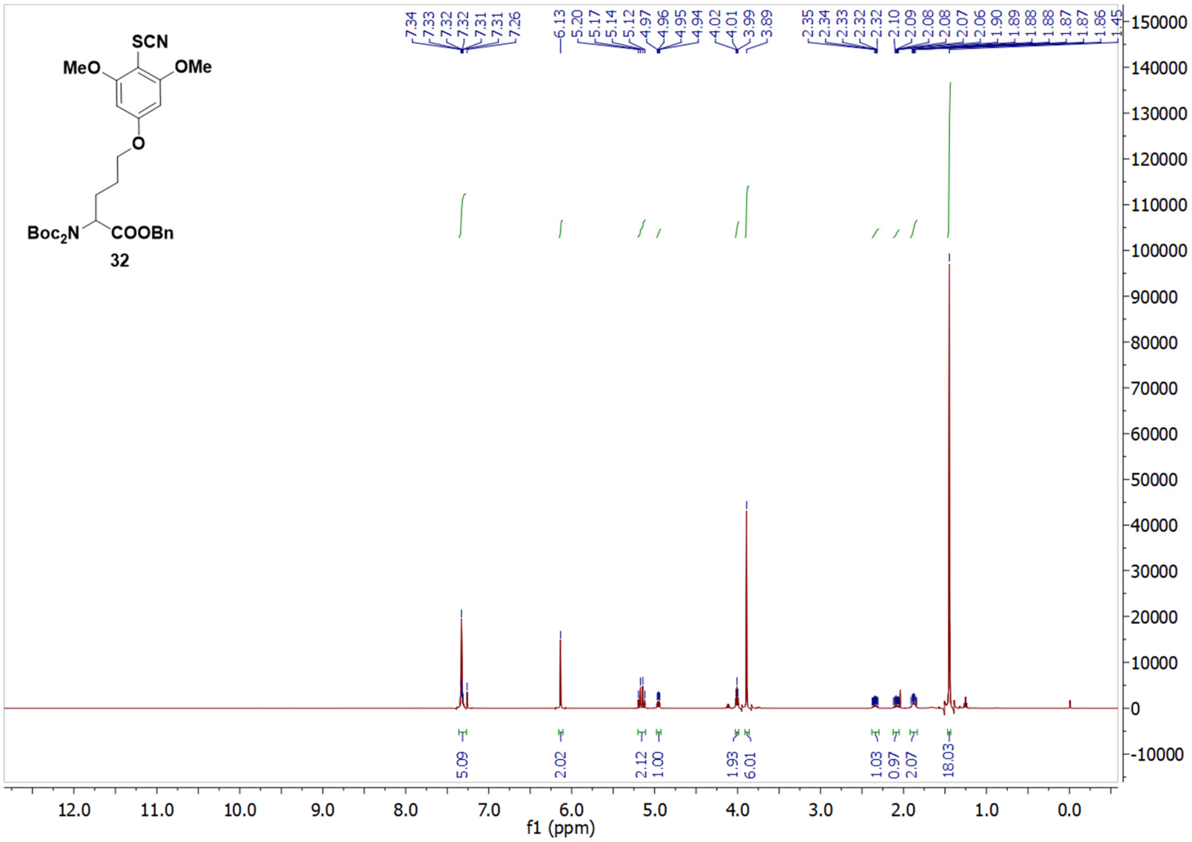
SCN
MeO
OMe
O
Boc2N
COOBn
32
7.34
7.33
7.32
7.32
7.31
7.31
7.26
6.13
5.20
5.17
5.14
5.12
4.97
4.96
4.95
4.94
4.02
4.01
3.99
3.89
2.35
2.34
2.33
2.32
2.32
2.10
2.09
2.08
2.08
2.07
2.06
1.90
1.89
1.88
1.88
1.87
1.87
1.86
1.45
5.09
2.02
2.12
1.00
1.93
6.01
1.03
0.97
2.07
18.03
f1 (ppm)

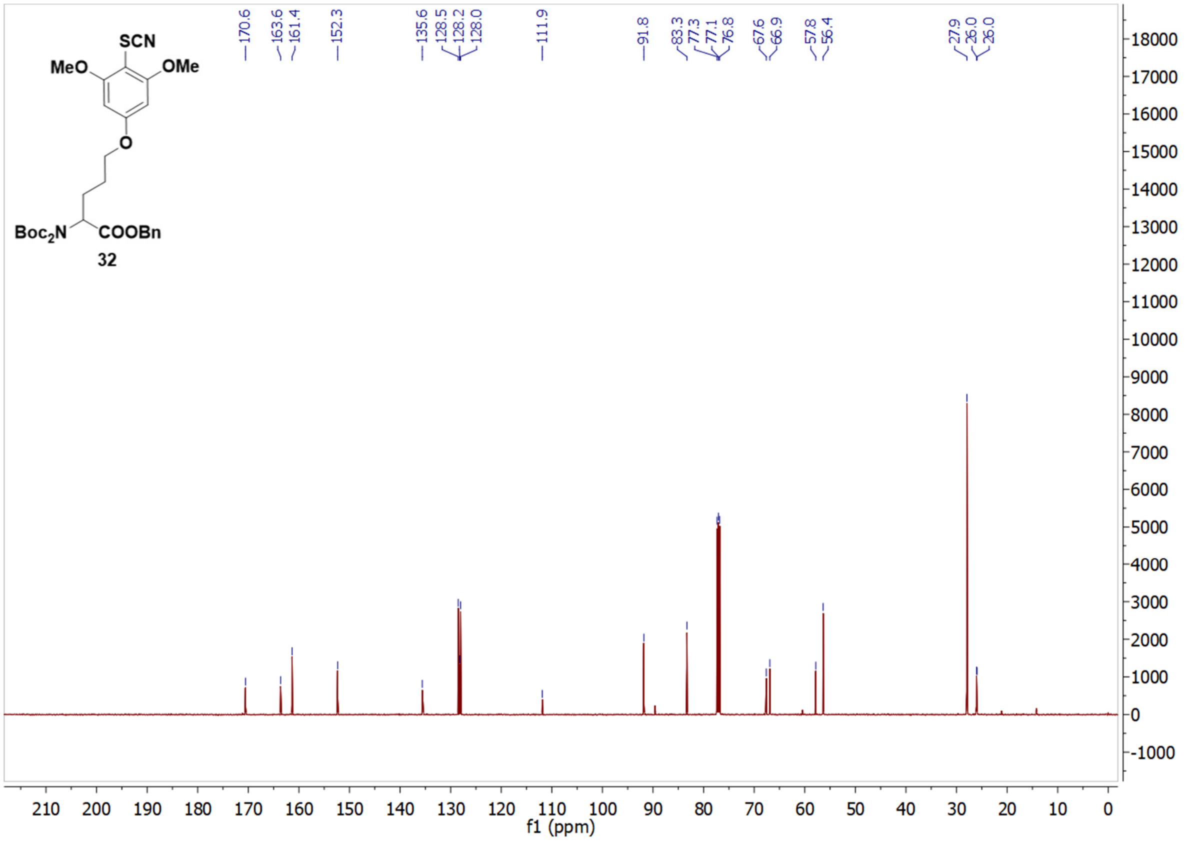
SCN
MeO
OMe
O
Boc2N
COOBn
32
170.6
163.6
161.4
152.3
135.6
128.5
128.2
128.0
111.9
91.8
83.3
77.3
77.1
76.8
67.6
66.9
57.8
56.4
27.9
26.0
26.0
f1 (ppm)

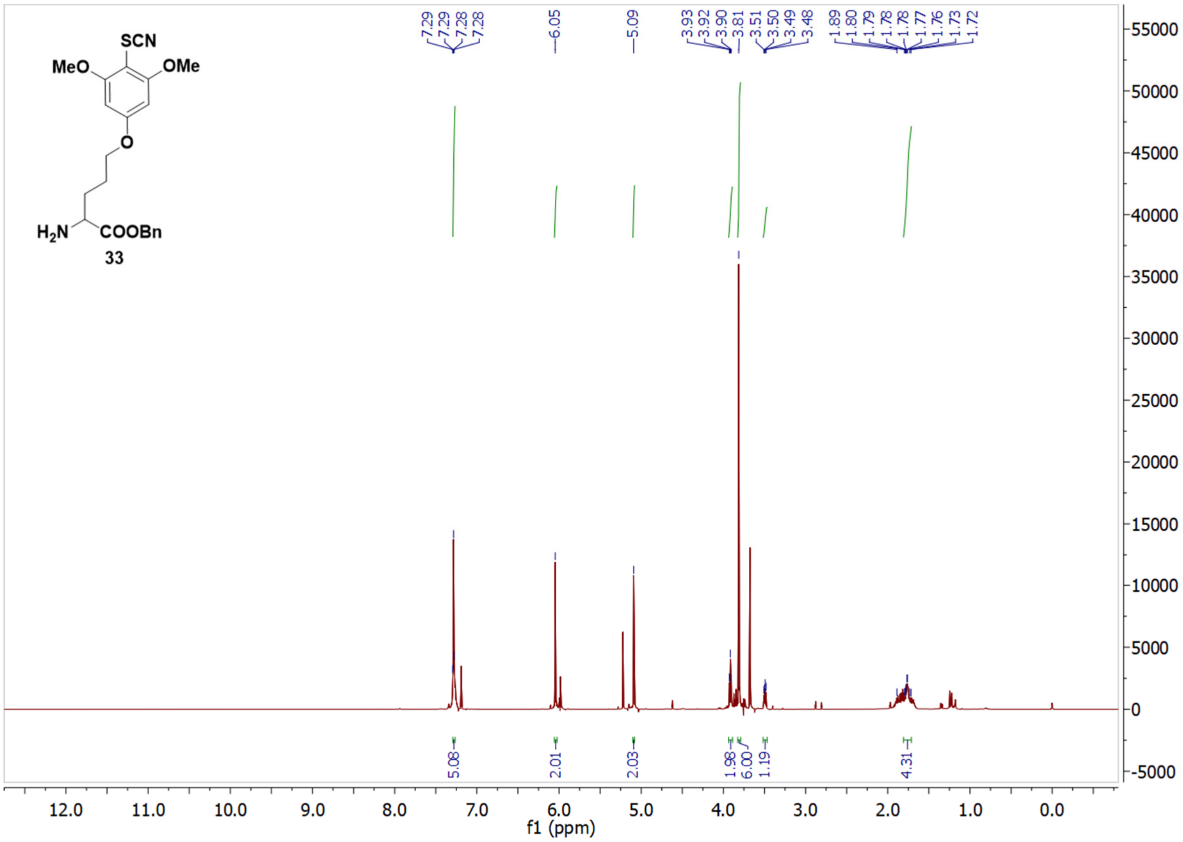

SCN
MeO
OMe
O
H2N
COOBn
33
7.29
7.29
7.28
7.28
6.05
5.09
3.93
3.92
3.90
3.81
3.51
3.50
3.49
3.48
1.89
1.80
1.79
1.78
1.78
1.77
1.76
1.73
1.72
5.08
2.01
2.03
1.98
6.00
1.19
4.31
f1 (ppm)


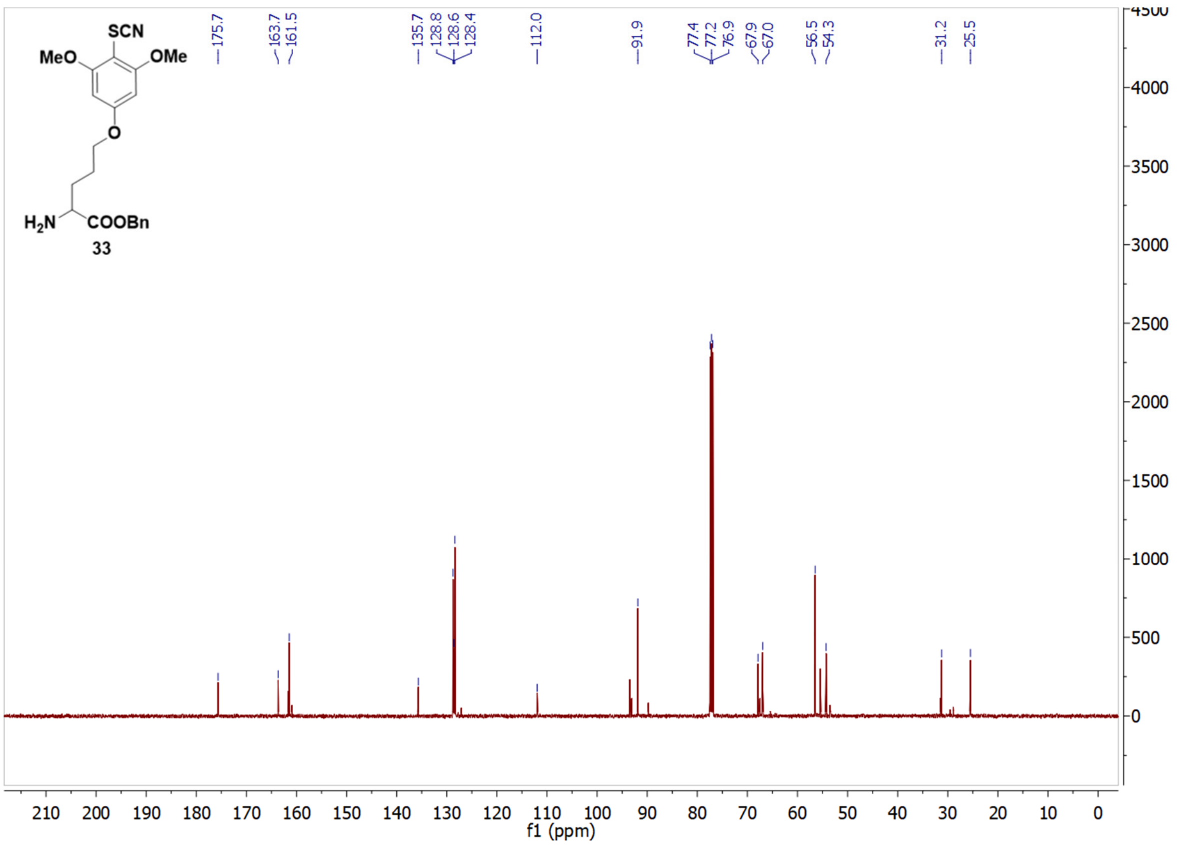

SCN
MeO
OMe
O
H2N
COOBn
33
175.7
163.7
161.5
135.7
128.8
128.6
128.4
112.0
91.9
77.4
77.2
76.9
67.9
67.0
56.5
54.3
31.2
25.5
f1 (ppm)

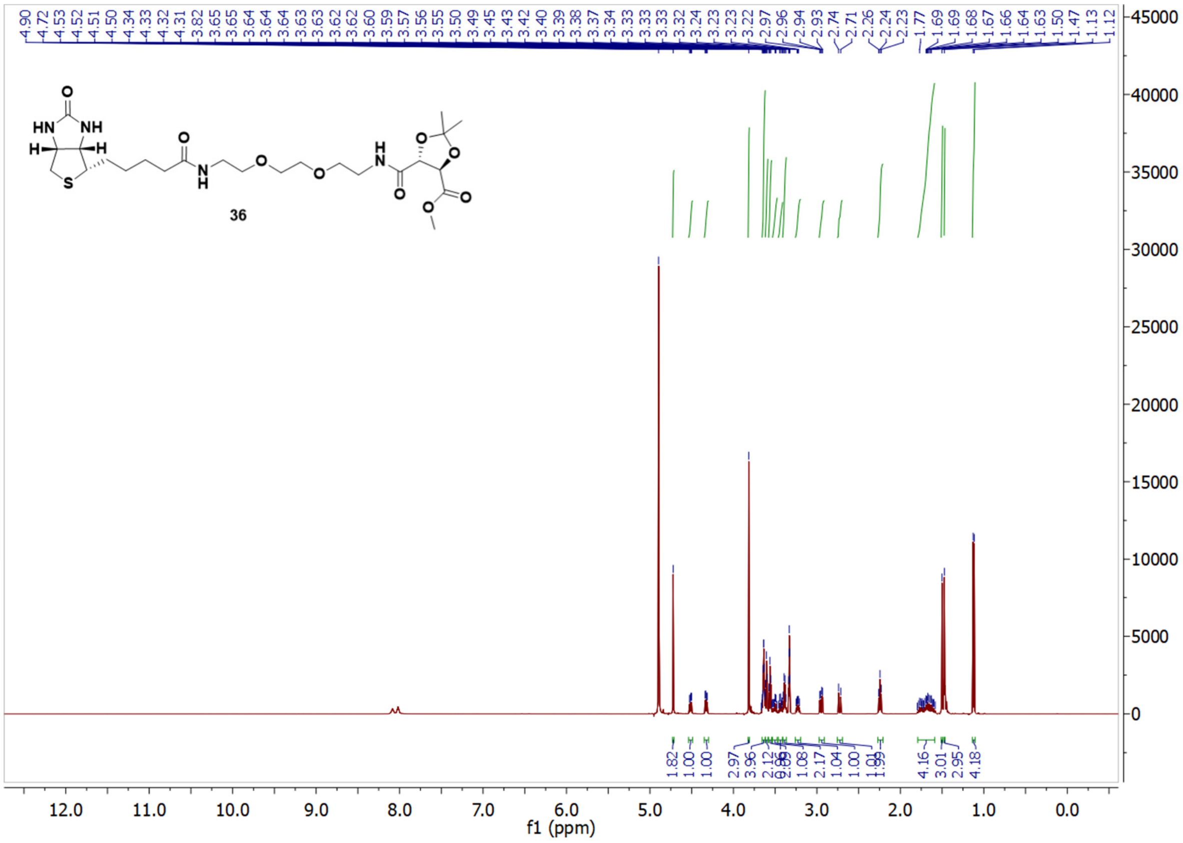
4.90 4.72 4.53 4.52 4.51 4.50 4.34 4.33 4.32 4.31 3.82 3.65 3.65 3.64 3.64 3.64 3.63 3.63 3.62 3.62 3.60 3.59 3.57 3.56 3.55 3.50 3.49 3.45 3.43 3.42 3.40 3.39 3.38 3.37 3.34 3.33 3.33 3.33 3.32 3.24 3.23 3.23 3.22 2.97 2.96 2.94 2.93 2.74 2.71 2.26 2.24 2.23 1.77 1.69 1.69 1.68 1.67 1.66 1.64 1.63 1.50 1.47 1.13 1.12
HN
NH
O
H
H
S
N
H
O
O
O
N
H
O
O
O
O
O
36
1.82 1.00 1.00 2.97 3.96 2.12 1.08 2.17 1.04 1.00 1.01 1.99 4.16 3.01 2.95 4.18
f1 (ppm)
12.0 11.0 10.0 9.0 8.0 7.0 6.0 5.0 4.0 3.0 2.0 1.0 0.0
45000 40000 35000 30000 25000 20000 15000 10000 5000 0

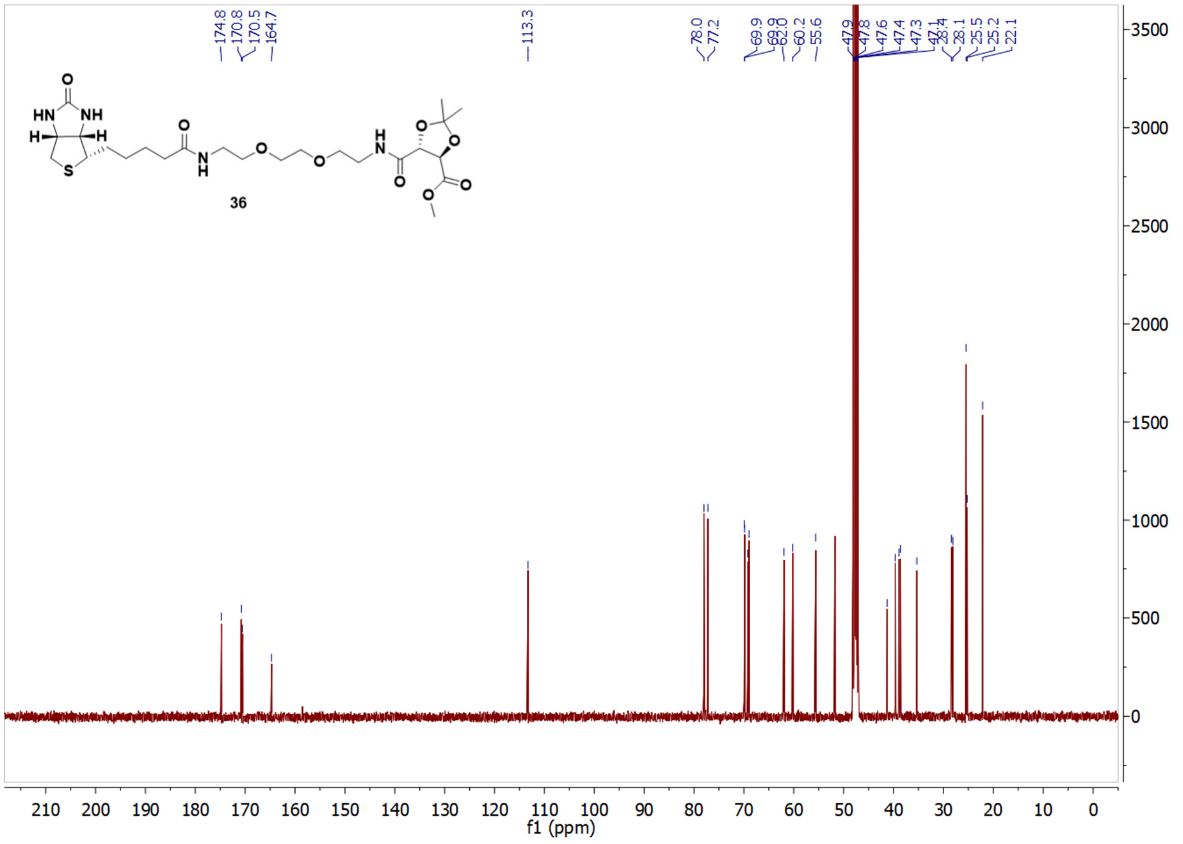
174.8 170.8 170.5 164.7 113.3 78.0 77.2 69.9 69.9 62.0 60.2 55.6 47.9 47.8 47.6 47.4 47.3 47.1 28.4 28.1 25.5 25.2 22.1
HN
NH
O
H
H
S
N
H
O
O
O
N
H
O
O
O
O
O
36
f1 (ppm)
210 200 190 180 170 160 150 140 130 120 110 100 90 80 70 60 50 40 30 20 10 0
3500 3000 2500 2000 1500 1000 500 0

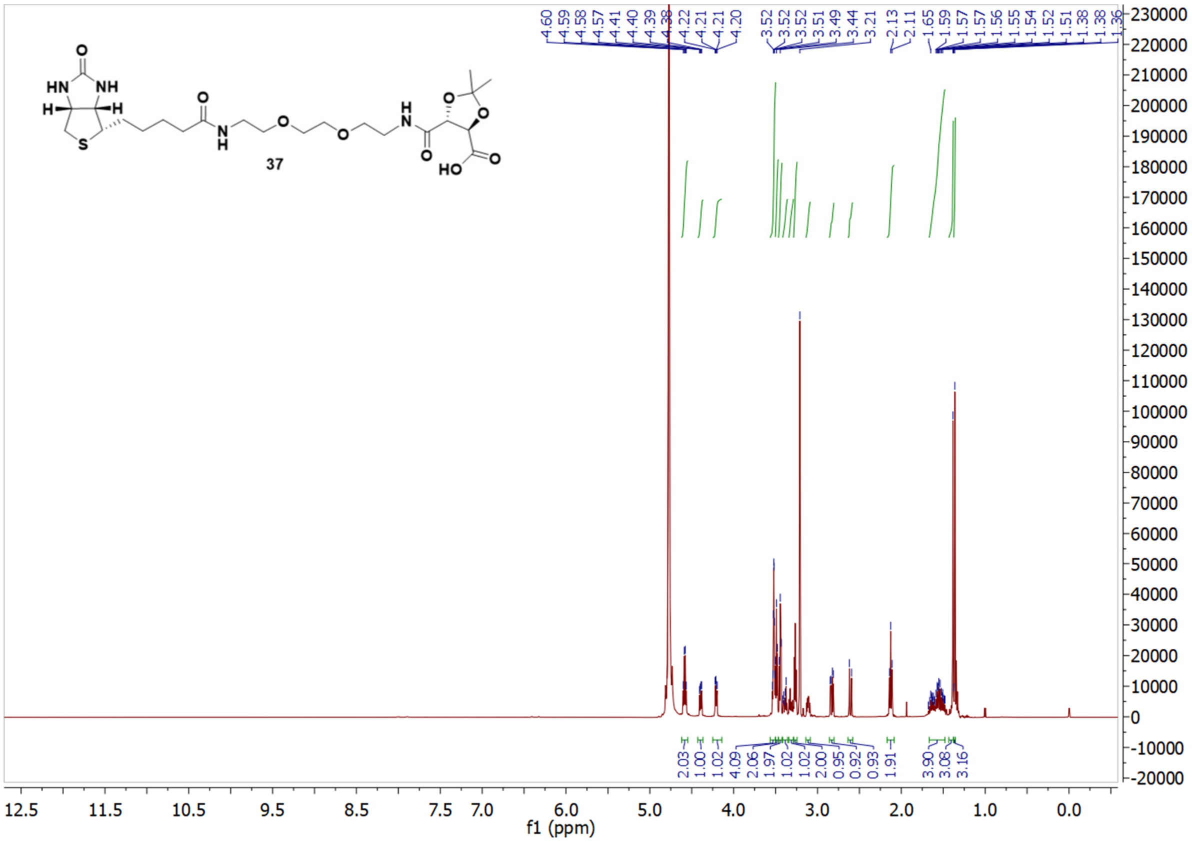

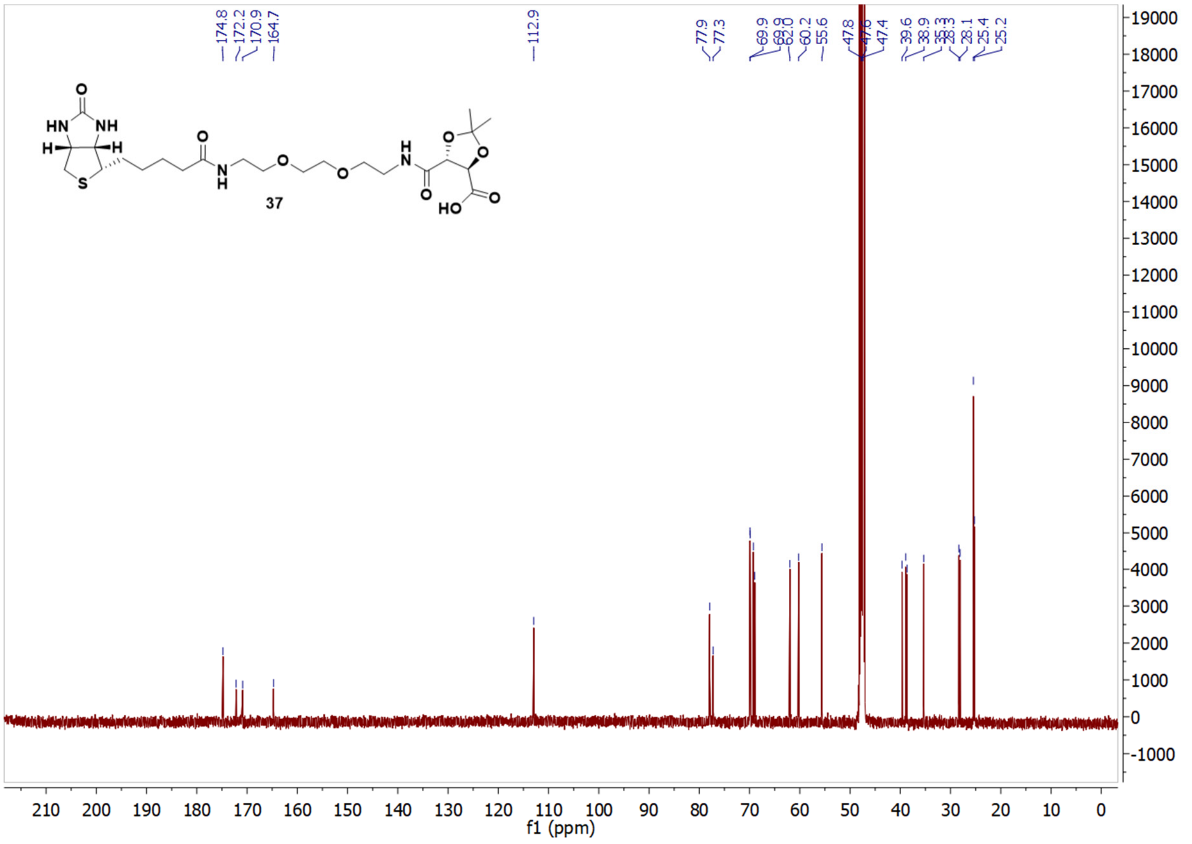

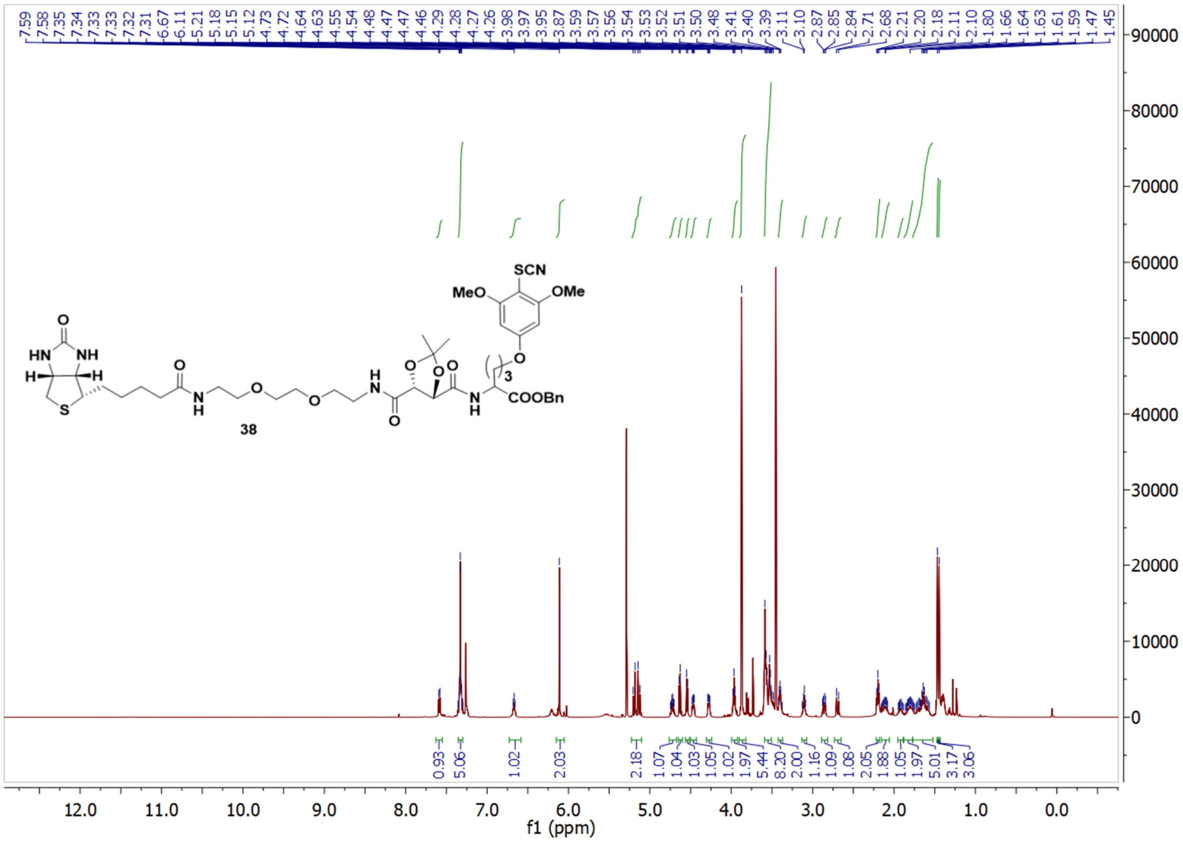

SCN
MeO
OMe
HN
NH
O
H
H
S
N
H
O
N
H
O
38
COOBn
3
12.0
11.0
10.0
9.0
8.0
7.0
6.0
5.0
4.0
3.0
2.0
1.0
0.0
f1 (ppm)


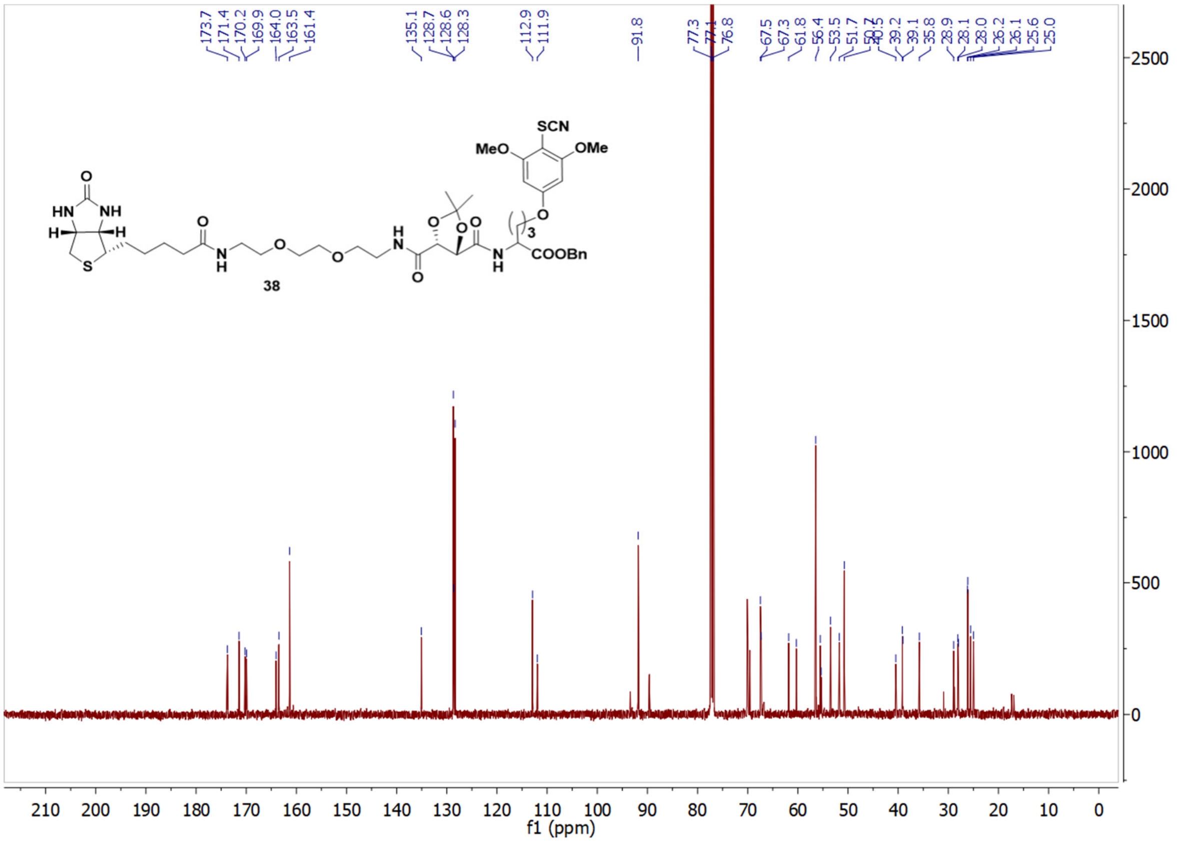

SCN
MeO
OMe
HN
NH
O
38
COOBn
3
210
200
190
180
170
160
150
140
130
120
110
100
90
80
70
60
50
40
30
20
10
0
f1 (ppm)

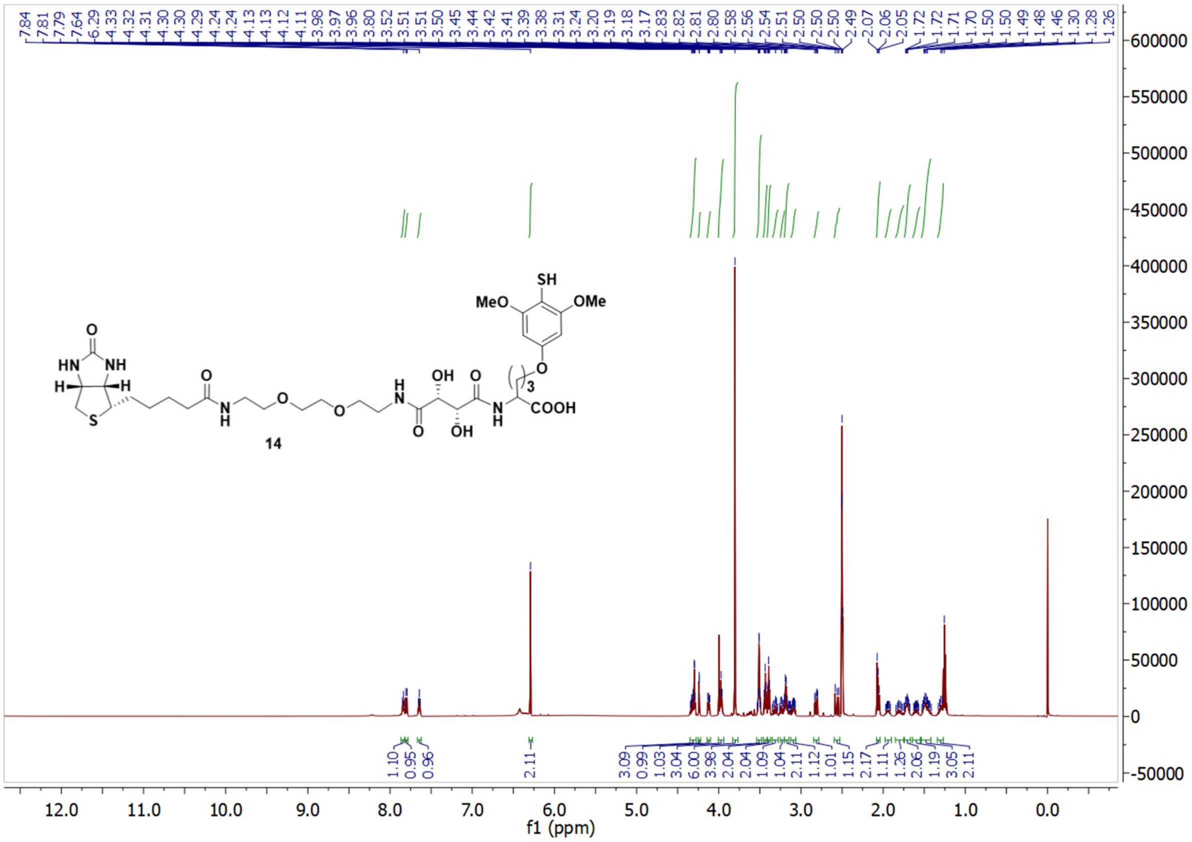
7.84 7.81 7.79 7.64 6.29 4.33 4.32 4.31 4.30 4.30 4.29 4.24 4.24 4.13 4.13 4.12 4.11 3.98 3.97 3.96 3.80 3.52 3.51 3.51 3.50 3.45 3.44 3.42 3.41 3.39 3.38 3.31 3.24 3.20 3.19 3.18 3.17 2.83 2.82 2.81 2.80 2.58 2.56 2.54 2.51 2.50 2.50 2.50 2.49 2.07 2.06 2.05 1.72 1.72 1.71 1.70 1.50 1.50 1.49 1.48 1.46 1.30 1.28 1.26
1.10 0.95 0.96 2.11 3.09 0.99 1.03 3.04 6.00 3.98 2.04 2.04 1.09 1.04 2.11 1.12 1.01 1.15 2.17 1.11 1.26 2.06 1.19 3.05 2.11
SH
MeO
OMe
O
HN
NH
H
H
S
O
N
H
O
O
N
H
OH
O
O
OH
N
H
3
COOH
14
f1 (ppm)

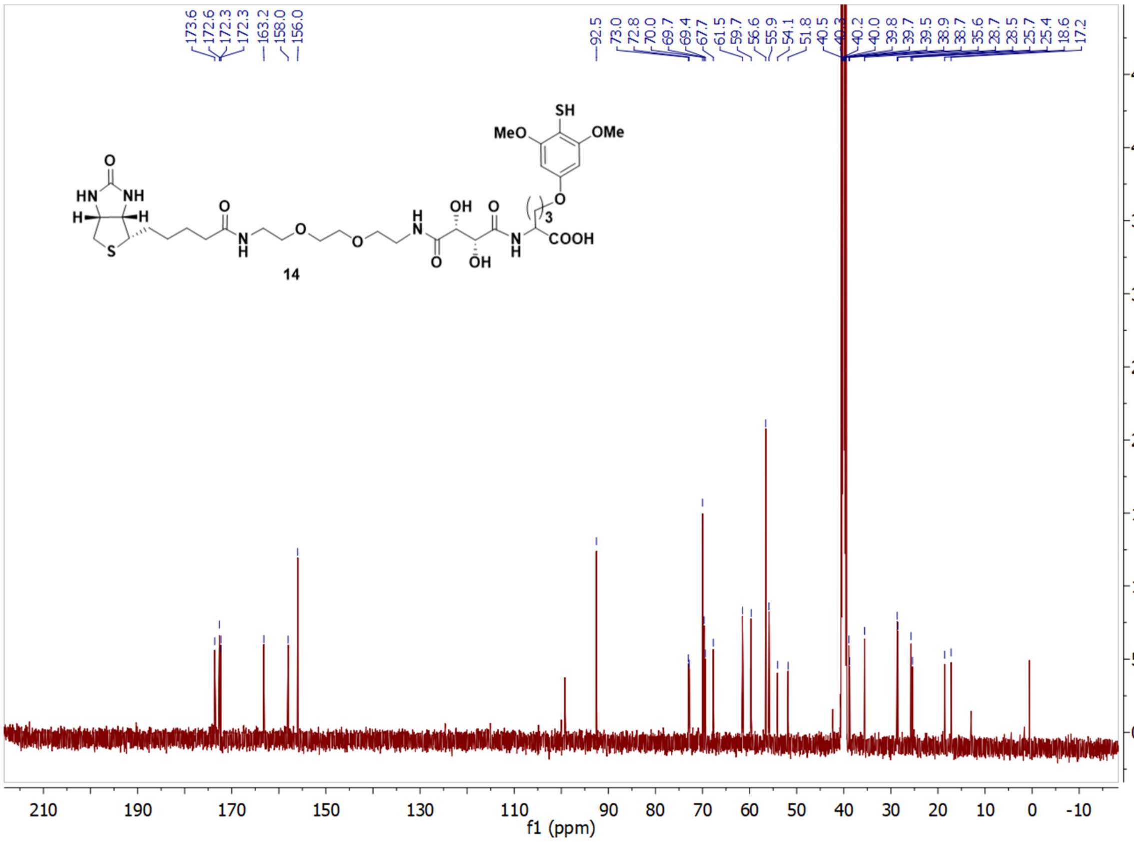
173.6 172.6 172.3 172.3 163.2 158.0 156.0 92.5 73.0 72.8 70.0 69.7 69.4 67.7 61.5 59.7 56.6 55.9 54.1 51.8 40.5 40.3 40.2 40.0 39.8 39.7 39.5 38.9 38.7 35.6 28.7 28.5 25.7 25.4 18.6 17.2
SH
MeO
OMe
O
HN
NH
H
H
S
O
N
H
O
O
N
H
OH
O
O
OH
N
H
3
COOH
14
f1 (ppm)

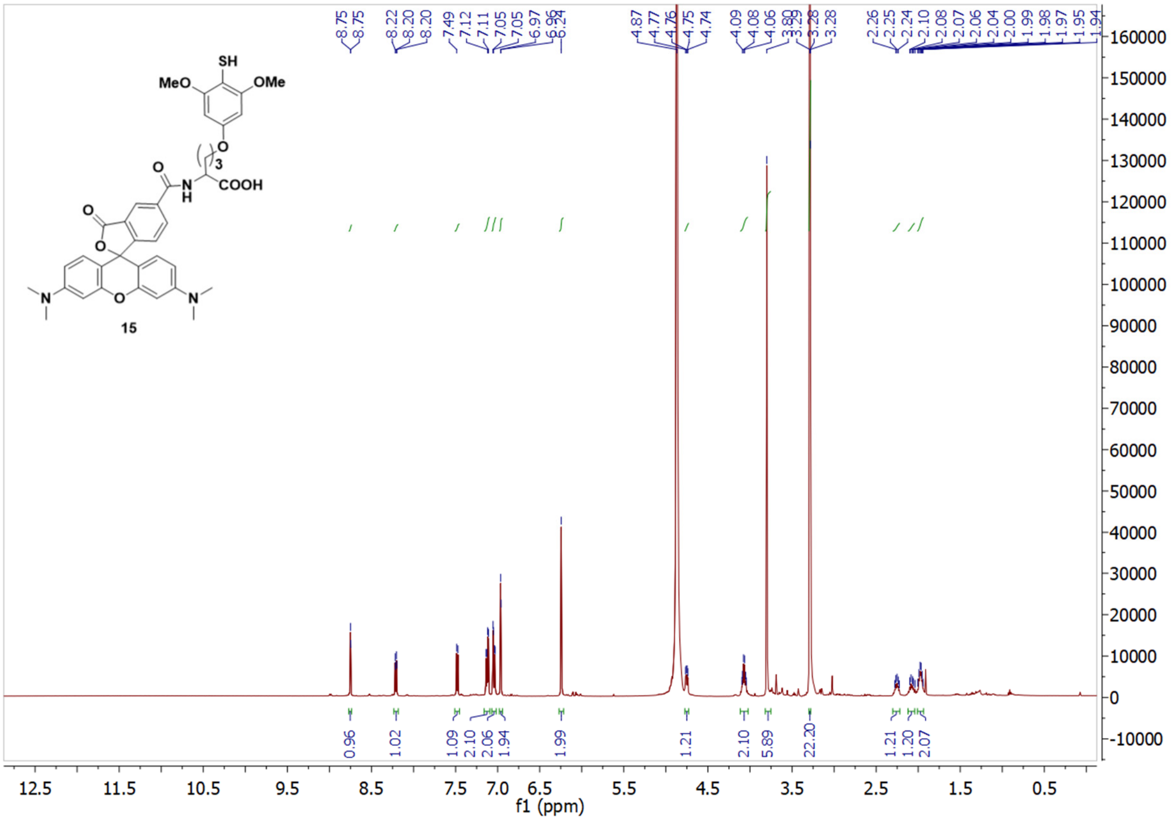

SH
MeO
OMe
O
3
O
N
H
COOH
O
O
N
O
N
15
f1 (ppm)


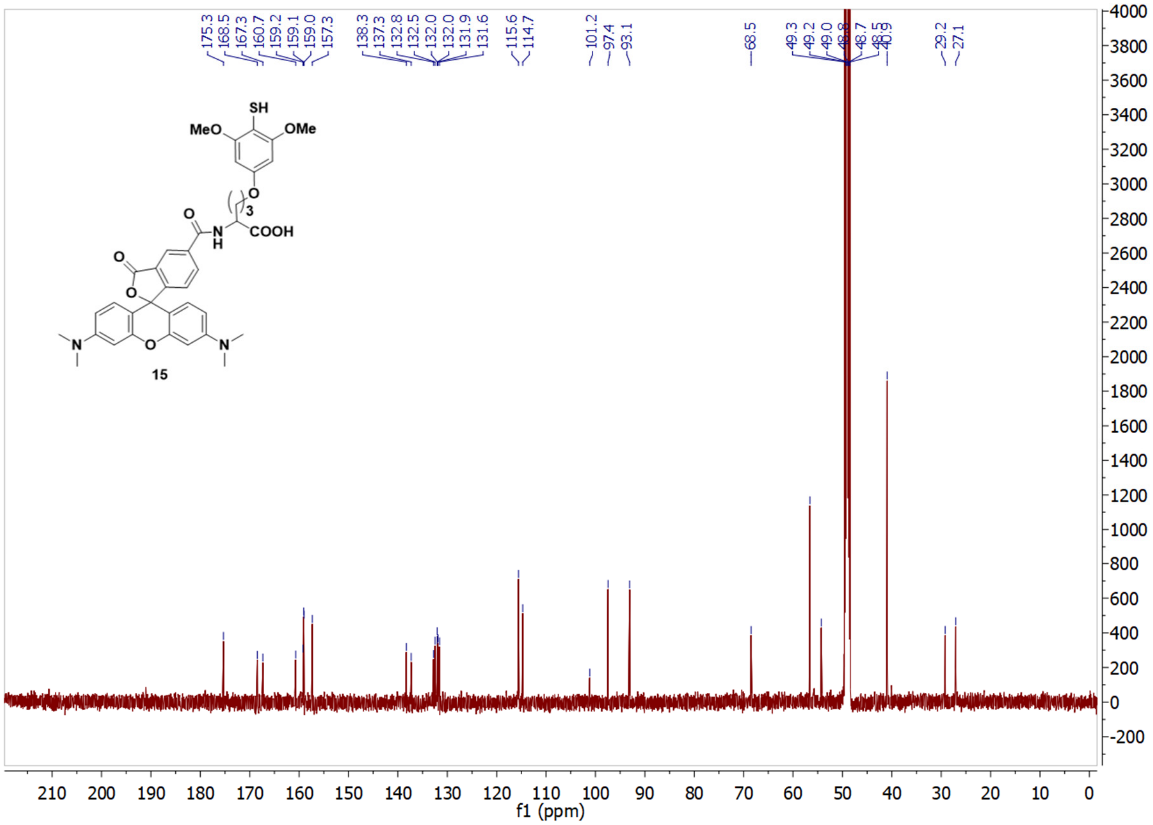

SH
MeO
OMe
O
3
O
N
H
COOH
O
O
N
O
N
15
f1 (ppm)

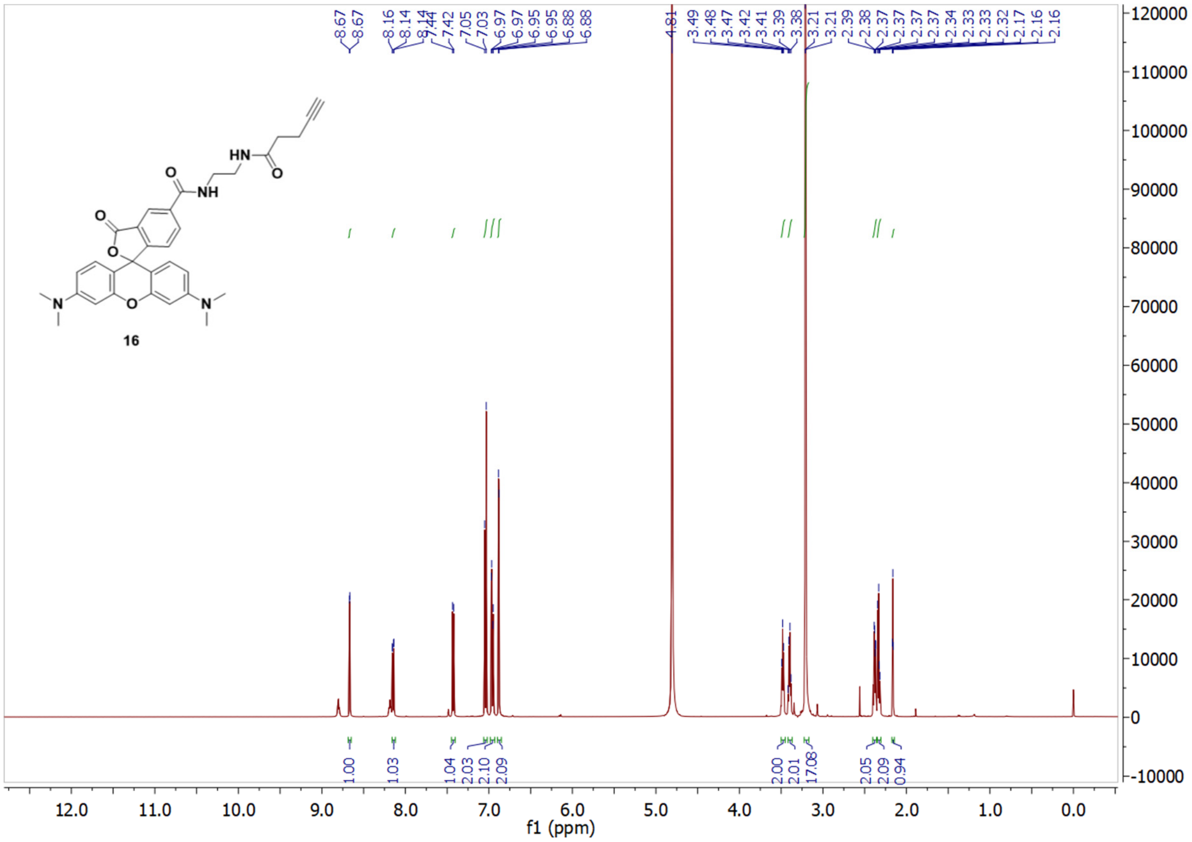
8.67
8.67
8.16
8.14
8.14
7.44
7.42
7.05
7.03
6.97
6.97
6.95
6.95
6.88
6.88
4.81
3.49
3.48
3.47
3.42
3.41
3.39
3.38
3.21
3.21
2.39
2.38
2.37
2.37
2.37
2.37
2.34
2.33
2.33
2.32
2.17
2.16
2.16
HN
O
NH
O
O
O
N
O
N
16
1.00
1.03
1.04
2.03
2.10
2.09
2.00
2.01
17.08
2.05
2.09
0.94
f1 (ppm)

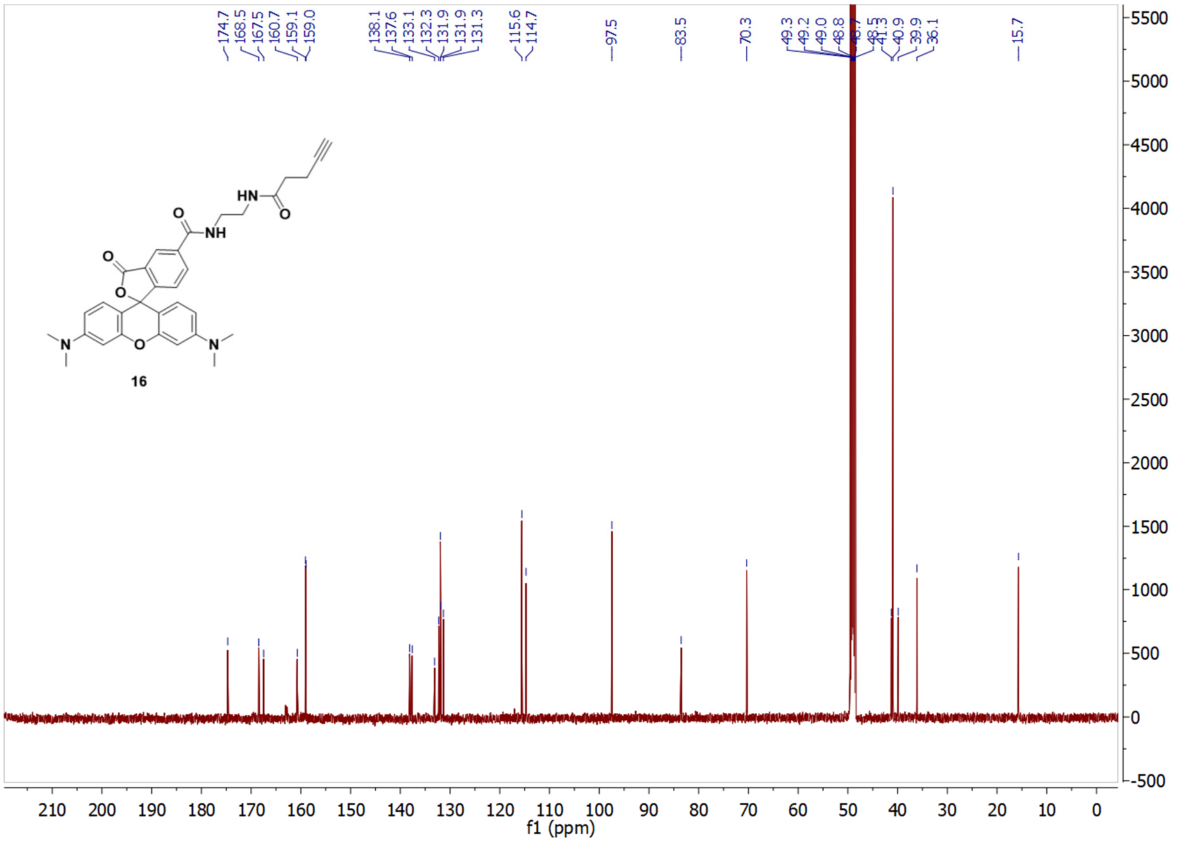
174.7
168.5
167.5
160.7
159.1
159.0
138.1
137.6
133.1
132.3
131.9
131.9
131.3
115.6
114.7
97.5
83.5
70.3
49.3
49.2
49.0
48.8
48.7
48.5
41.3
40.9
39.9
36.1
15.7
HN
O
NH
O
O
O
N
O
N
16
f1 (ppm)
